# Immune-cancer analyses across mammals reveal a potential trophic level and platelet-linked tradeoff between cancer and trauma mortality

**DOI:** 10.64898/2025.12.09.693265

**Authors:** Stefania E. Kapsetaki, Sareh Seyedi, Zachary T. Compton, Shawn M. Rupp, Elizabeth G. Duke, Joshua D. Schiffman, Brigid V. Troan, Tara M. Harrison, Carlo C. Maley, Lisa M. Abegglen, Amy M. Boddy

**Author notes:** co-senior authors (Carlo C. Maley, Lisa M. Abegglen, Amy M. Boddy).

## Abstract

There may be fitness tradeoffs between wound healing, immune responses, and cancer development due to shared pathways, limited resources and conflicting selective pressures. The immune system is important in both response to injury and carcinogenesis. We initially investigated correlations between cancer prevalence and immune cells, controlling for known associations with body mass and lifespan. We analyzed data from 216 mammalian species from at least 20 individuals per species. Body mass correlated positively with segmented neutrophil-to-lymphocyte ratios and negatively with lymphocyte concentrations. However, only platelet concentration correlated (negatively) with cancer prevalence (*P*-value = 0.006). To further understand this association, we investigated whether a fitness tradeoff could exist between preventing death from cancer versus injury. We discovered a negative correlation between cancer and trauma mortalities (*P-*value ≤ 0.0006), even when we accounted for the fact that different causes of death must sum to 100%. Platelet size and trophic level negatively correlated with trauma mortality, but not when controlling for cancer mortality (*P-*value = 0.06). If trauma mortality is an indirect measure of wound healing, this suggests a fitness tradeoff may exist between cancer suppression and wound healing across mammals, mediated in part through platelet size and trophic level.

## Introduction

A fundamental question in comparative oncology is why some species are more susceptible to cancer than others. We do not yet understand what evolutionary tradeoffs constrain cancer suppression, nor the mechanisms that determine cancer susceptibility. Despite a number of recent comparative oncology studies, only a small proportion of the variance in cancer prevalence or risk has been explained by factors such as body mass ^1,2^, gestation time^2^, litter/clutch size^3,4^, germline mutation rate^5^, and carnivory^6^. Selection for diversity in the MHC class I gene complex explains approximately 37% of the variation in cancer risk across 28 mammalian species^7^. Given the central role the immune system plays in both cancer promotion (through chronic inflammation) and cancer surveillance, we first hypothesized that a relationship may exist between immune system components and cancer prevalence across species. We found a relationship between platelet concentration and cancer prevalence reminiscent of a previous hypothesis for a fitness tradeoff between wound healing and cancer suppression, based on common mechanisms that drive cancer and wound healing^8–11^.

Currently, little is known about tradeoffs between immune system function, wound healing and cancer suppression. In humans, the immune system plays an important role in wound healing in response to physical trauma^12–14^, as well as in identifying and eliminating tumor cells^15,16^. Tumors often co-opt the wound healing response, which led to their characterization as “wounds that do not heal.”^17,18^

Although the effectiveness of each species’ immune system in suppressing cancer has not yet been quantified across species, studies have tested for associations between body mass and various immune cell concentrations. Larger birds tend to have higher concentrations of white blood cells^19^ and lymphocytes^19,20^ than smaller birds. Larger mammals tend to have higher concentrations of neutrophils (functionally equivalent to heterophils)^21,22^, white blood cells^22,23^, (and monocytes in felids^24^) but lower concentrations of lymphocytes compared to smaller mammals^21,22,24^. Evidence on the relationship between body mass and cancer prevalence across species conflicts^1,3,4,6,25^, but a recent study suggests that larger animals tend to get slightly more cancer than smaller animals, particularly when the protective effects of gestation time is taken into account^2^.

Apart from the abovementioned immune cells, platelets may also explain some of the variation in cancer prevalence and risk across species. Platelets play a dual role in cancer; sometimes leading to tumor inhibition^26^ and other times to tumor growth and metastasis^26–29^. Platelets increase in size when they are activated, and larger platelets are associated with worse outcomes for patients with pancreatic ductal adenocarcinoma^30^. Platelets are also involved in blood hemostasis, inflammation, and clot formation^31^, which can stop bleeding after injury and decrease trauma mortality. Although functional studies on the role of platelets across species are lacking, the dual role of platelets in cancer and wound healing in studies of a few species (dogs^32–34^, mice^26,35,36^, rats^37,38^, humans^30,39–41^) is a hint that there may be a fitness tradeoff between cancer suppression and wound healing/prevention of death from injury across mammals.

This potential tradeoff may vary with trophic level, i.e., the position of an organism in the food chain. This position affects the evolution of its physiology, which may include its immune system and vulnerability to different causes of mortality. Both carnivores and herbivores get injured from intraspecific conflict and predator-prey encounters. For example, carnivores and herbivores sustain traumatic injuries from accidents, predation or interspecies aggression, parental aggression, self-inflicted wounds, or during breeding-related interactions^42,43^. Veterinary observation suggests that wounds heal faster in carnivores than herbivores (Tara Harrison, DVM, pers. comm.), which may improve their protection from trauma mortality. Carnivores may have been under more selective pressure to heal wounds than herbivores. Initiating pathways of wound healing may mean initiating pathways of cancer development, given that wound healing and cancer have common hallmarks^10^. So this might explain why carnivores get more cancer^6^ and neoplasia (when controlling for domestication)^44^ than herbivores. While the functional aspects of cancer and its connections to wound healing, trophic levels, and platelets remain largely unexplored across species, a study found that two carnivorous species—cats and dogs—exhibit larger normal baseline platelet sizes compared to other herbivorous species, including horses, cows, and goats^45^.

We hypothesized that a fitness tradeoff exists between cancer suppression and wound healing/preventing death from injury mediated, in part, by platelets, and trophic levels. In the absence of data on wound healing rates, we used data on trauma mortality as a proxy to test our hypothesis.

To test for a tradeoff between cancer mortality and trauma mortality, we used a phylogenetic regression, using an additive log-ratio transformation of the mortality data to control for the fact that all sources of mortality must sum to 100% (compositionality). To investigate whether immune cell components mediated that relationship we used phylogenetic regressions to test immune cell type concentrations as predictors of cancer prevalence, cancer mortality or trauma mortality. Specifically, we tested platelet size and concentration, white blood cell concentrations, lymphocyte concentrations, monocyte concentrations, segmented neutrophil concentrations, and the segmented neutrophil-to-lymphocyte ratio, as predictors, and controlled for trophic level, body mass, and/or maximum longevity across up to 216 mammalian species.

## Results

To determine if a relationship could exist between immune cell type concentrations, platelet size, and cancer prevalence/mortality, we performed phylogenetic regressions between body mass, maximum lifespan, immune cells, cancer prevalence, and cancer mortality. Body mass positively correlated with the segmented neutrophil-to-lymphocyte ratio (n = 72 species; PGLS, *P*-value < 0.00001, regression coefficient = 1.03) and negatively correlated with lymphocyte concentration (cells per liter) in whole blood (n = 62 species; PGLS, *P*-value = 0.001, regression coefficient = –0.65), when controlling both analyses for species’ maximum lifespan (supp. Fig.s 3A, 4A; Table 1). No other immune or blood cell parameter (i.e., overall concentration of white blood cells, segmented neutrophils, monocytes, median platelet size, and median platelet concentration in whole blood) correlated with body mass or lifespan, across mammalian species after corrections for multiple testing (supp. Fig.s 1, 2, 3B, 4B, 5, 6, 7; Table 1).

**Figure 1.**
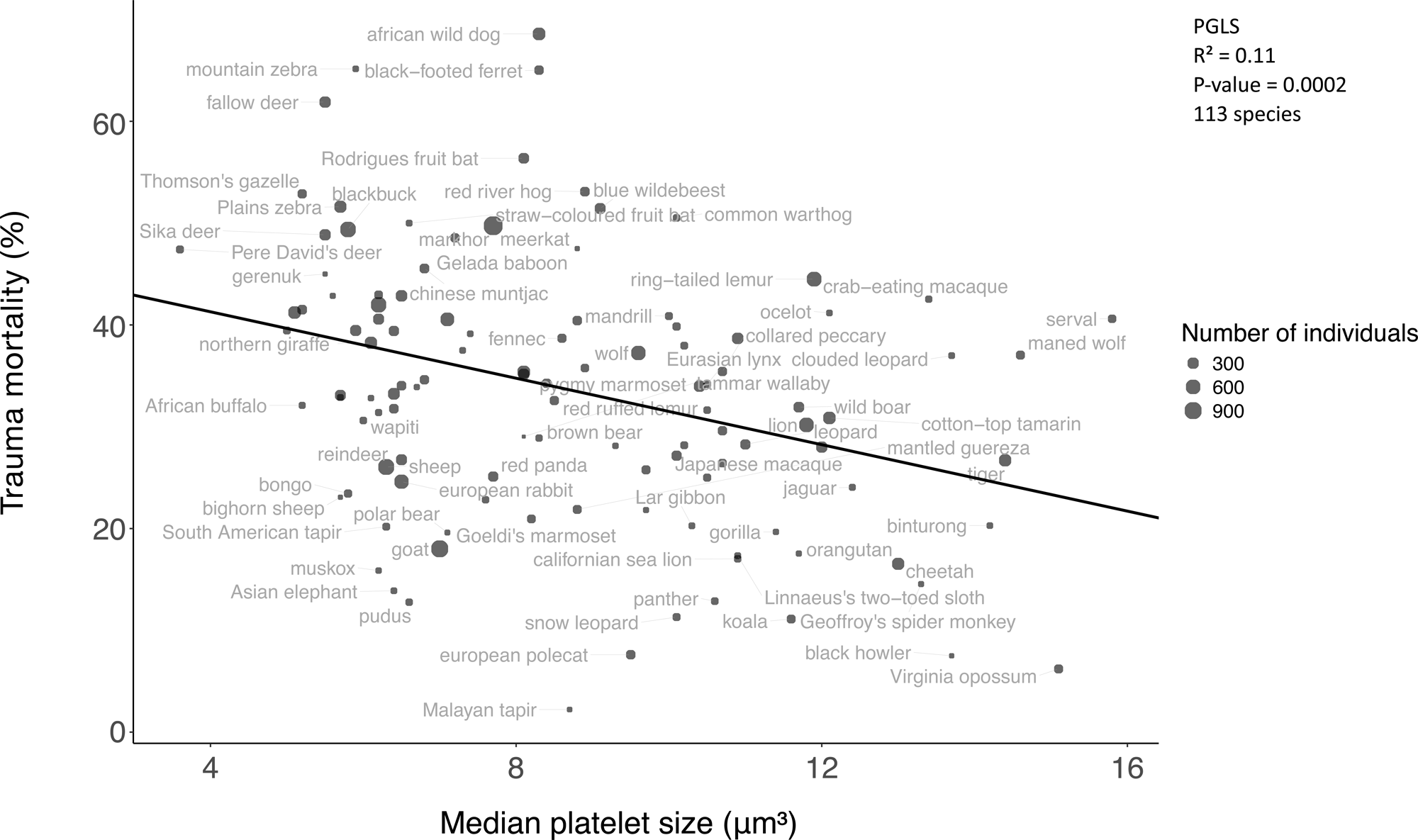
Trauma mortality is negatively correlated with median platelet size. Each dot shows the trauma mortality and median platelet size of one species. The regression line shows the phylogenetically-controlled linear regression between the trauma mortality and median platelet size.

Concentrations of the majority of tested immune cells (concentration of white blood cells, lymphocytes, segmented neutrophils, platelets, monocytes, segmented neutrophil-to-lymphocyte ratio) and platelet size did not correlate with cancer prevalence or cancer mortality (Fig. 2; Supp. Fig.s 8-12, 13A, B, E; Table 1). Only platelet concentration correlated (negatively) with cancer prevalence (n = 73 species; PGLS, *P*-value = 0.006, regression coefficient = –0.60; Fig. 2). This correlation remained when controlling the analysis for trophic level alone or all the other factors that had previously been associated with neoplasia or cancer prevalence: trophic level, body mass, and gestation length (Table 1).

**Figure 2.**
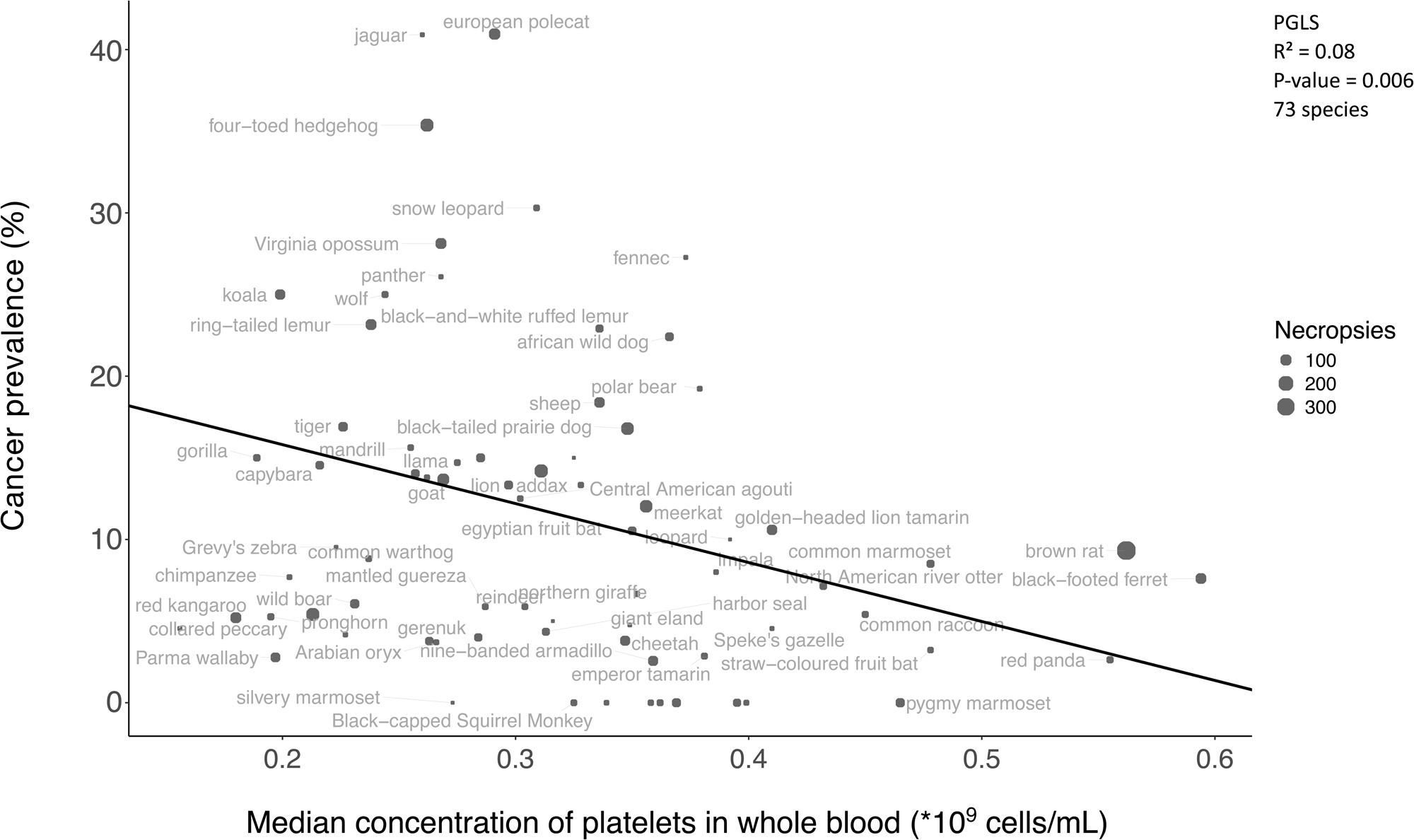
Cancer prevalence is negatively correlated with median platelet concentration. Each dot shows the cancer prevalence and median platelet concentration of one species. The regression line shows the phylogenetically-controlled linear regression between cancer prevalence and median platelet concentration.

Because measuring platelets in Felidae can be technically challenging due to their tendency to aggregate more than platelets of other species^46^, we also excluded Felidae from our platelet concentration analysis but still found that platelet concentration remained negatively correlated with cancer prevalence (PGLS, *P*-value = 0.01, regression coefficient = –0.57; Supp. Fig. 16). Neither cancer prevalence nor cancer mortality correlated with body mass in univariate analyses, however, consistent with prior work^2,4,6^, after adjusting for gestation length, body mass was a significant predictor of cancer prevalence but not of cancer mortality (Table 1).

This finding of a negative correlation between platelet concentration and cancer prevalence, and higher concentration of innate immune cells in larger species, led us to test whether there may be a trade-off between wound healing/preventing death from injury and cancer defenses mediated by platelets, the segmented neutrophil-to-lymphocyte ratio, and trophic levels. After performing phylogenetic regressions between trauma mortality, cancer prevalence, cancer mortality, platelet concentrations, platelet size, the segmented neutrophil-to-lymphocyte ratio, and trophic levels (Fig. 1; Supp. Fig. 13C; Table 1), we found the following partly supporting our hypothesis. Trauma mortality negatively correlated with cancer prevalence (n = 78 species; PGLS, *P*-value = 0.01, regression coefficient = –0.26; Fig. 4; Table 1) and cancer mortality (n = 200 species; PGLS, *P*-value ≤ 0.0006, regression coefficient < 0; Fig. 3; Table 1; even when removing the 8 statistically significant outlier species with the highest cancer mortality from the analysis: PGLS, *P-*value = 0.003, lambda = 0.64, R² = 0.08, regression coefficient = –0.35). Trauma and cancer mortality may be negatively correlated due to the fact that they have the same denominator (if the proportion of deaths due to one cause increases, the sum of the proportions of the other causes must decrease). However, this negative correlation became even more statistically significant after using additive log-ratio transformations^47,48^ that are typically used to control for this source of correlation (PGLS, *P-*value < 2 x 10^-16^ , lambda = 0.33, R² = 0.48, regression coefficient = – 0.08). Cancer and trauma are not the only causes of death in the examined mammals, and in fact make up an average of 39.5% of deaths for any given species (range 5.9% to 73.6%) (Supp. Fig. 15). When controlling the trauma mortality ∼ cancer mortality regression for either platelet size, platelet concentration, maximum lifespan, or body mass and trophic level, platelet size (Table 1; PGLS, *P-*value = 0.06, regression coefficient = –0.01) and secondary carnivores (Table 1; PGLS, *P-*value = 0.06, regression coefficient = –0.13), but not platelet concentration (Table 1; PGLS, *P*-value = 0.75), maximum lifespan (Table 1; PGLS, *P*-value = 0.25), or body mass (Table 1; PGLS, *P*-value = 0.40), partially mediated the negative correlation between trauma and cancer mortality. Furthermore, trauma mortality did not correlate with the segmented neutrophil-to-lymphocyte ratio (n = 70 species; Table 1). Platelet size negatively correlated with trauma mortality across 113 mammalian species (Fig. 1; PGLS, *P-*value = 0.0002, regression coefficient = –0.02). This correlation remained even when controlling the analysis for just trophic level, or all three of the following variables: trophic level, body mass, and gestation length (Table 1). Median platelet size and concentration were inversely correlated (n = 111 species; PGLS, *P-*value = 0.001, regression coefficient = –0.01; Supp. Fig. 18), and median platelet size (Supp. Fig. 17) nor concentration did not correlate with trophic level (Table 1). Higher trophic levels had greater cancer prevalence (Table 1; PGLS, *P-*value ≤ 0.007, primary or secondary carnivores versus herbivores, regression coefficient > 0) and cancer mortality (Table 1; PGLS, *P-*value ≤ 0.002, invertivores or primary carnivores versus herbivores, regression coefficient > 0), but less trauma mortality (Fig. 5; PGLS, *P-*value = 0.01, regression coefficient = –0.17). Cancer prevalence and cancer mortality were positively correlated (Supp. Fig. 19; Table 1).

**Figure 3.**
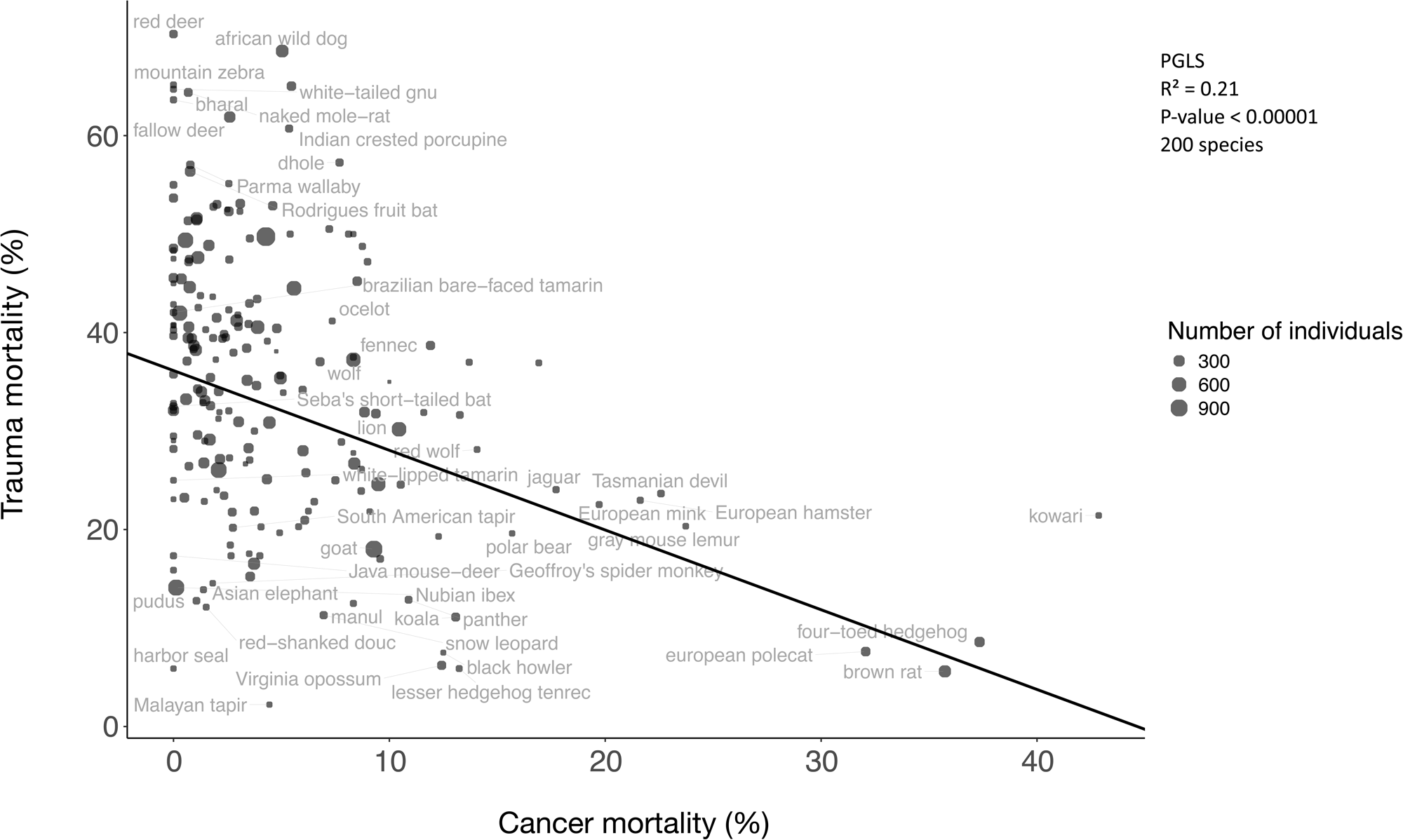
Trauma mortality is negatively correlated with cancer mortality. Each dot shows the trauma mortality and cancer mortality of one species. The regression line shows the phylogenetically-controlled linear regression between trauma mortality and cancer mortality.

**Figure 4.**
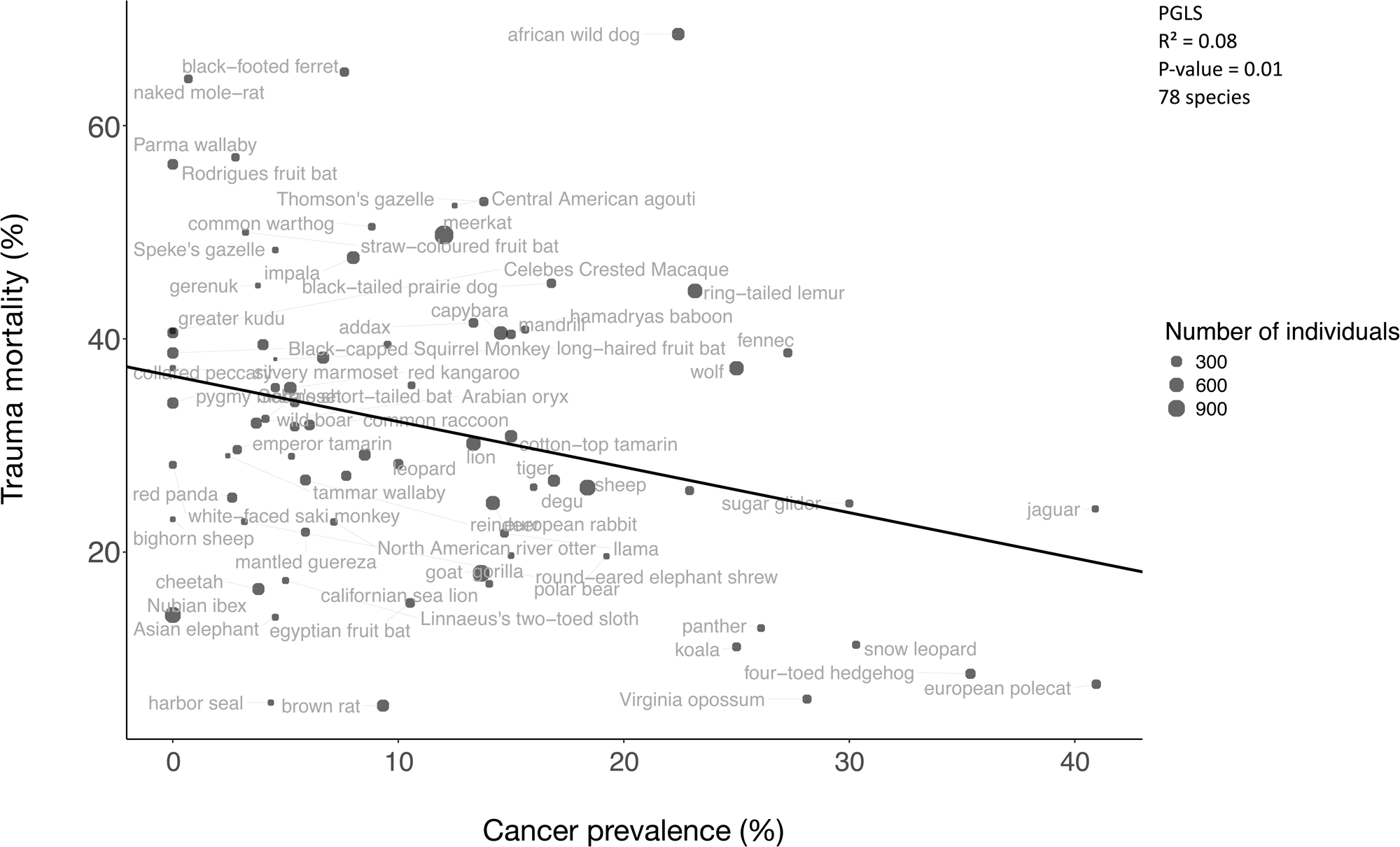
Trauma mortality is negatively correlated with cancer prevalence. Each dot shows the trauma mortality and cancer prevalence of one species. The regression line shows the phylogenetically-controlled linear regression between trauma mortality and cancer prevalence.

**Figure 5.**
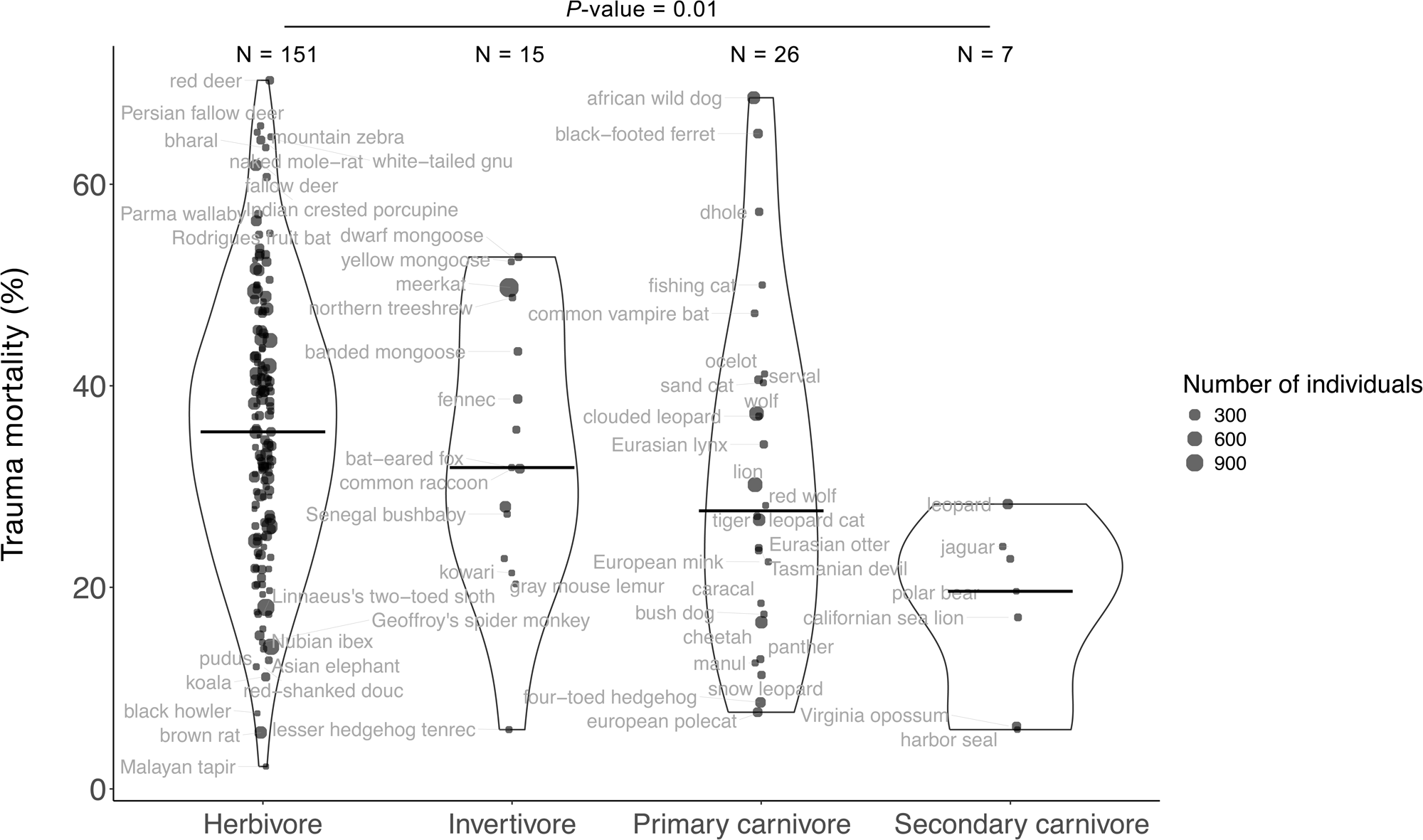
Trauma mortality is negatively correlated with trophic level. Each dot shows the trauma mortality and trophic level of one species. The *P*-value comes from a phylogenetically-controlled linear regression between the arcsine-square root transformed trauma mortality and trophic level variables. N refers to the total number of species in the trophic level.

Two examples are illustrative. The straw-colored fruit bat (*Eidolon helvum*) is an example of a species that is a herbivore with median platelet size 6.6 fL (from 49 different individuals), 0.478 x 10^12^ platelets/L of whole blood (from 180 different individuals)^49^, 50% trauma mortality (from 74 different individuals)^50^, 3.2% cancer prevalence (from 31 necropsies), and 5.4% cancer mortality (from 74 different individuals)^50^. In contrast, the European polecat (*Mustela putorius*) belongs to a higher trophic level, it has almost one and a half times as large platelets (9.5 fL from 52 different individuals), lower platelet concentration (0.29 x 10^12^ platelets/L of whole blood from 351 different individuals)^51^, more than six times lower trauma mortality (7.6% trauma mortality from 184 different individuals)^50^, more than 12 times higher cancer prevalence (41% cancer prevalence from 105 necropsies), and around six times higher cancer mortality (32% cancer mortality from 184 different individuals)^50^.

## Discussion

Evolution can shape disease risk through trade-offs, such as antagonistic pleiotropy, where a trait that offers benefits in one context imposes costs in another. The ability to repair tissue after injury is essential for survival, especially in species that have high trauma risk. This ability to rapidly regenerate tissue may come at a cost to cancer risk later in life with enhanced signaling pathways for cell growth. Our discovery of a negative correlation between trauma and cancer mortality supports this hypothesis.

The relationship between trauma and cancer mortality is partially mediated by platelet size and trophic level. Carnivores and herbivores experience different ecological pressures due to how they extract resources from their environments and the risks involved in these extractions. We found that species at higher trophic levels had less trauma mortality than species at lower trophic levels. This suggests that carnivores may have evolved rapid and efficient wound healing abilities compared to herbivores, though they may also suffer less trauma. Some observations suggest that wounds of carnivores heal faster than the wounds of herbivores (Tara Harrison, DVM, pers. comm.). Wound healing and cancer share common biological pathways, including cell proliferation, angiogenesis, cell migration and invasion, immune evasion, inflammation, and resistance to cell death^10^. These hallmarks may be enhanced in carnivores, compared to herbivores, aiding more efficient and effective wound repair and lower trauma mortality.

Platelets are small blood cell fragments involved in blood clotting, stopping a wound from bleeding, and regulating blood hemostasis. The concentration and size of platelets can indicate activation. Platelets increase in size when activated and larger platelets can have better clotting ability^52–54^. Larger platelets have been shown to clot more effectively in rabbits^52^. In humans, aggregation of platelets occurs faster when platelets are larger^53^ and surgical patients with larger platelets were less likely to experience postoperative bleeding^54^. Patients with severe injuries do not have significantly different platelet sizes than controls, suggesting that the injuries were not causing a growth in platelet size^55^. In another study, patients with mild head trauma had smaller platelets than healthy controls^56^. Fractures in patients that have smaller platelets heal faster than those with larger platelets^57^. These findings suggest that large platelets facilitate the acute process of blood clotting but small platelets facilitate the slow process of wound healing. A small comparative study reported that two carnivorous species (cats and dogs) had larger normal baseline mean platelet size than three herbivores: horses, cows, and goats^45^.

However, in our study of ≥110 mammalian species, carnivores and herbivores did not differ significantly in platelet size. This suggests that the relationship between trophic level and trauma mortality is not explained by platelet size. Independent of trophic level, we found that platelet size negatively correlated with trauma mortality, and there was a positive trend between platelet size and cancer mortality that did not pass FDR correction (Table 1; *P*-value = 0.02). Platelet size also appeared to explain part of the relationship between cancer and trauma mortality (Table 1; multivariate regression, *P-*value = 0.06). Beyond clotting, platelets are involved in inflammation^31^ and cancer biology^58^. Tumor cells can activate and induce the aggregation of platelets^59–61^, and active inflammation (often seen following platelet activation) has been associated with cancer development^62–64^. Depending on the tumor cell type and microenvironment, the interaction between tumor cells and platelets is complex^65,66^. Studies show platelets may inhibit tumor growth^26^, protect tumor cells from the immune system^67^ and potentially promote metastasis^26–29,65,66^.

Notably, platelet size and concentration are inversely correlated^53^. However, the connection of platelet concentration with trophic level, cancer prevalence/mortality, and trauma mortality is not as clean as we might hope. Higher trophic levels did not have statistically significantly different concentrations of platelets than lower trophic levels. Mammals with higher platelet concentrations have lower cancer prevalence (Table 1; *P*-value ≤ 0.006), but not lower cancer mortality (Table 1; *P*-value = 0.12) nor higher trauma mortality (Table 1; *P*-value ≥ 0.37). These mixed patterns (Fig. 6; Supplementary Movie) warrant larger, phylogenetically controlled studies with standardized platelet indices to establish whether the observed associations are robust.

**Figure 6.**
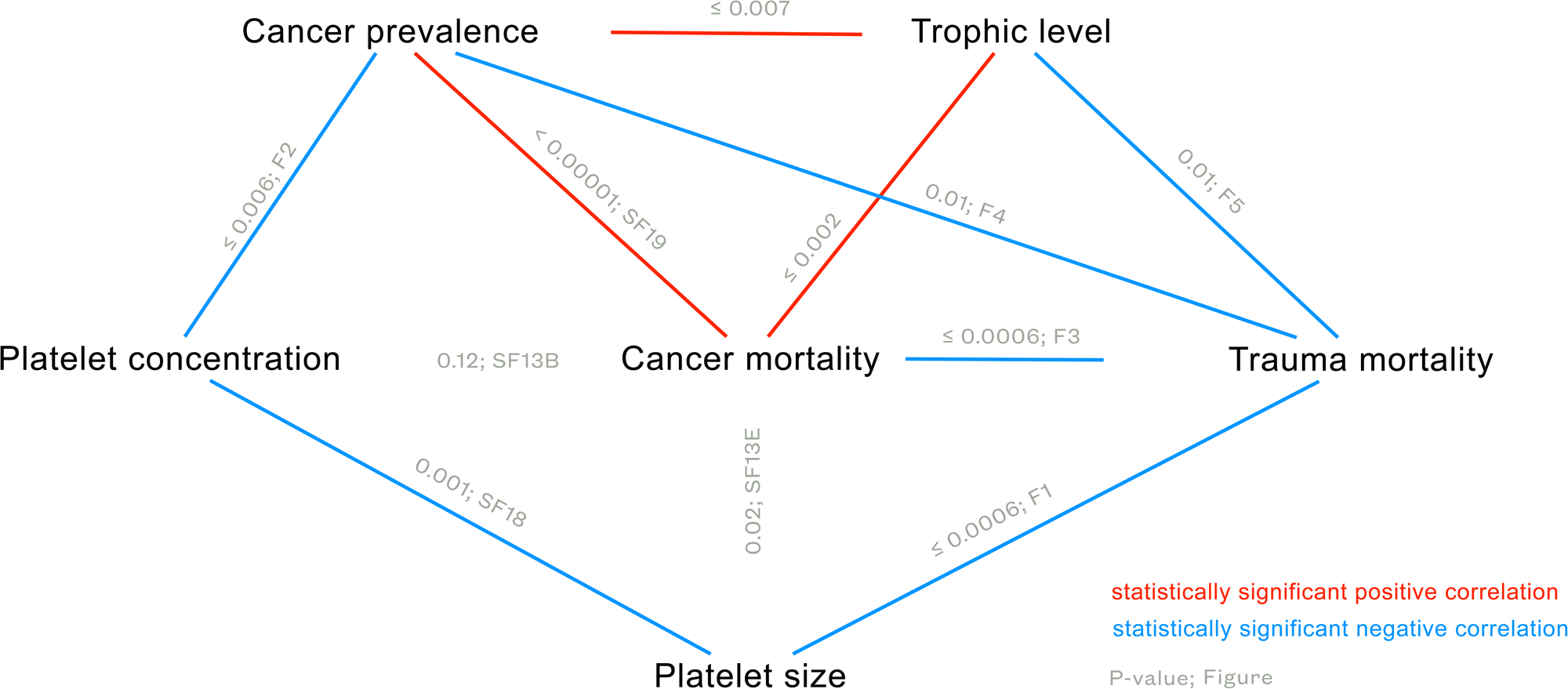
Schematic of the summary findings of correlations between platelets, trauma mortality, cancer mortality, cancer prevalence, and trophic levels. Variables that are connected with a red line are positively correlated. Variables that are connected with a blue line are negatively correlated. Variables that are not connected are not statistically significantly correlated after multiple testing corrections. We provide the detailed statistics of each regression in Table 1. Each association is labeled with the *P*-value and the corresponding figure number for that relationship (F for figure and SF for supplementary figure). We do not provide a figure for the correlation between cancer prevalence and trophic level because this has been previously shown using a slightly different group of species^44,119^. Similarly, we do not provide a figure for the correlation between cancer mortality and trophic level because this has already been shown using a different cancer mortality dataset and different measurement of diet^6^.

### Innate immunity scales with size, not lifespan

Neutrophil-to-lymphocyte ratio is a common immune marker that can indicate immune activation. Our results show that larger mammals have a higher neutrophil-to-lymphocyte ratio and a lower lymphocyte concentration than smaller mammals. Previous studies in mammals have also found a higher concentration of neutrophils and lower concentration of lymphocytes in larger species^21,22,24^. Larger host species have been shown to have higher parasite richness than smaller host species (across 488 vertebrate species^68^; across 96 Artiodactyla and Perissodactyla species^69^; across 138 bat species^70^). Downs et al.^21^ propose the performance-safety tradeoff hypothesis^71^ as an explanation for the higher concentration of neutrophils in larger compared to smaller mammalian species. Larger mammals may have evolved better safety mechanisms as in a higher concentration of innate immune cells, such as neutrophils, as the first line of defense, quickly responding to pathogens^72,73^. However, a high baseline concentration of innate immune cells in one species versus another does not necessarily mean that the former species will have a higher concentration of innate immune cells than the latter during infection. Additionally, higher immune cell concentration does not always mean better immune function^74^. Clarifying these relationships requires data linking baseline levels to within-individual responses under infectious challenge.

Our results imply a complex relationship between immune cell concentrations, body size, and cancer prevalence. Innate immune cell profiles scale with body size, where larger mammals show higher neutrophil-to-lymphocyte ratios and lower lymphocyte concentrations than smaller mammals. Yet this does not yield a straightforward protective effect against cancer (PGLS; *P*-value ≥ 0.05 for associations of neutrophil-to-lymphocyte ratios and lymphocyte concentrations with cancer mortality; *P-*value ≥ 0.54 for associations of neutrophil-to-lymphocyte ratios and lymphocyte concentrations with cancer prevalence).

Our previous analyses of the relationship between body size and cancer prevalence were based on 255 vertebrates, including 100 mammals^2^. In univariate analyses, body size is not associated with cancer prevalence or mortality, consistent with our previous results, based on similar data^2,4,6^. After adjusting for gestation time, we found that body mass is again positively associated with cancer prevalence^2^. However, in a new analysis, we find that cancer mortality is still not associated with body mass, even when controlling for gestation time.

### Limitations

This study offers a broad, first-pass comparative view of mammalian immune cell concentrations in relation to cancer risk, wound-healing ecology, and body size. Nonetheless, there are inherent limitations to broad comparative analyses. Notably, immune function encompasses many dynamic processes that extend beyond the static measures of immune cell concentrations typically available in cross-species datasets. Species360’s Zoological Information Management System (ZIMS; https://species360.org/) Expected Test Result resource includes the minimum and maximum values observed for many immune cell measurements within species. However, we have not analyzed associations between the variance of immune cell measurements within species and mortality outcomes. Even if we had access to individual level data, the data for any one animal is often incomplete. We obtained aggregated data across healthy animals from the same species and across institutions. This inevitably injects noise. In addition, it is difficult to find data on the following variables that have been shown to affect immune cell values in certain species^75–82^. Immune cell concentrations are associated with an animal’s age^77,83–91^, season^82,92–95^ or time of day of blood withdrawal, moisture^96^, pregnancy, inflammation^97^, levels of thrombocytes, body temperature, promiscuity^98–100^, husbandry conditions, social organization^101^, population density, exercise^97^, levels of stress induced by capturing, handling, and sampling^97,102^, feeding behavior, and exact diet. Measuring these components along with clinical outcomes in every individual would give us more insight on the interactions of these components. Technology developed for measuring immune cells in human blood has not been validated in all mammalian species in our data set. It is difficult to distinguish nucleated red blood cells from white blood cells in birds and reptiles, thus ZIMS data for immune cells from these species in ZIMS are currently not ideal for comparative studies. Accordingly, patterns reported in this paper should be viewed as hypothesis-generating rather than definitive, motivating future work with standardized immune and platelet measures, longitudinal or experimental designs, and explicit phylogenetic controls to test mechanisms. Future work might also genetically manipulate cancer pathways, platelet size and/or wound healing pathways in traditional laboratory model systems to test if those genetic components mediate a tradeoff between cancer suppression and wound healing.

We have used trauma mortality (in mammals under managed care) as a proxy for wound healing. We acknowledge that the cause of death is sometimes difficult to determine in species and accordingly, there may be species that tend to die from trauma because they have proportionally thinner bones and/or are prone to take more risks, regardless of the quality of their wound healing. However, there was no relationship between body mass and trauma mortality in our multivariate analyses (Table 1). There may be other species with poor wound healing capabilities that don’t tend to suffer much trauma (and so have not been under selection to evolve improved wound healing). In the institutions where the animals are housed, some species are housed separately to reduce the risk of intra- and inter-species aggression. They may be brought together only when receptive for breeding to reduce the risk of aggression. This management strategy reduces the frequency of trauma-related wounds, so perhaps they are less likely to experience mortality from traumas, because they are less likely to incur significant traumatic injuries in the first place. This may explain the low trauma mortality in some species of our dataset, however, we do not have data on which species in each institution were housed separately and for how long. Future studies should develop direct measures of the various components of wound healing, as well as the different causes of trauma mortality, so as to identify any tradeoffs with cancer susceptibility. Lastly, comparing data from animals under managed care versus wild animals will also be valuable in order to identify whether the environment impacts immunity and the tradeoff between wound healing and cancer defenses.

## Conclusion

Our comparative study supports a fitness tradeoff hypothesis for cancer risk, suggesting an evolutionary compromise between cancer suppression and wound healing across mammals. We propose that this tradeoff may arise from shared underlying mechanisms, which manifest functionally as overlapping hallmarks between wound healing and cancer development^10^. This potential fitness trade-off between cancer and trauma mortality may be mediated partially by platelet size, and trophic level, but not by body mass, maximum lifespan, or platelet concentration. Unravelling exactly how and why these correlations exist may require functional studies as well as further correlation studies with more physiological and ecological data from both wild animals and animals under human care. Such data will enable evolutionary biologists, veterinarians, and medical doctors to better understand the interplay between cancer, trauma mortality, diet, life history evolution, and immune cells across species.

## Methods

### Collection of immune cell data

We collected immune cell data for 184 mammal species from ZIMS^103^; a database with records from over 1400 institutions (aquariums, zoos, universities, research and government institutions) across 102 countries. All immune data are from the whole blood of healthy animals^104^. Specifically, these data included the median concentration of lymphocytes (automated, i.e., counted using automated cell counters) (in 10^6^ cells/mL), monocytes (automated) (in 10^6^ cells/mL), segmented neutrophils (automated) (in 10^6^ cells/mL), white blood cells (automated) (in 10^6^ cells/mL), and platelets (automated) (in 10^9^ cells/mL), and the median values of the mean platelet volume data (automated) (in fL or μm³); where mean platelet volume is a derived measurement of the average size of platelets^105^. Non-zero percentages of band neutrophils were only available for four species in the ZIMS database, therefore it was not possible to add the concentration of band neutrophils and the concentration of segmented neutrophils to calculate the concentration of neutrophil values across all species in our dataset. We measured the segmented neutrophil-to-lymphocyte ratio by dividing the median concentration of segmented neutrophils by the median concentration of lymphocytes, with the caveat that these two immune cell types were not always measured from the same individuals in each species. We only included median immune cell data that were available from at least 20 animals per species in the ZIMS database. These immune cell data were not available for every species for which we had cancer prevalence or cancer mortality data.

### Collection of cancer prevalence, cancer mortality, trauma mortality, and life history data

We used cancer prevalence data from the Arizona Cancer Evolution (ACE) Center’s dataset^2,106,107^ and cancer mortality data from ZIMS^50^. The ACE data included cancer prevalence from 5,843 necropsy records across 102 mammalian species. These data are gathered from at least 20 individuals per species from several zoological and veterinary institutions^2,106,107^. We have updated the cancer prevalence and necropsy data of *Didelphis virginiana* in comparison to our previous studies^106,107^ and in comparison to the *Didelphis marsupialis* data point in ^2^, because the previous necropsy records of *Didelphis virginiana* and *Didelphis marsupialis* actually belonged to one species (*Didelphis virginiana*). Using data from 37,072 animals with a single reported circumstance related to death in the ZIMS database, we measured cancer mortality, i.e., the percent of animals per species that died from cancer, from 211 mammalian species (Supp.Fig.14H) with records from at least 20 different individuals per species. The date range of the query we used was January 10th, 2010 to April 29th, 2024. Note that this dataset largely overlaps with the data reported in ^108^ (date range of their query: January 1st, 2010 to May 30th, 2020). The methodology for measuring cancer mortality was the same as previously described^5^. We collected trauma mortality data, i.e., the percent of animals per species that died from trauma, where trauma was the single reported circumstance related to death, from the ZIMS database for the same 211 species (Supp.Fig.14I). We only collected trauma mortality data from species that had records from at least 20 different individuals. The methodology for collecting trauma mortality data was similar to the methodology previously used to collect cancer mortality data^5^ with the exception that the numerator was the total number of animals in that species that had died from trauma. In both cases of cancer mortality and trauma mortality, the denominator only had data of animals reported to die of a single cause, not multiple causes. We collected adult body mass (grams) data for each species from Compton et al.^2^, and from additional species, not reported in Compton et al.^2^, from AnAge^109^. We also collected gestation length (months), and lifespan (months) data from previous comparative oncology studies^2^, AnAge^110^, and Animal Diversity^111^. We obtained trophic level data for 112 out of 216 species in our dataset from a previously published dataset^106^, and for 98 species from the IUCN red list of threatened species and Animal Diversity (https://animaldiversity.org/).

### Statistical analyses

We conducted all analyses in R version 4.0.5^112^ using the packages CAPER^113^, phytools^114^, geiger^115^, and tidyverse^116^. We tested the analyses for statistically significant heteroscedasticity (Fligner-Killeen test), and if present, we mention this in our results. We used Grubbs’ and Rosner’s tests to test for outliers in distributions. In order to control for the phylogenetic non-independence between species, we conducted phylogenetic generalized least squares (PGLS) regressions using a phylogenetic tree from TimeTree (timetree.org). In each analysis that had a subset of species from the dataset, we pruned this phylogenetic tree using the functions setdiff and keep.tip/drop.tip in R. We also weighted the species data points in the PGLS analyses by 1/(square root of the number of records per species of the y variable) (from ^114^) to control for the variation in the number of records or necropsies sampled per species. Due to the proportional nature of the cancer prevalence, neoplasia prevalence, cancer mortality, and trauma mortality data, we arcsine-square-root-transformed those data in the PGLS analyses, but present the non-transformed data in the figures to allow visualization of the original data. We obtained the ‘R2.resid’ values using the ‘rr2’ package (R2(mod = main_model, mod.r = reduced_model, phy = pruned.tree); where main_model is the univariate or multivariate model that has one or more independent variables (not the null model), and reduced_model is the model where the independent variable is 1 (null model))^117^. We performed False Discovery Rate (FDR) corrections for multiple testing in the group of PGLS analyses where: (1) maximum lifespan is the independent variable and immune cells are the dependent variable (7 tests); (2) body mass is the independent variable and immune cells are the dependent variable (7 tests); (3) immune cells are the independent variable and cancer mortality or cancer prevalence is the dependent variable (16 tests); (4) immune cells are the independent variable and trauma mortality is the dependent variable (7 tests); (5) trophic level is the independent variable (28 tests).

Due to body mass correlating with lifespan^118^, and body mass correlating with the concentration of white blood cells, lymphocytes, monocytes, and neutrophils^24^, we controlled for lifespan in the bivariate analyses where body mass was one of the independent variables, whereas we controlled for body mass in the bivariate analyses where lifespan was one of the independent variables. Because previous studies have reported associations of cancer prevalence with gestation length^2^, neoplasia prevalence with trophic levels^44^, and neoplasia prevalence with body mass across vertebrates^2^, we controlled for gestation length, body mass, and trophic levels in the correlations between “trauma mortality ∼ platelet size” and “cancer prevalence ∼ platelet concentration” (Table 1). Given that the platelets of Felidae species tend to aggregate more than other species^46^ leading to potential inaccuracies in automated measures of platelets, we also tested whether the correlation between cancer prevalence and platelet concentration remained when excluding the Felidae. We also controlled the “trauma mortality∼cancer mortality” analyses for species’ maximum lifespan (Table 1) given that lifespan is positively correlated with neoplasia prevalence across vertebrates^2^.

To compare the immune profiles, cancer mortality, and trauma mortality of different taxonomic groups (orders), we performed Kruskal-Wallis tests.

To conduct an analysis that addresses the issue of cancer mortality and trauma mortality being negatively correlated due to those values of a given species having the same denominator (compositionality), we applied an additive log-ratio transformation to the mortality statistics^47,48^. For each species i, we computed the value z_i by: z_i = log(x_i/y_i), where x_i was cancer mortality and y_i was trauma mortality. Then, using PGLS we fitted the regression Y = beta_0 + beta_1*Z, where Y and Z were vectors containing y_i’s and z_i’s.

## Supplementary Material

### Supplementary Information

In our dataset of 216 mammalian species across 18 different taxonomic orders, we found that different taxonomic orders have different median concentrations of white blood cells (Supp. Fig. 14A; no statistically significant outliers), lymphocytes (Supp. Fig. 14B; with 2 statistically significant outliers: the Egyptian fruit bat *Rousettus aegyptiacus* and the Linnaeus’s two-toed sloth *Choloepus didactylus*), monocytes (Supp. Fig. 14E; with 2 statistically significant outliers: the Egyptian fruit bat *Rousettus aegyptiacus* and the Celebes crested macaque *Macaca nigra*), platelet size (Supp. Fig. 14F; no statistically significant outliers), platelet concentration (Supp. Fig. 14G; no statistically significant outliers), cancer mortality (Supp. Fig. 14H; with 8 statistically significant outliers: the European mink *Mustela lutreola,* the European hamster *Cricetus cricetus,* the Tasmanian devil *Sarcophilus harrisii,* the gray mouse lemur *Microcebus murinus,* the European polecat *Mustela putorius,* the brown rat *Rattus norvegicus,* the four-toed hedgehog *Atelerix albiventris,* and the kowari *Dasyuroides byrnei*), and trauma mortality (Supp. Fig. 14I; no statistically significant outliers). However, the concentration of segmented neutrophils (Supp. Fig. 14C; with one statistically significant outlier: the pygmy marmoset *Cebuella pygmaea*) or the segmented neutrophil-to-lymphocyte ratio (Supp. Fig. 14D) does not differ among the examined taxonomic orders.

The reason for the very high concentration of immune cells in outlier species is largely unknown. There were two outlier species with relatively high lymphocyte concentrations in our dataset, the Egyptian fruit bat *Rousettus aegyptiacus* and the Linnaeus’s two-toed sloth *Choloepus didactylus*. Vogel et al.^120^ found a similar mean value of 5.0 x 10^9^ cells/L lymphocytes from 66 wild-caught healthy individual Linnaeus’s two-toed sloths (*Choloepus didactylus*) to the median value of 5.78 x 10^9^ cells/L lymphocytes from 94 healthy individual Linnaeus’s two-toed sloths (*Choloepus didactylus*) under human care^121^ in our study. However, the reason for the relatively high lymphocyte concentrations in these species is unknown. The Egyptian fruit bat (*Rousettus aegyptiacus*) was also an outlier species with very high (9.2 x 10^9^ cells/L; from 26 different individuals; both healthy male and female Egyptian fruit bats *Rousettus aegyptiacus* in ZIMS of unspecified animal age^122^) monocyte concentrations, whereas previous literature in 3 mock-inoculated juvenile male Egyptian fruit bats (*Rousettus aegyptiacus*) has reported monocyte concentrations lower than 2 x 10^9^ cells/L^123^. Celebes crested macaques (*Macaca nigra*), are an outlier species with very high monocyte concentrations; a median across 203 healthy individuals^124^. There is no reference value of monocyte counts in previous literature for this species and no clear explanation as to why this species has such high monocyte concentrations. Similarly in the case of the pygmy marmoset (*Cebuella pygmaea*), an outlier with relatively high concentrations of segmented neutrophils across 65 healthy individuals, there is no previous literature on reference counts of segmented neutrophils for this species or a clear explanation for its high segmented neutrophil concentrations. The way that these animals were handled for blood collection, potentially anesthesia or manual restraint, may have affected the concentrations of these immune cells^97^. In the ZIMS database, consistent data entry errors, confusion over test methodology, or unit of measurement conversations over time by different individuals entering the data may also have an impact on data quality.

Regarding immune profiles and body mass, our comparative analyses showed distinct immune profiles in larger versus smaller species but no indication of which immune profiles are better at preventing mortality from cancer cells, ‘non-self’ cells, and/or autoimmune diseases. Our findings of higher segmented neutrophil-to-lymphocytes ratios and lower lymphocyte concentrations in larger species align with previous findings in some taxa. Downs et al.^21^ and Naidenko and Alshinetskiy^24^ also used the ZIMS database to collect species’ immune cell data but from a different time period (which explains why their values may not be exactly the same as ours). Downs et al.^21^ found that body mass is positively correlated with neutrophil concentrations across 259 terrestrial mammalian species, and Naidenko and Alshinetskiy found that body mass is negatively correlated with lymphocyte concentration across 25 felids (*P-*value = 0.0094)^24^. Without measures of immune cell function across species it is difficult to understand why and how these correlations exist, and which immune profiles are indicative of ability to prevent death from different threats.

All *P*-values in the article are un-adjusted for multiple testing corrections.

## Supplementary Movie

Figure 6 in the form of a movie. Variables that have the same font size at the same time are positively correlated (a trend: no line joining them; statistical significance after FDR correction: a line joining them). Whereas when one variable has a large font size and the other has a small font size at the same time, this represents that they are negative correlated (a trend: no line joining them; statistical significance after FDR correction: a line joining them).

## Supplementary Data

The species-level life history, immune cell, cancer prevalence, cancer mortality, and trauma mortality data used in this study. We will share these data upon acceptance of the manuscript for publication.

**Supplementary Figure 1. Body mass (A) and lifespan (B) are not correlated with the median concentration of white blood cells.** The analysis in A is controlled for lifespan, whereas the analysis in B is controlled for body mass. Each dot shows the median concentration of white blood cells and body mass (A) or the lifespan (B) of one species. The regression lines show the phylogenetically-controlled linear regression of body mass (A) and lifespan (B) versus the median concentration of white blood cells.

**Supplementary Figure 2. Body mass (A) and lifespan (B) are not correlated with the median concentration of segmented neutrophils.** The analysis in A is controlled for lifespan, whereas the analysis in B is controlled for body mass. Each dot shows the median concentration of segmented neutrophils and body mass (A) or the lifespan (B) of one species. The regression lines show the phylogenetically-controlled linear regression of body mass (A) and lifespan (B) versus the median concentration of segmented neutrophils.

**Supplementary Figure 3. Body mass (A), but not lifespan (B), is negatively correlated with the median concentration of lymphocytes.** The analysis in A is controlled for lifespan, whereas the analysis in B is controlled for body mass. Each dot shows the x and y variables of one species. The regression lines show the phylogenetically-controlled linear regression of body mass (A) and lifespan (B) versus the median concentration of lymphocytes.

**Supplementary Figure 4. Body mass (A), but not lifespan (B), is positively correlated with the ratio of segmented neutrophils-to-lymphocytes.** Brackets indicate concentrations. The analysis in A is controlled for lifespan, whereas the analysis in B is controlled for body mass. Each dot shows the x and y variables of one species. The regression lines show the phylogenetically-controlled linear regression of body mass (A) and lifespan (B) versus the ratio of segmented neutrophils-to-lymphocytes.

**Supplementary Figure 5. Body mass (A) and lifespan (B) are not correlated with the median concentration of monocytes.** The analysis in A is controlled for lifespan, whereas the analysis in B is controlled for body mass. Each dot shows the median concentration of monocytes and body mass (A) or lifespan (B) of one species. The regression lines show the phylogenetically-controlled linear regression of body mass (A) and lifespan (B) versus the median concentration of monocytes.

**Supplementary Figure 6. Body mass (A) and lifespan (B) are not correlated with median platelet size.** The analysis in A is controlled for lifespan, whereas the analysis in B is controlled for body mass. Each dot shows median platelet size and body mass (A) or lifespan (B) of one species. The regression lines show the phylogenetically-controlled linear regression of body mass (A) and lifespan (B) versus median platelet size.

**Supplementary Figure 7. Body mass (A) and lifespan (B) are not correlated with median platelet concentration.** The analysis in A is controlled for lifespan, whereas the analysis in B is controlled for body mass. Each dot shows median platelet concentration and body mass (A) or lifespan (B) of one species. The regression lines show the phylogenetically-controlled linear regression of body mass (A) and lifespan (B) versus median platelet concentration.

**Supplementary Figure 8. The median concentration of segmented neutrophils is not correlated with cancer prevalence (A) nor cancer mortality (B).** Each dot shows the cancer prevalence (A) or cancer mortality (B) and the median concentration of segmented neutrophils of one species. The regression lines show the correlation between the median concentration of segmented neutrophils and cancer prevalence (A) or cancer mortality (B).

**Supplementary Figure 9. The ratio of segmented neutrophils-to-lymphocytes is not correlated with cancer prevalence (A) nor cancer mortality (B).** Brackets indicate concentrations. Each dot shows the cancer prevalence (A) or cancer mortality (B) and the ratio of segmented neutrophils-to-lymphocytes of one species. The regression lines show the correlation between the ratio of segmented neutrophils-to-lymphocytes and cancer prevalence (A) or cancer mortality (B).

**Supplementary Figure 10. The median concentration of lymphocytes is not correlated with cancer prevalence (A) nor with cancer mortality (B).** Each dot shows the cancer prevalence (A) or cancer mortality (B) and the median concentration of lymphocytes of one species. The regression lines show the correlation between the median concentration of lymphocytes and cancer prevalence (A) or cancer mortality (B).

**Supplementary Figure 11. The median concentration of monocytes is not correlated with cancer prevalence (A) nor cancer mortality (B).** Each dot shows the cancer prevalence (A) or cancer mortality (B) and the median concentration of monocytes of one species. The regression lines show the correlation between the median concentration of monocytes and cancer prevalence (A) or cancer mortality (B).

**Supplementary Figure 12. The median concentration of white blood cells is not correlated with cancer prevalence (A) nor cancer mortality (B).** Each dot shows the cancer prevalence (A) or cancer mortality (B) and the median concentration of white blood cells of one species. The regression lines show the correlation between the median concentration of white blood cells and cancer prevalence (A) or cancer mortality (B).

**Supplementary Figure 13. Correlations between platelet size and concentration, cancer prevalence, cancer mortality, trauma mortality, and lifespan.** There are no correlations between the examined variables (A–E). Each dot shows the x and y variables of one species. Each regression line shows the phylogenetically-controlled linear regression between the independent and dependent variable.

**Supplementary Figure 14. Immune profiles, cancer mortality, and trauma mortality across different mammalian taxonomic orders.** The median concentration of white blood cells (A), monocytes (E), platelet size (F), platelet concentration (G), cancer mortality (H), and trauma mortality (I) are different among orders. Each dot shows the x and y variables of one species. The horizontal line in each violin shows the median value of the y variable in the taxonomic order shown on the x axis.

**Supplementary Figure 15.** The sum of trauma and cancer mortality percentages across the examined species follow a normal distribution (211 species, Shapiro’s test: *P*-value = 0.61). Each point is a species.

**Supplementary Figure 16. When excluding the Felidae, cancer prevalence remains negatively correlated with platelet concentration.** Each dot shows the median platelet concentration and cancer prevalence of one species. The regression line shows the phylogenetically-controlled linear regression of median platelet concentration versus cancer prevalence.

**Supplementary Figure 17. There is a non-statistically significant trend for higher trophic levels to have larger platelets.** Each dot shows the median platelet concentration and cancer prevalence of one species. The horizontal line in each violin shows the median value of the y variable in the taxonomic order shown on the x axis. Herbivores do not have smaller platelets than the other trophic levels (P > 0.05; Table 1).

**Supplementary Figure 18. Median platelet size is negatively correlated with median platelet concentration.** Each dot shows the median platelet size and the median platelet concentration of one species. The regression line shows the correlation between median platelet size and median platelet concentration.

**Supplementary Figure 19. Cancer prevalence is positively correlated with cancer mortality.** Each dot shows the cancer prevalence and the cancer mortality of one species. The regression line shows the correlation between cancer prevalence and cancer mortality.

Table 1. **Summary phylogenetic regression (PGLS) statistics.** We note with an asterisk (*) on the *P*-value and **in bold the regressions that are considered statistically significant after the False Discovery Rate corrections**. High lambda values show that the associations are mainly explained by common ancestry. Multivariable additive PGLS models are shown as xrows with multiple independent variables but only one dependent variable. In the cases where trophic level is the independent variable, herbivores are the reference category. Prevalence and mortality data are on the unit interval and many values were near zero, so these data are arcsine-square root transformed for the regression analyses.

## Author contributions

A.M.B. conceived the idea to compare immune cell levels and cancer prevalence in December 2021. S.E.K.. Z.C., A.M.B., E.G.D., and T.M.H. helped collect and coordinate the cancer prevalence data across institutions. A.M.B., L.M.A., C.C.M., and S.E.K. initiated the idea to add trauma mortality analyses in the manuscript. L.M.A. conceived the idea to add the segmented neutrophil-to-lymphocyte ratio analyses. S.E.K. extended this idea to include life history traits, collected data on immune system components (concentrations of different immune cell types and median platelet size), trauma and cancer mortality data from ZIMS, and analyzed the data. S.M.R., Z.C., and A.M.B., provided life history data as well as cancer prevalence and necropsy data for each species. S.E.K. and S.S. searched for relevant articles in the literature. S.E.K., S.S., L.M.A., C.C.M., and A.M.B. generated hypotheses. S.E.K. wrote the first draft of the manuscript. S.S. wrote parts of the manuscript, and reviewed the analyses and figures under the guidance of A.M.B., L.M.A., and C.C.M. who also provided helpful comments and guidance throughout the project. All authors edited the final version of the manuscript.

## Acknowledgements

We thank all the pathologists, veterinarians, and staff at the zoos, aquariums, and private veterinary centers for their contribution in collecting data by diagnosing malignancies. Specifically, we would like to acknowledge the following institutions: Akron Zoo, Atlanta Zoo, Audubon Nature Institute, Bergen County Zoo, Birmingham Zoo, Buffalo Zoo, Capron Park Zoo, Central Florida Zoo, Dallas Zoo, El Paso Zoo, Elmwood Park Zoo, Fort Worth Zoo, Gladys Porter Zoo, Greensboro Science Center, Henry Doorly Zoo, Utah’s Hogle Zoo, Jacksonville Zoo, John Ball Zoo, Los Angeles Zoo, Louisville Zoo, Mesker Park Zoo, Miami Zoo, Oakland Zoo, Oklahoma City Zoo, Philadelphia Zoo, Phoenix Zoo, Pueblo Zoo, San Antonio Zoo, Santa Ana Zoo, Santa Barbara Zoo, Sedgwick County Zoo, Seneca Park Zoo, The Brevard Zoo, The Detroit Zoo, The Oregon Zoo, and Toledo Zoo. We also thank Esther Borges Florsheim, Rachel Thompson, Orsolya Vincze, and Mathieu Giraudeau for their valuable comments on the manuscript, and Ping-Han Huang for help with the code for measuring R² values, accounting for the issue of compositionality in the cancer and trauma mortality data, and controlling for the number of records in the PGLS analyses. Aktipis and Maley developed the wound healing-cancer defenses tradeoff hypothesis in conversations in 2012. We are grateful to OpenAI, and all the human knowledge that went into the construction of ChatGPT. In 2024, ChatGPT suggested that we consider wound healing when we asked it about the role of platelet size across animals, which reminded us of the hypothesis of a tradeoff between wound healing and cancer suppression. This work was supported in part by NIH grants U54 CA217376, U2C CA233254, P01 CA91955, and R01 CA140657 as well as CDMRP Breast Cancer Research Program Award BC132057 and the Arizona Biomedical Research Commission grant ADHS18-198847. Z.T.C. was supported by grant T32CA272303. The findings, opinions and recommendations expressed here are those of the authors and not necessarily those of the universities where the research was performed or the National Institutes of Health.

