## Supplementary Figures for "Immune-cancer analyses across mammals reveal a potential trophic level and platelet-linked tradeoff between cancer and trauma mortality"

Supp.Fig.1A

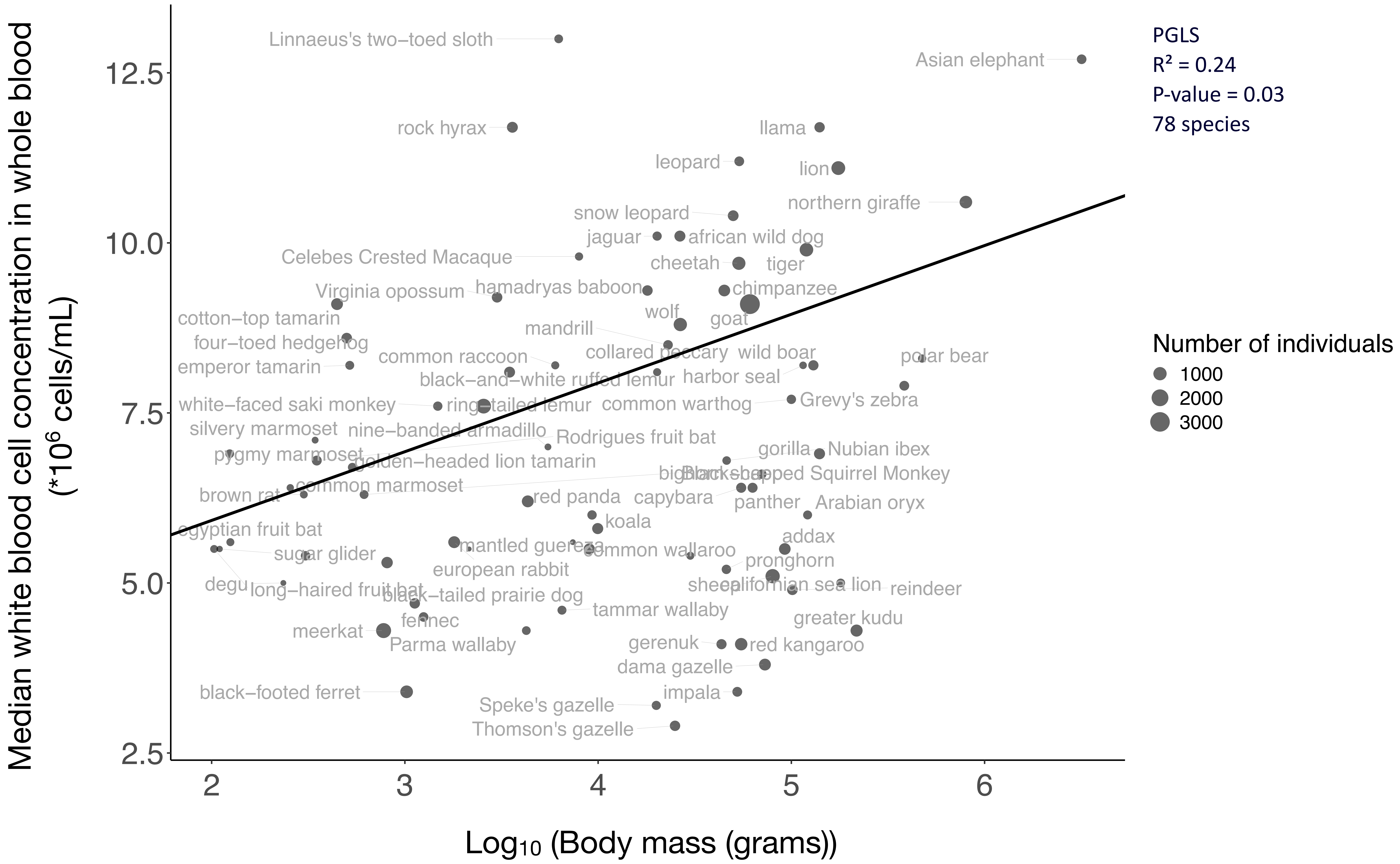

Median white blood cell concentration in whole blood
(*10^6 cells/mL)
Log10 (Body mass (grams))
PGLS
R² = 0.24
P-value = 0.03
78 species
Number of individuals
1000
2000
3000
12.5
10.0
7.5
5.0
2.5
2
3
4
5
6
Linnaeus's two–toed sloth
Asian elephant
rock hyrax
llama
leopard
lion
northern giraffe
snow leopard
jaguar
african wild dog
Celebes Crested Macaque
cheetah
tiger
Virginia opossum
hamadryas baboon
chimpanzee
cotton–top tamarin
wolf
goat
mandrill
four–toed hedgehog
common raccoon
collared peccary
wild boar
polar bear
emperor tamarin
black–and–white ruffed lemur
harbor seal
white–faced saki monkey
ring–tailed lemur
common warthog
Grevy's zebra
silvery marmoset
nine–banded armadillo
Rodrigues fruit bat
gorilla
Nubian ibex
pygmy marmoset
golden–headed lion tamarin
Black–capped Squirrel Monkey
common marmoset
brown rat
red panda
capybara
panther
Arabian oryx
koala
egyptian fruit bat
addax
sugar glider
mantled guereza
common wallaroo
pronghorn
european rabbit
degu
long–haired fruit bat
black–tailed prairie dog
californian sea lion
reindeer
tammar wallaby
fennec
greater kudu
meerkat
Parma wallaby
gerenuk
red kangaroo
dama gazelle
black–footed ferret
impala
Speke's gazelle
Thomson's gazelle

Supp.Fig.1B

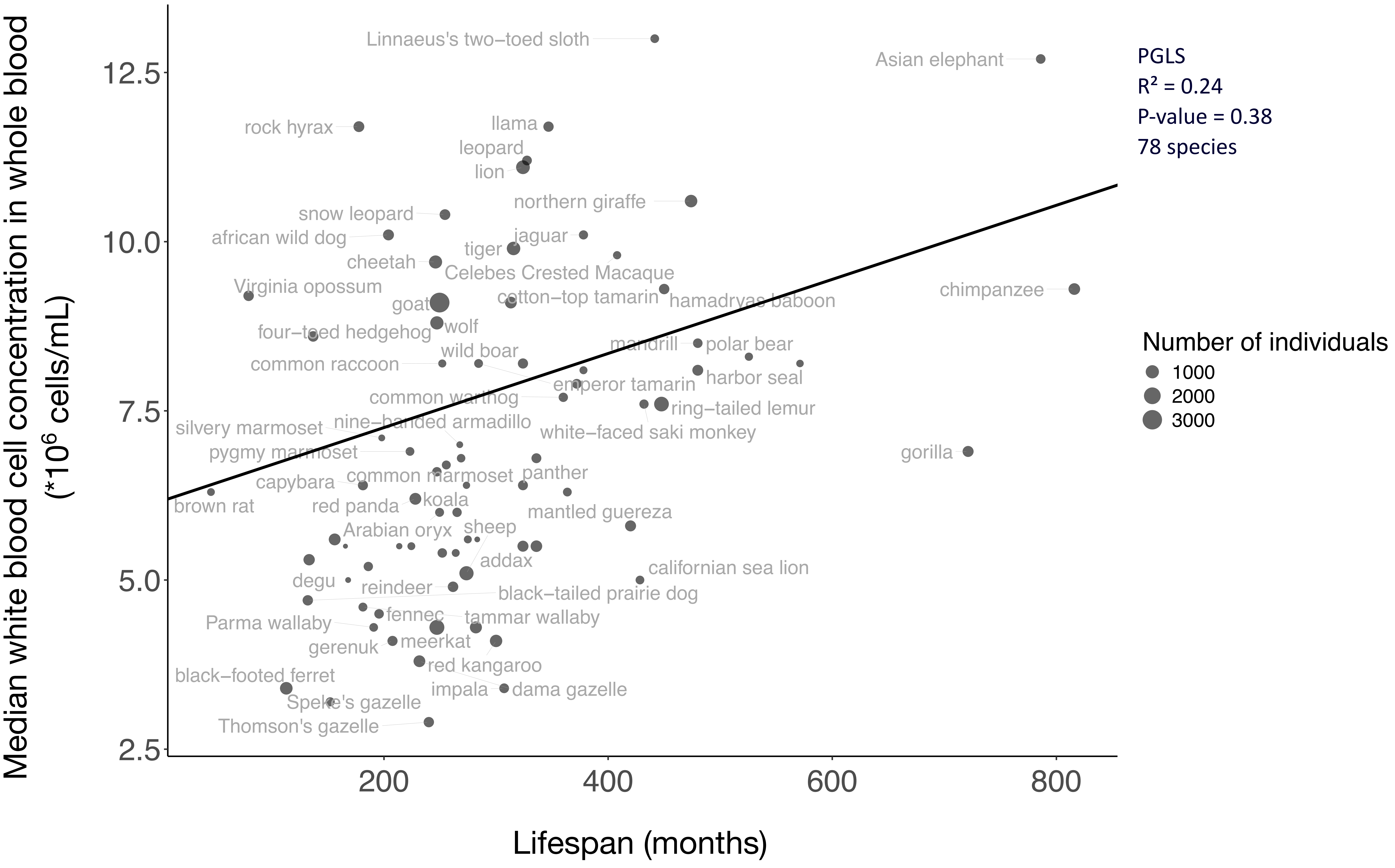

Median white blood cell concentration in whole blood (*10^6 cells/mL)
Lifespan (months)
12.5
10.0
7.5
5.0
2.5
200
400
600
800
PGLS
R² = 0.24
P-value = 0.38
78 species
Number of individuals
1000
2000
3000
Linnaeus's two-toed sloth
Asian elephant
rock hyrax
llama
leopard
lion
northern giraffe
snow leopard
african wild dog
jaguar
tiger
cheetah
Celebes Crested Macaque
Virginia opossum
chimpanzee
goat
cotton-top tamarin
hamadryas baboon
wolf
four-toed hedgehog
mandrill
polar bear
wild boar
common raccoon
harbor seal
emperor tamarin
common warthog
ring-tailed lemur
silvery marmoset
nine-banded armadillo
white-faced saki monkey
pygmy marmoset
gorilla
common marmoset
panther
capybara
brown rat
red panda
koala
mantled guereza
Arabian oryx
sheep
addax
californian sea lion
degu
reindeer
black-tailed prairie dog
fennec
tammar wallaby
Parma wallaby
meerkat
gerenuk
red kangaroo
black-footed ferret
impala
dama gazelle
Speke's gazelle
Thomson's gazelle

Supp.Fig.2A

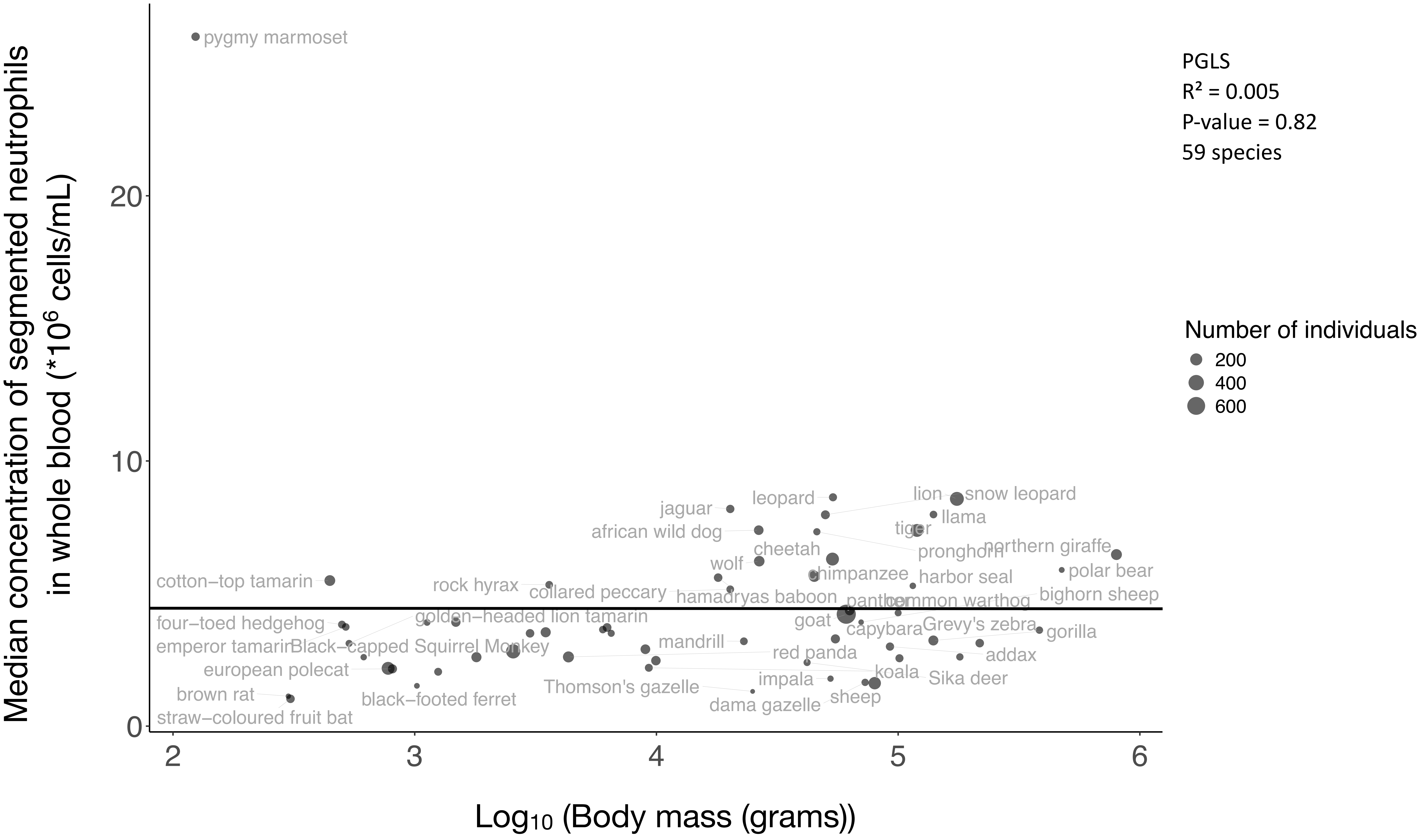

Median concentration of segmented neutrophils in whole blood (*10^6 cells/mL)
Log10 (Body mass (grams))
0
10
20
2
3
4
5
6
pygmy marmoset
cotton−top tamarin
four−toed hedgehog
emperor tamarin
Black−capped Squirrel Monkey
european polecat
brown rat
straw−coloured fruit bat
black−footed ferret
golden−headed lion tamarin
rock hyrax
collared peccary
Thomson's gazelle
mandrill
jaguar
african wild dog
wolf
leopard
cheetah
chimpanzee
hamadryas baboon
goat
red panda
impala
dama gazelle
sheep
koala
panther
capybara
lion
snow leopard
llama
tiger
pronghorn
harbor seal
common warthog
Grevy's zebra
Sika deer
addax
northern giraffe
polar bear
bighorn sheep
gorilla
PGLS
R² = 0.005
P-value = 0.82
59 species
Number of individuals
200
400
600

Supp.Fig.2B

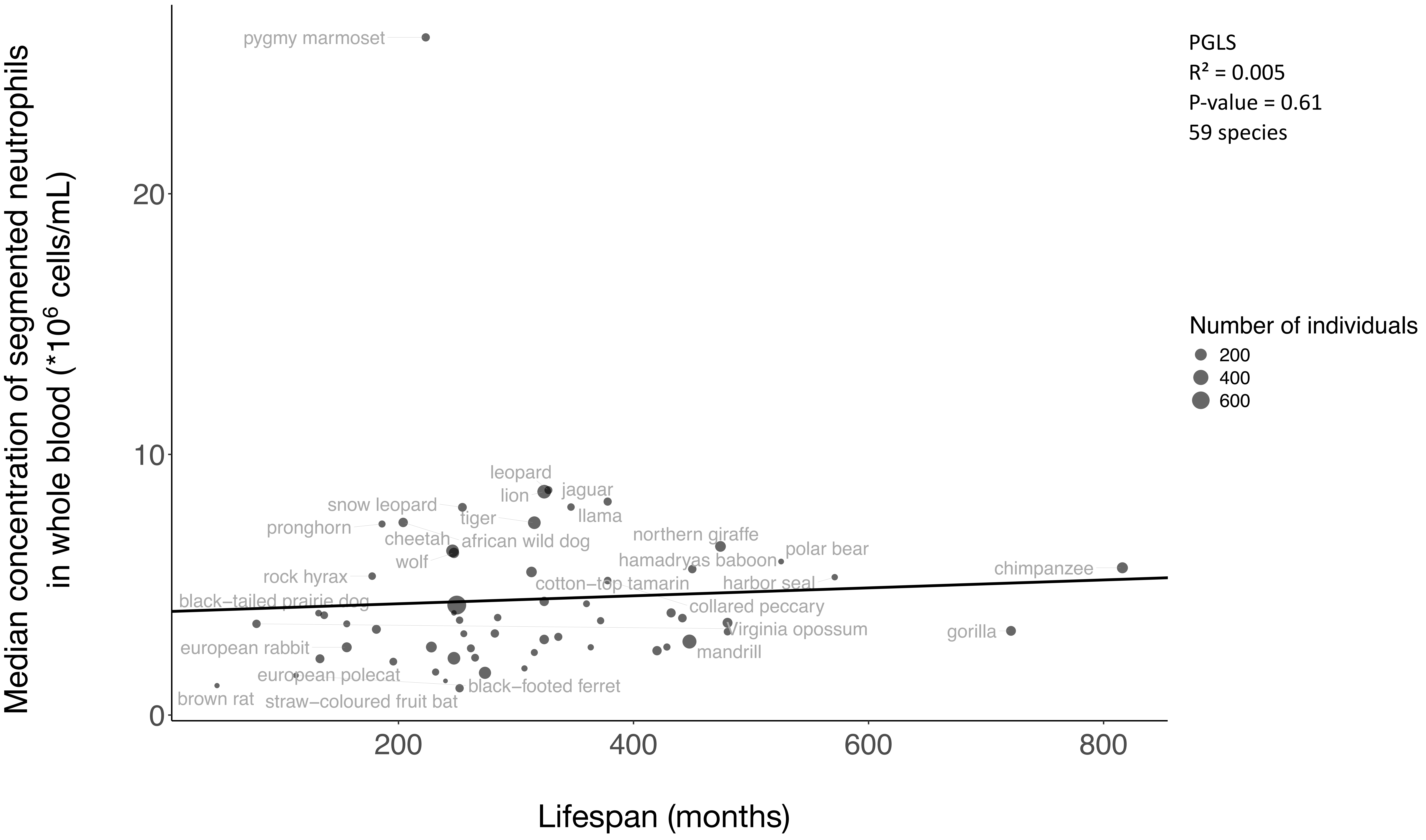

PGLS
R² = 0.005
P-value = 0.61
59 species
Number of individuals
200
400
600
Median concentration of segmented neutrophils in whole blood (*10^6 cells/mL)
Lifespan (months)
0
10
20
200
400
600
800
pygmy marmoset
leopard
lion
jaguar
snow leopard
llama
tiger
pronghorn
cheetah
african wild dog
wolf
northern giraffe
polar bear
hamadryas baboon
rock hyrax
chimpanzee
cotton−top tamarin
harbor seal
black−tailed prairie dog
collared peccary
Virginia opossum
gorilla
european rabbit
mandrill
european polecat
black−footed ferret
brown rat
straw−coloured fruit bat

Supp.Fig.3A

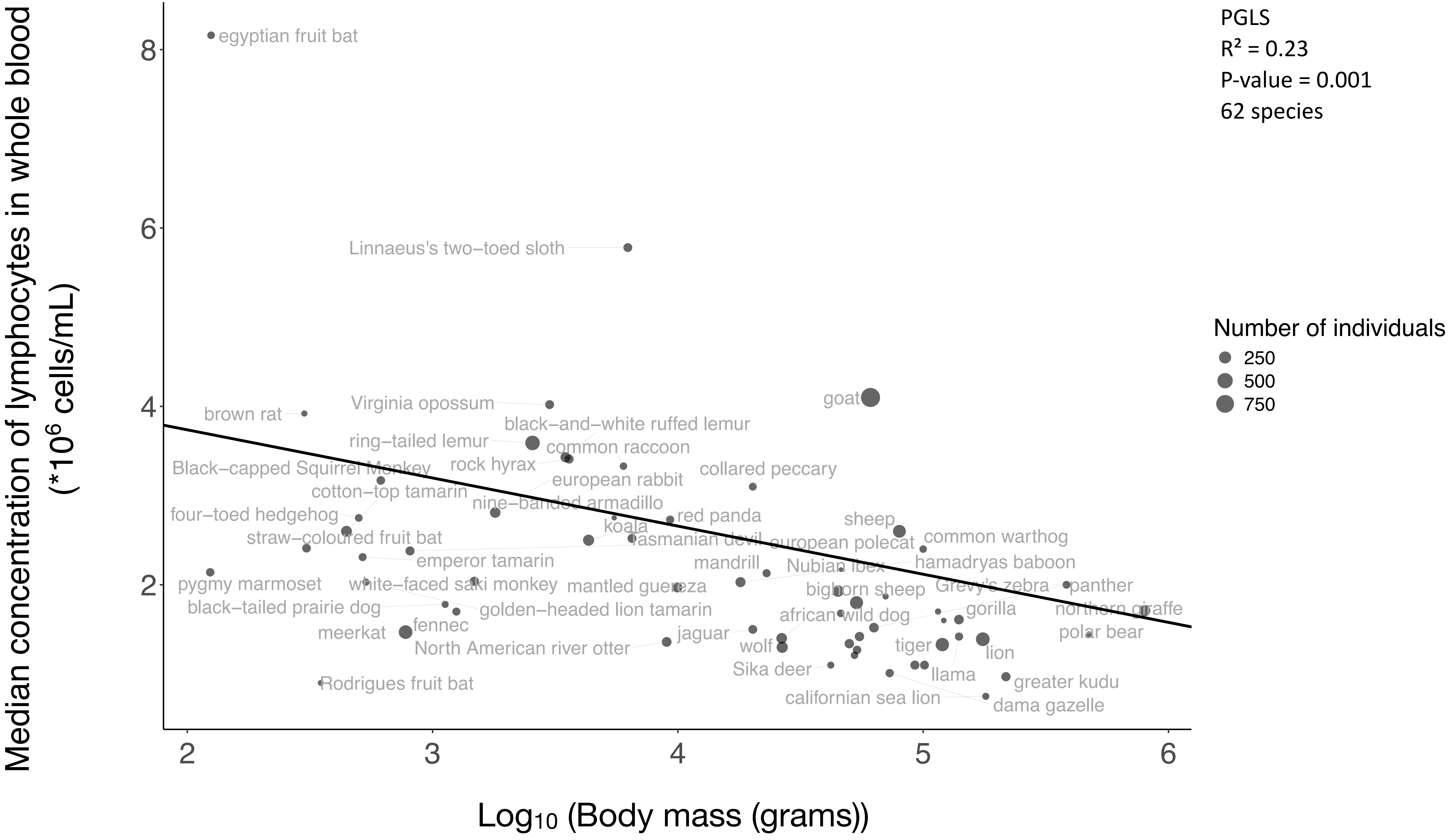

PGLS
R² = 0.23
P-value = 0.001
62 species
Number of individuals
250
500
750
Median concentration of lymphocytes in whole blood (*10^6 cells/mL)
Log10 (Body mass (grams))
8
6
4
2
2
3
4
5
6
egyptian fruit bat
Linnaeus's two−toed sloth
Virginia opossum
brown rat
goat
black−and−white ruffed lemur
ring−tailed lemur
common raccoon
rock hyrax
Black−capped Squirrel Monkey
collared peccary
european rabbit
cotton−top tamarin
nine−banded armadillo
four−toed hedgehog
red panda
koala
sheep
straw−coloured fruit bat
tasmanian devil
european polecat
common warthog
emperor tamarin
mandrill
Nubian ibex
hamadryas baboon
pygmy marmoset
white−faced saki monkey
mantled guereza
bighorn sheep
Grevy's zebra
panther
black−tailed prairie dog
golden−headed lion tamarin
african wild dog
gorilla
northern giraffe
fennec
meerkat
jaguar
polar bear
North American river otter
wolf
tiger
lion
Sika deer
llama
Rodrigues fruit bat
greater kudu
californian sea lion
dama gazelle

Supp.Fig.3B

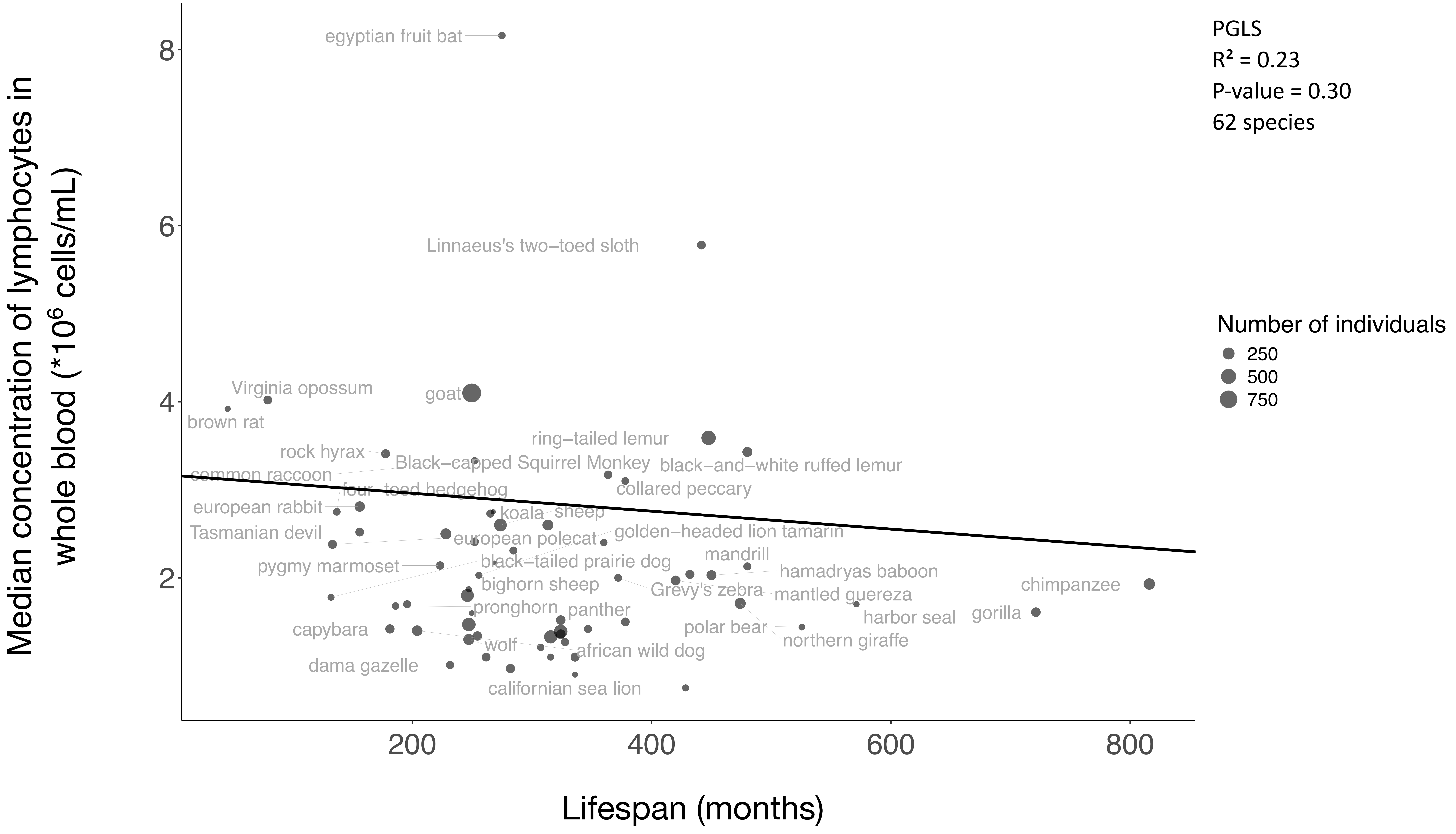

PGLS
R² = 0.23
P-value = 0.30
62 species
Number of individuals
250
500
750
Median concentration of lymphocytes in whole blood (*10^6 cells/mL)
Lifespan (months)
egyptian fruit bat
Linnaeus's two−toed sloth
Virginia opossum
goat
brown rat
ring−tailed lemur
rock hyrax
Black−capped Squirrel Monkey
black−and−white ruffed lemur
common raccoon
collared peccary
four−toed hedgehog
european rabbit
koala
sheep
Tasmanian devil
golden−headed lion tamarin
european polecat
mandrill
black−tailed prairie dog
pygmy marmoset
hamadryas baboon
bighorn sheep
Grevy's zebra
mantled guereza
chimpanzee
pronghorn
panther
harbor seal
gorilla
capybara
polar bear
northern giraffe
wolf
african wild dog
dama gazelle
californian sea lion

Supp.Fig.4A

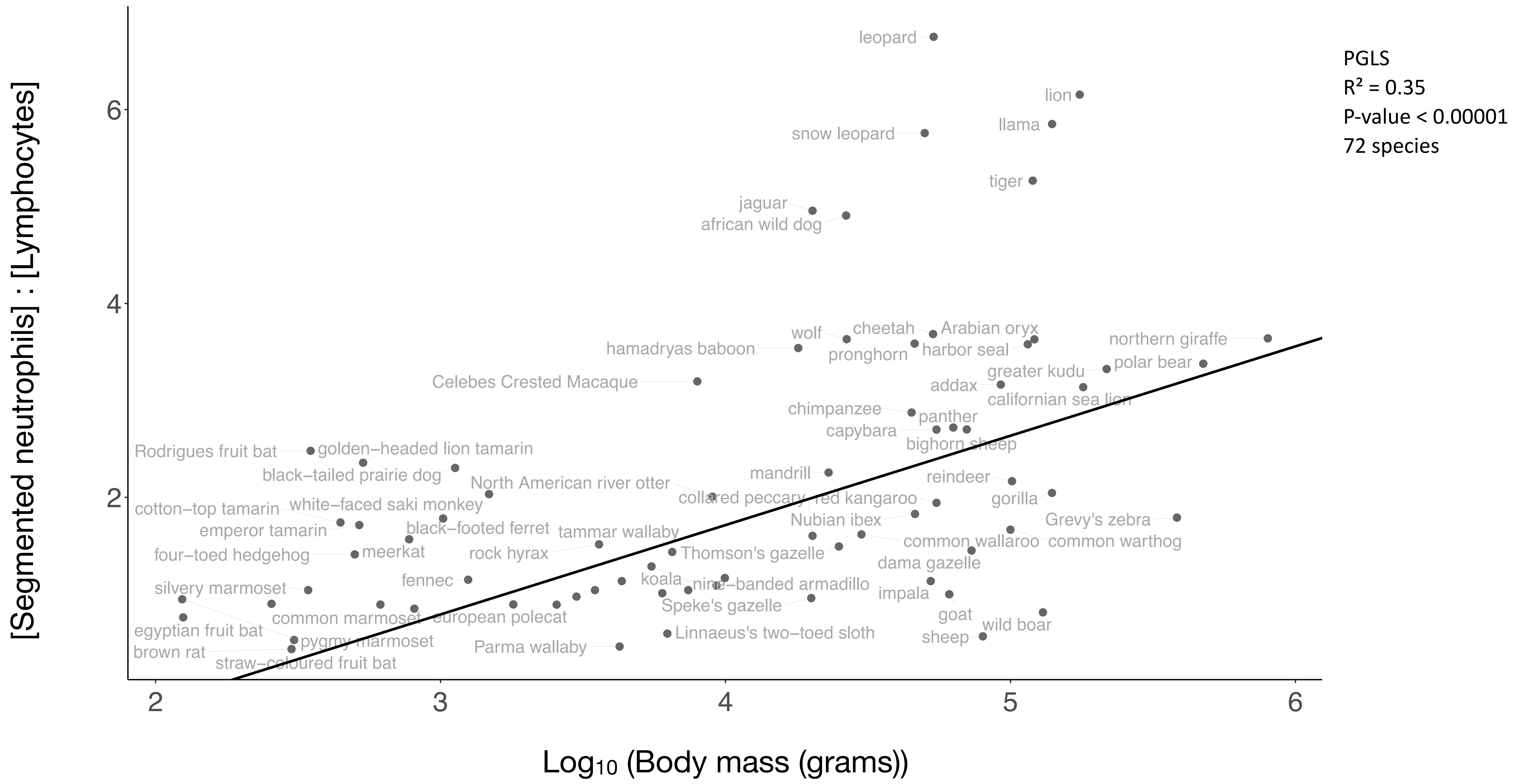
PGLS
R² = 0.35
P-value < 0.00001
72 species
[Segmented neutrophils] : [Lymphocytes]
Log10 (Body mass (grams))
2
4
6
2
3
4
5
6
leopard
lion
llama
snow leopard
tiger
jaguar
african wild dog
wolf
cheetah
Arabian oryx
hamadryas baboon
pronghorn
harbor seal
northern giraffe
polar bear
greater kudu
Celebes Crested Macaque
addax
californian sea lion
chimpanzee
panther
capybara
bighorn sheep
Rodrigues fruit bat
golden–headed lion tamarin
black–tailed prairie dog
mandrill
North American river otter
reindeer
collared peccary
red kangaroo
gorilla
cotton–top tamarin
white–faced saki monkey
Nubian ibex
Grevy's zebra
emperor tamarin
black–footed ferret
tammar wallaby
common wallaroo
common warthog
four–toed hedgehog
meerkat
rock hyrax
Thomson's gazelle
dama gazelle
fennec
koala
nine–banded armadillo
silvery marmoset
impala
Speke's gazelle
goat
common marmoset
european polecat
wild boar
egyptian fruit bat
Linnaeus's two–toed sloth
sheep
pygmy marmoset
brown rat
Parma wallaby
straw–coloured fruit bat

Supp.Fig.4B

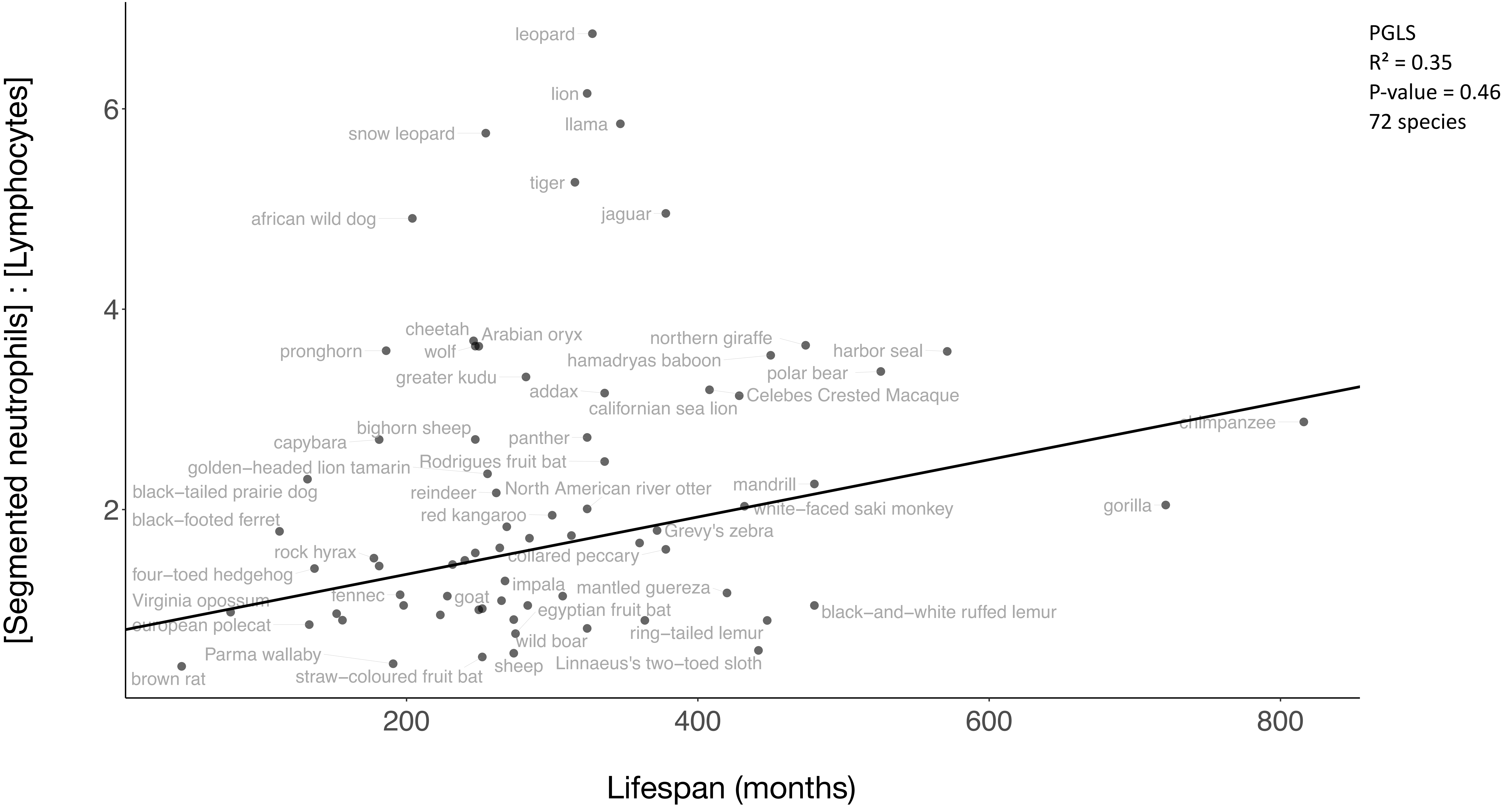
PGLS
R² = 0.35
P-value = 0.46
72 species
[Segmented neutrophils] : [Lymphocytes]
Lifespan (months)
2
4
6
200
400
600
800
leopard
lion
llama
snow leopard
tiger
jaguar
african wild dog
cheetah
Arabian oryx
wolf
pronghorn
northern giraffe
harbor seal
hamadryas baboon
polar bear
greater kudu
addax
Celebes Crested Macaque
californian sea lion
chimpanzee
bighorn sheep
panther
capybara
Rodrigues fruit bat
golden−headed lion tamarin
black−tailed prairie dog
reindeer
North American river otter
mandrill
white−faced saki monkey
gorilla
red kangaroo
black−footed ferret
Grevy's zebra
rock hyrax
collared peccary
four−toed hedgehog
impala
mantled guereza
Virginia opossum
fennec
goat
egyptian fruit bat
black−and−white ruffed lemur
european polecat
ring−tailed lemur
wild boar
Parma wallaby
Linnaeus's two−toed sloth
sheep
straw−coloured fruit bat
brown rat

Supp.Fig.5A

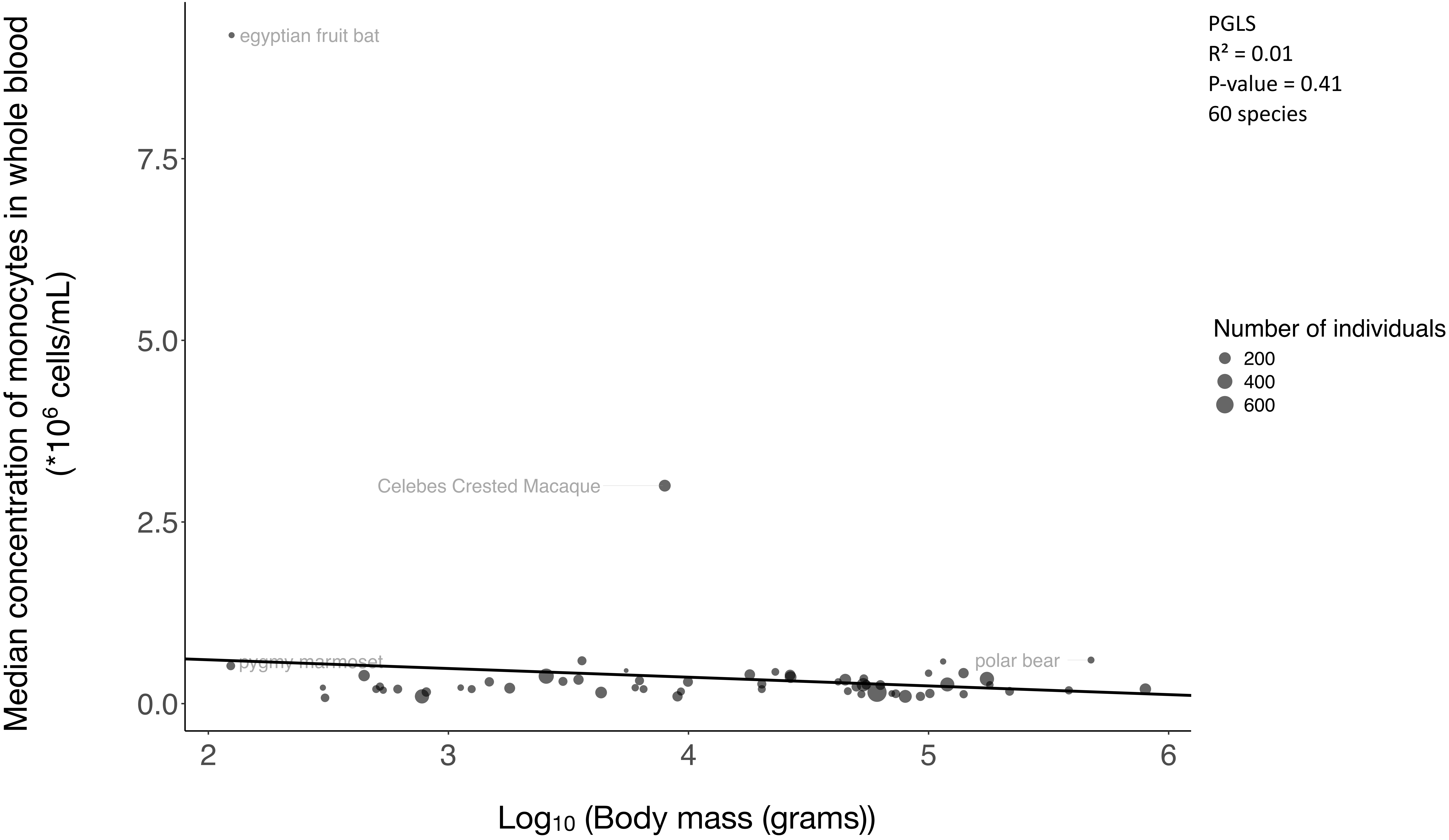

Median concentration of monocytes in whole blood
(*10^6 cells/mL)
7.5
5.0
2.5
0.0
2
3
4
5
6
Log10 (Body mass (grams))
egyptian fruit bat
Celebes Crested Macaque
pygmy marmoset
polar bear
PGLS
R² = 0.01
P-value = 0.41
60 species
Number of individuals
200
400
600

Supp.Fig.5B

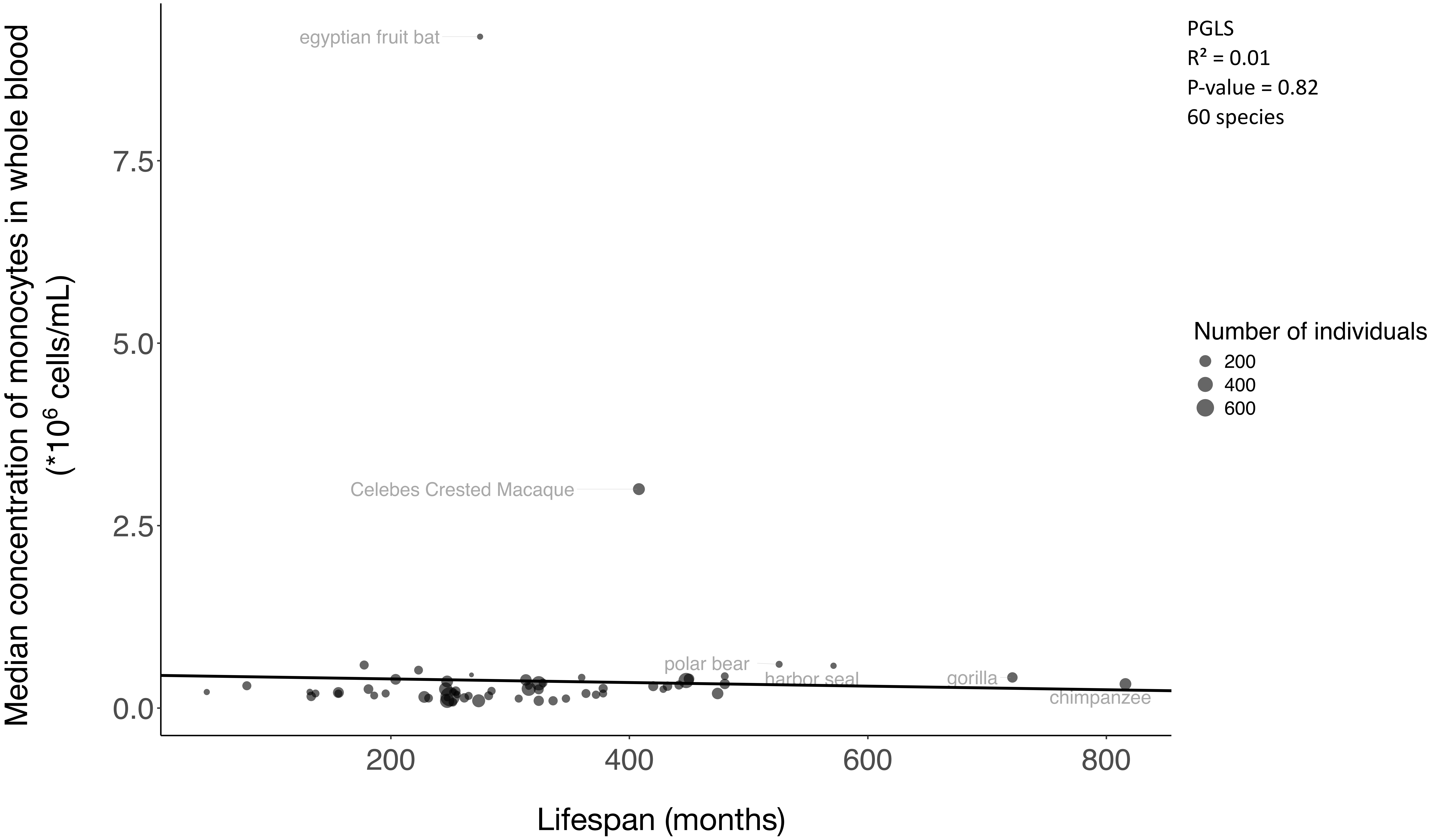
PGLS
R² = 0.01
P-value = 0.82
60 species
Number of individuals
200
400
600
Median concentration of monocytes in whole blood
(*10^6 cells/mL)
7.5
5.0
2.5
0.0
egyptian fruit bat
Celebes Crested Macaque
polar bear
harbor seal
gorilla
chimpanzee
200
400
600
800
Lifespan (months)

Supp.Fig.6A

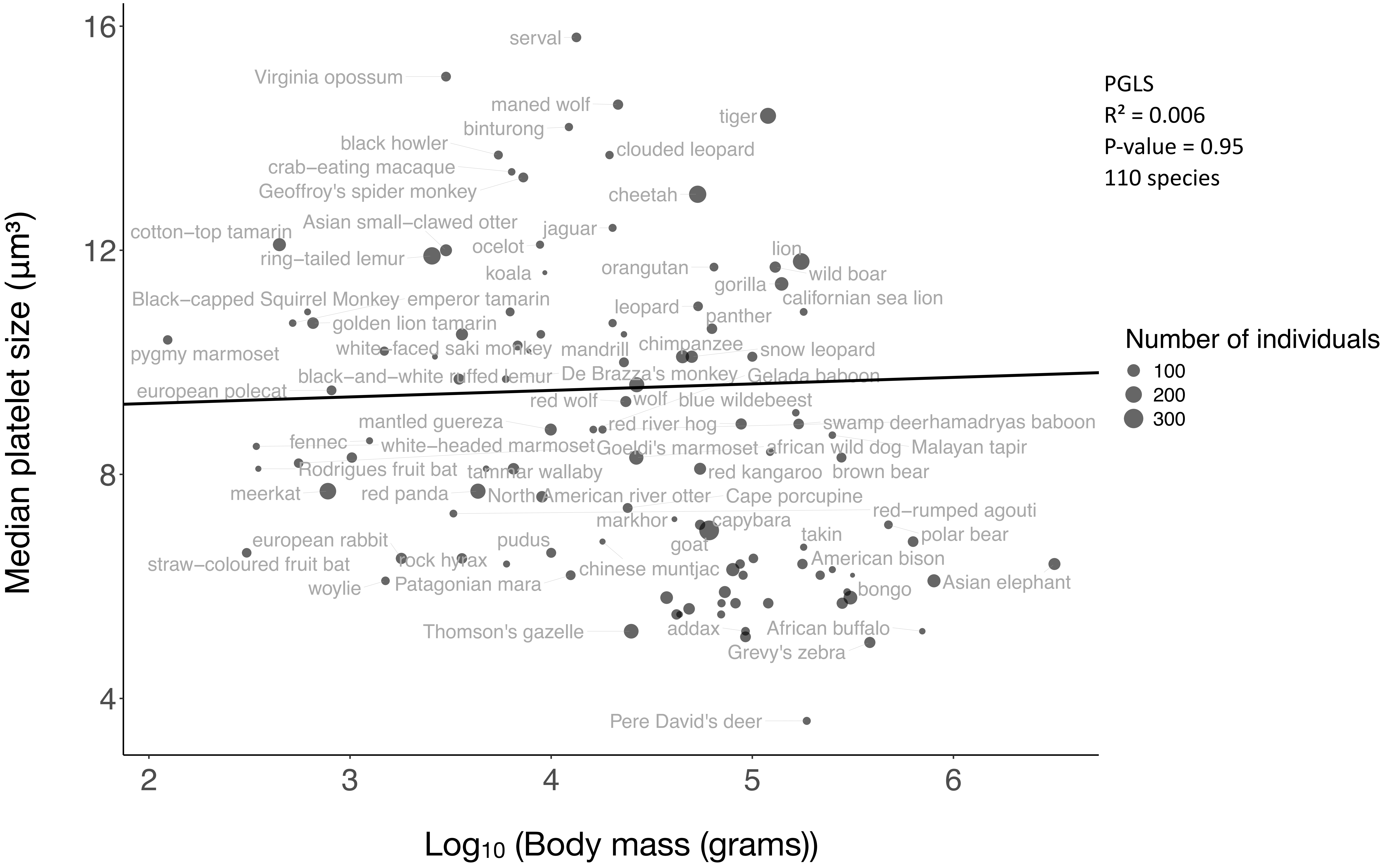

Median platelet size (µm³)
Log10 (Body mass (grams))
16
12
8
4
2
3
4
5
6
PGLS
R² = 0.006
P-value = 0.95
110 species
Number of individuals
100
200
300
serval
Virginia opossum
maned wolf
tiger
binturong
black howler
clouded leopard
crab–eating macaque
Geoffroy's spider monkey
cheetah
cotton–top tamarin
Asian small–clawed otter
jaguar
ocelot
ring–tailed lemur
koala
orangutan
lion
wild boar
gorilla
Black–capped Squirrel Monkey
emperor tamarin
californian sea lion
leopard
panther
golden lion tamarin
chimpanzee
white–faced saki monkey
mandrill
snow leopard
pygmy marmoset
black–and–white ruffed lemur
De Brazza's monkey
Gelada baboon
european polecat
red wolf
wolf
blue wildebeest
mantled guereza
red river hog
swamp deer
hamadryas baboon
fennec
white–headed marmoset
Goeldi's marmoset
african wild dog
Malayan tapir
Rodrigues fruit bat
tammar wallaby
red kangaroo
brown bear
meerkat
red panda
North American river otter
Cape porcupine
red–rumped agouti
markhor
capybara
european rabbit
pudus
goat
takin
polar bear
straw–coloured fruit bat
rock hyrax
chinese muntjac
American bison
woylie
Patagonian mara
Asian elephant
bongo
Thomson's gazelle
addax
African buffalo
Grevy's zebra
Pere David's deer

Supp.Fig.6B

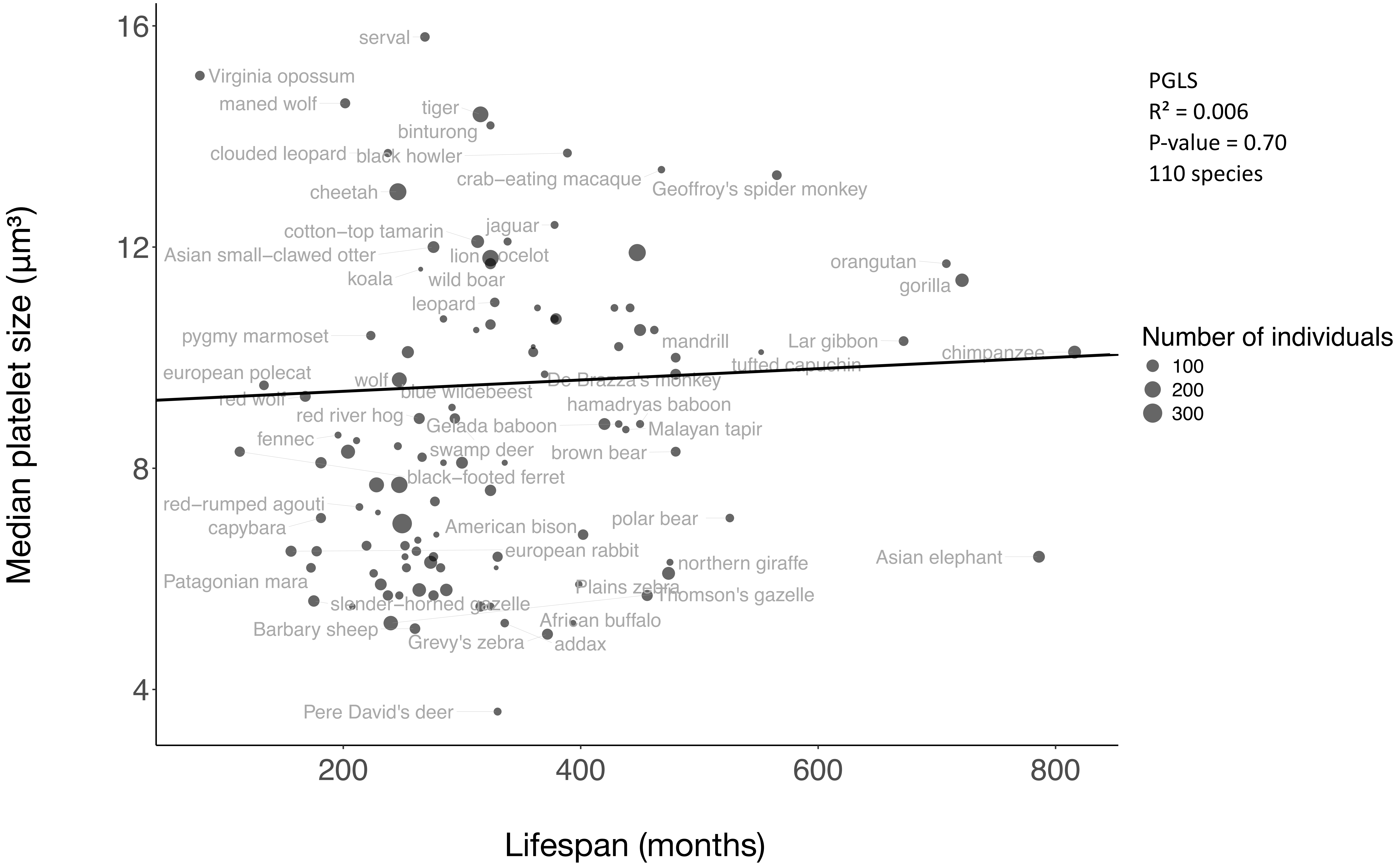
PGLS
R² = 0.006
P-value = 0.70
110 species
Number of individuals
100
200
300
Median platelet size (µm³)
Lifespan (months)
16
12
8
4
200
400
600
800
serval
Virginia opossum
maned wolf
tiger
binturong
clouded leopard
black howler
crab−eating macaque
Geoffroy's spider monkey
cheetah
jaguar
cotton−top tamarin
Asian small−clawed otter
lion
ocelot
koala
wild boar
orangutan
gorilla
leopard
pygmy marmoset
mandrill
Lar gibbon
chimpanzee
tufted capuchin
european polecat
wolf
De Brazza's monkey
red wolf
blue wildebeest
hamadryas baboon
red river hog
Gelada baboon
Malayan tapir
fennec
swamp deer
brown bear
black−footed ferret
red−rumped agouti
polar bear
capybara
American bison
european rabbit
Asian elephant
northern giraffe
Patagonian mara
Plains zebra
Thomson's gazelle
slender−horned gazelle
African buffalo
Barbary sheep
Grevy's zebra
addax
Pere David's deer

Supp.Fig.7A

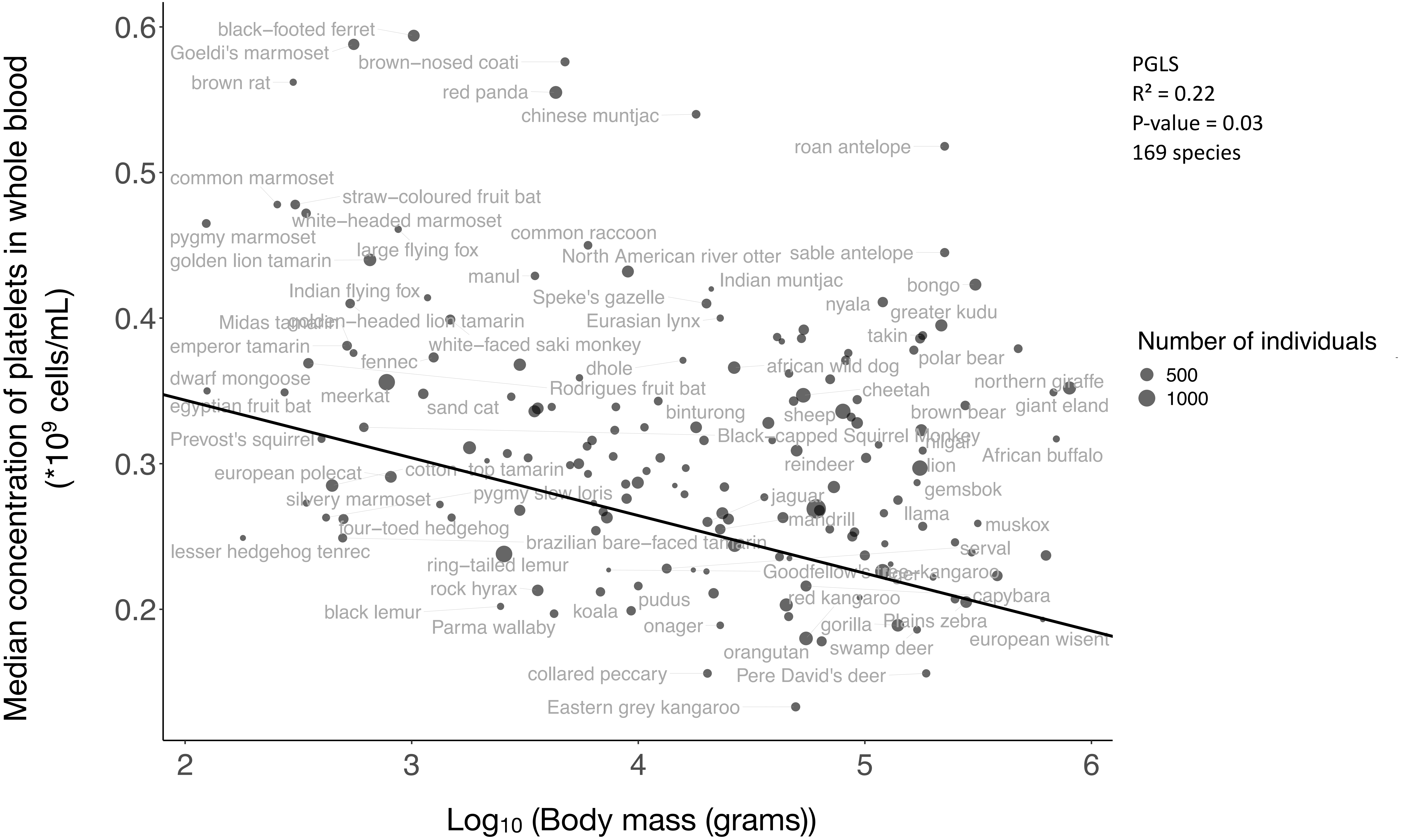
Median concentration of platelets in whole blood
(*10⁹ cells/mL)
Log10 (Body mass (grams))
PGLS
R² = 0.22
P-value = 0.03
169 species
Number of individuals
500
1000
black−footed ferret
Goeldi's marmoset
brown−nosed coati
brown rat
red panda
chinese muntjac
roan antelope
common marmoset
straw−coloured fruit bat
white−headed marmoset
common raccoon
pygmy marmoset
large flying fox
sable antelope
golden lion tamarin
North American river otter
manul
Indian muntjac
Indian flying fox
Speke's gazelle
bongo
nyala
greater kudu
golden−headed lion tamarin
Eurasian lynx
takin
emperor tamarin
white−faced saki monkey
polar bear
fennec
dhole
african wild dog
dwarf mongoose
Rodrigues fruit bat
northern giraffe
meerkat
cheetah
egyptian fruit bat
sand cat
binturong
sheep
brown bear
giant eland
Prevost's squirrel
Black−capped Squirrel Monkey
nilgai
African buffalo
european polecat
cotton−top tamarin
reindeer
lion
silvery marmoset
pygmy slow loris
jaguar
gemsbok
four−toed hedgehog
mandrill
llama
muskox
brazilian bare−faced tamarin
serval
lesser hedgehog tenrec
ring−tailed lemur
rock hyrax
red kangaroo
capybara
pudus
black lemur
koala
gorilla
Plains zebra
Parma wallaby
onager
european wisent
swamp deer
orangutan
collared peccary
Pere David's deer
Eastern grey kangaroo
0.2
0.3
0.4
0.5
0.6
2
3
4
5
6

Supp.Fig.7B

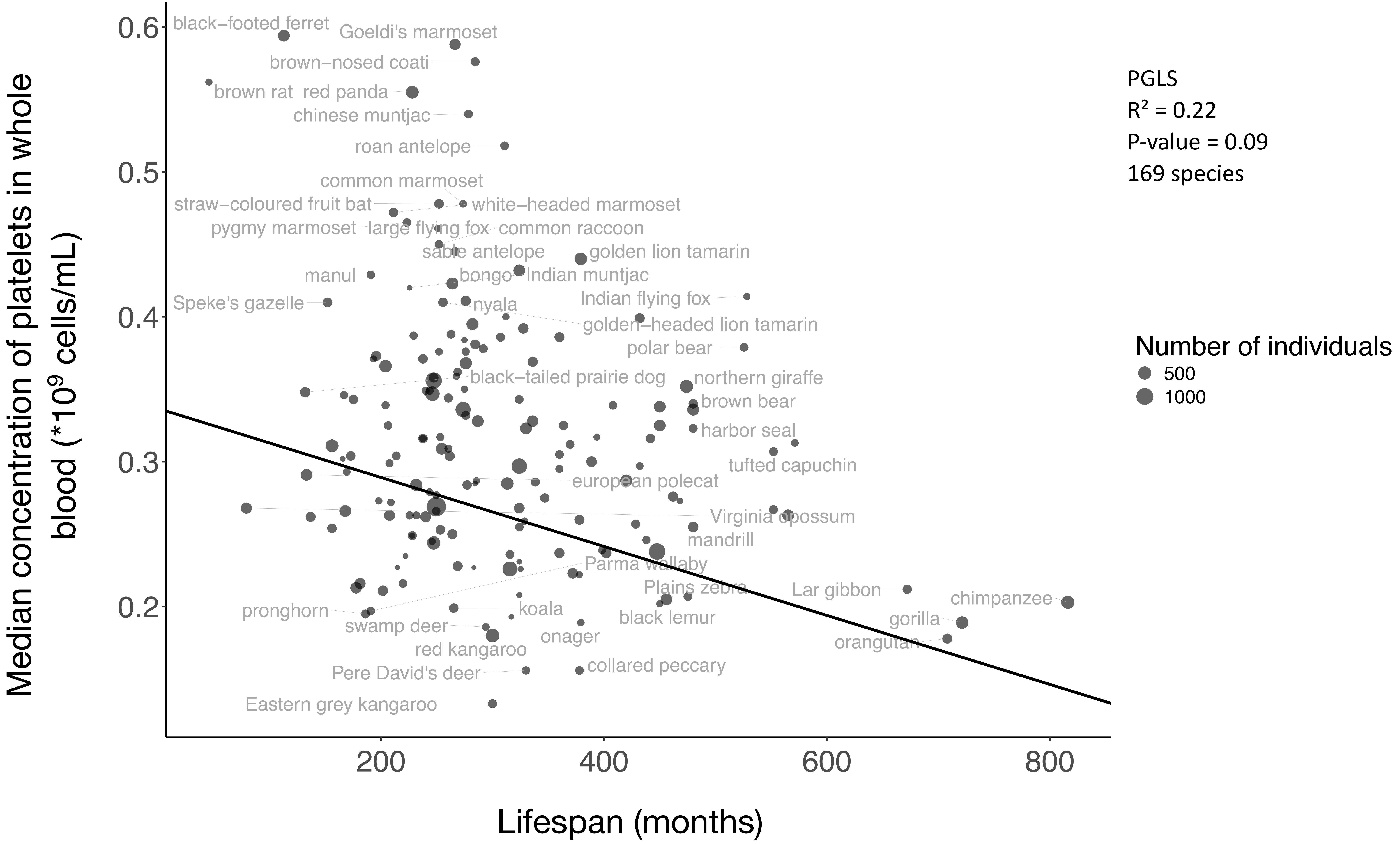

Median concentration of platelets in whole blood (*10^9 cells/mL)
Lifespan (months)
0.6
0.5
0.4
0.3
0.2
200
400
600
800
black−footed ferret
Goeldi's marmoset
brown−nosed coati
brown rat
red panda
chinese muntjac
roan antelope
common marmoset
straw−coloured fruit bat
white−headed marmoset
pygmy marmoset
large flying fox
common raccoon
sable antelope
golden lion tamarin
manul
bongo
Indian muntjac
Speke's gazelle
nyala
Indian flying fox
golden−headed lion tamarin
polar bear
black−tailed prairie dog
northern giraffe
brown bear
harbor seal
tufted capuchin
european polecat
Virginia opossum
mandrill
Parma wallaby
Plains zebra
Lar gibbon
chimpanzee
pronghorn
koala
black lemur
swamp deer
gorilla
onager
red kangaroo
orangutan
collared peccary
Pere David's deer
Eastern grey kangaroo
PGLS
R² = 0.22
P-value = 0.09
169 species
Number of individuals
500
1000

Supp.Fig.8A

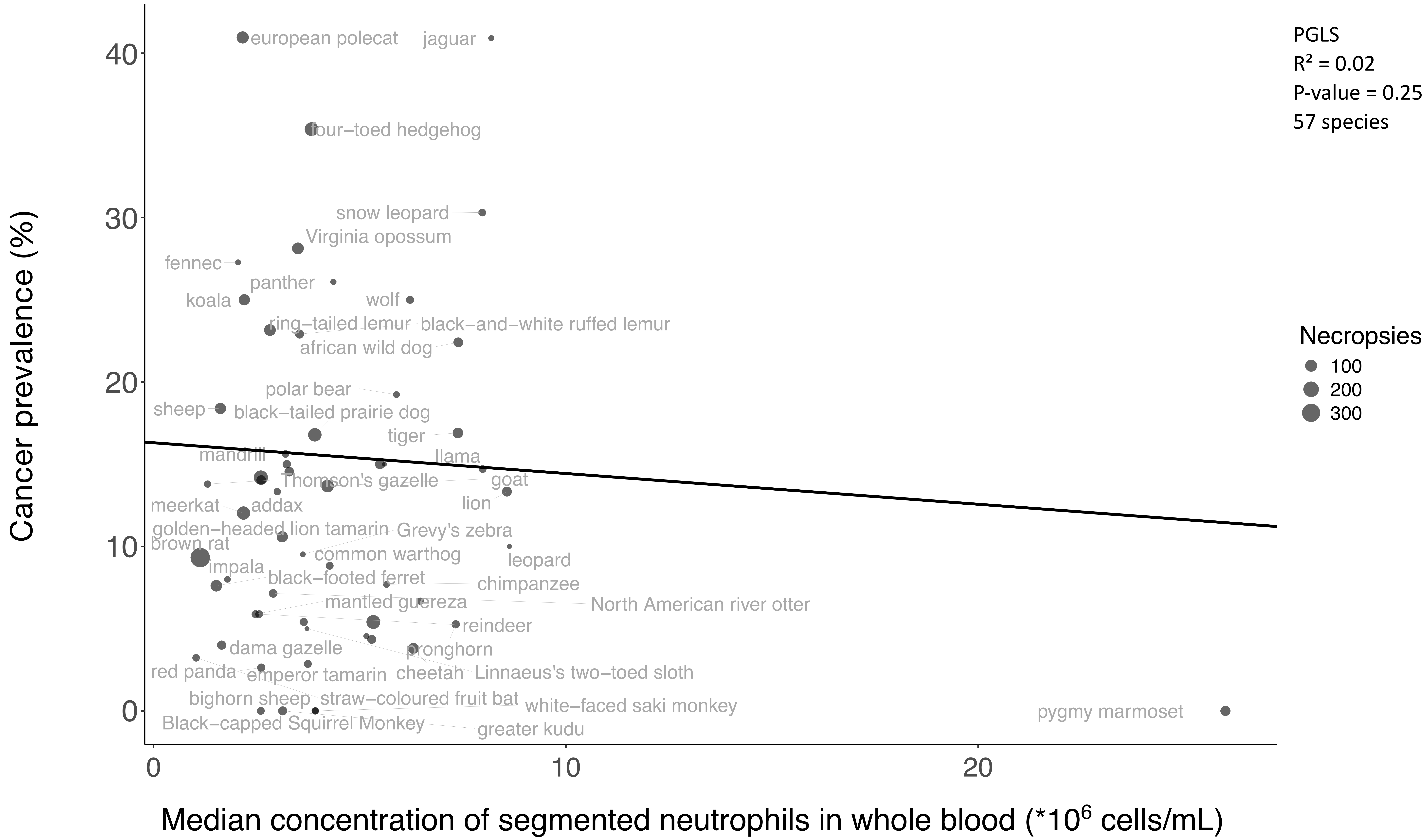
PGLS
R² = 0.02
P-value = 0.25
57 species
Necropsies
100
200
300
Cancer prevalence (%)
Median concentration of segmented neutrophils in whole blood (*10^6 cells/mL)
0
10
20
30
40
european polecat
jaguar
four−toed hedgehog
snow leopard
Virginia opossum
fennec
panther
koala
wolf
ring−tailed lemur
black−and−white ruffed lemur
african wild dog
polar bear
sheep
black−tailed prairie dog
tiger
mandrill
llama
Thomson's gazelle
goat
lion
meerkat
addax
golden−headed lion tamarin
Grevy's zebra
brown rat
common warthog
leopard
impala
black−footed ferret
chimpanzee
mantled guereza
North American river otter
reindeer
dama gazelle
pronghorn
red panda
emperor tamarin
cheetah
Linnaeus's two−toed sloth
bighorn sheep
straw−coloured fruit bat
white−faced saki monkey
pygmy marmoset
Black−capped Squirrel Monkey
greater kudu

Supp.Fig.8B

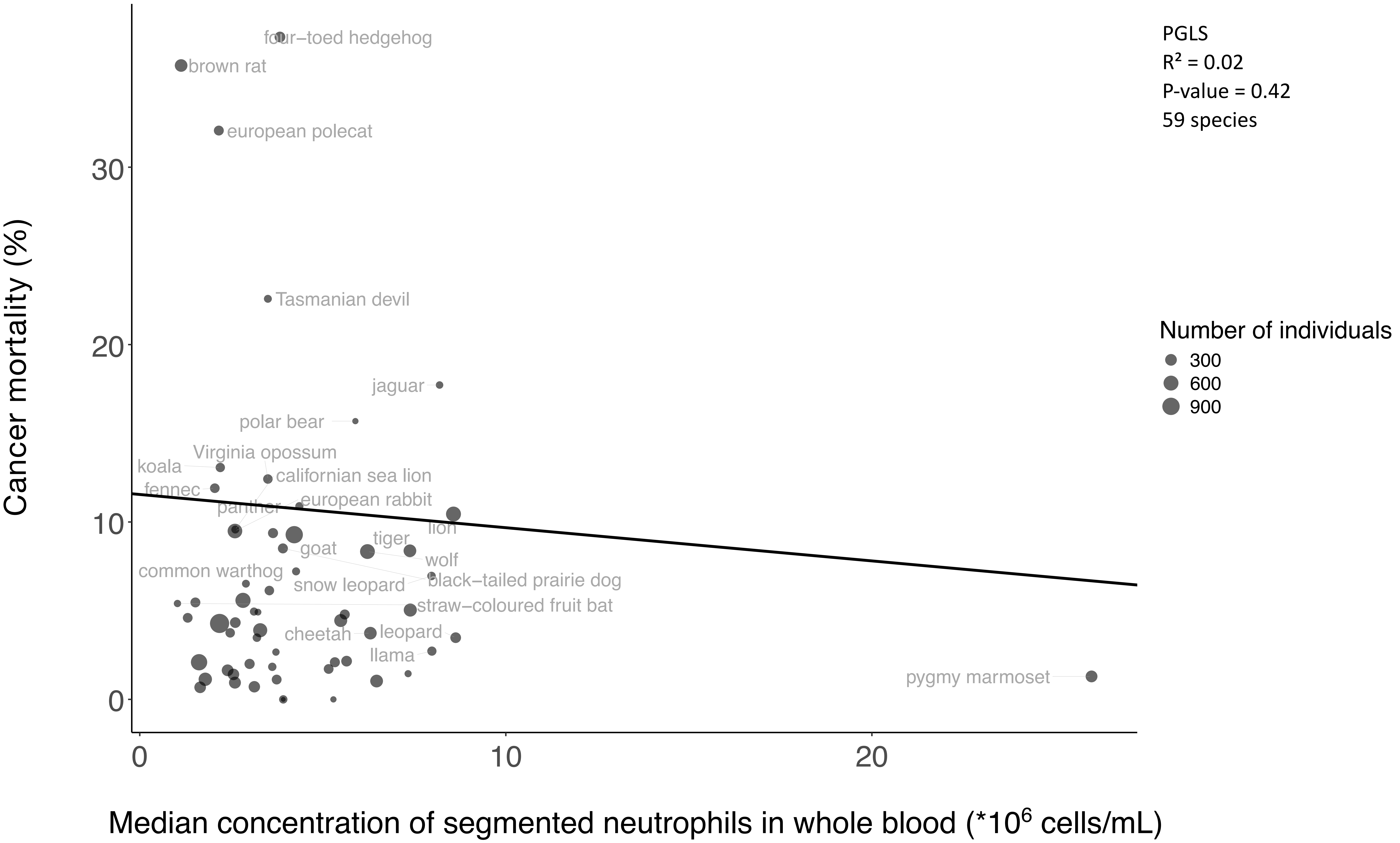

PGLS
R² = 0.02
P-value = 0.42
59 species
Number of individuals
300
600
900
Cancer mortality (%)
Median concentration of segmented neutrophils in whole blood (*10^6 cells/mL)
0
10
20
30
four−toed hedgehog
brown rat
european polecat
Tasmanian devil
jaguar
polar bear
Virginia opossum
koala
californian sea lion
fennec
european rabbit
panther
lion
tiger
goat
wolf
common warthog
black−tailed prairie dog
snow leopard
straw−coloured fruit bat
cheetah
leopard
llama
pygmy marmoset

Supp.Fig.9A

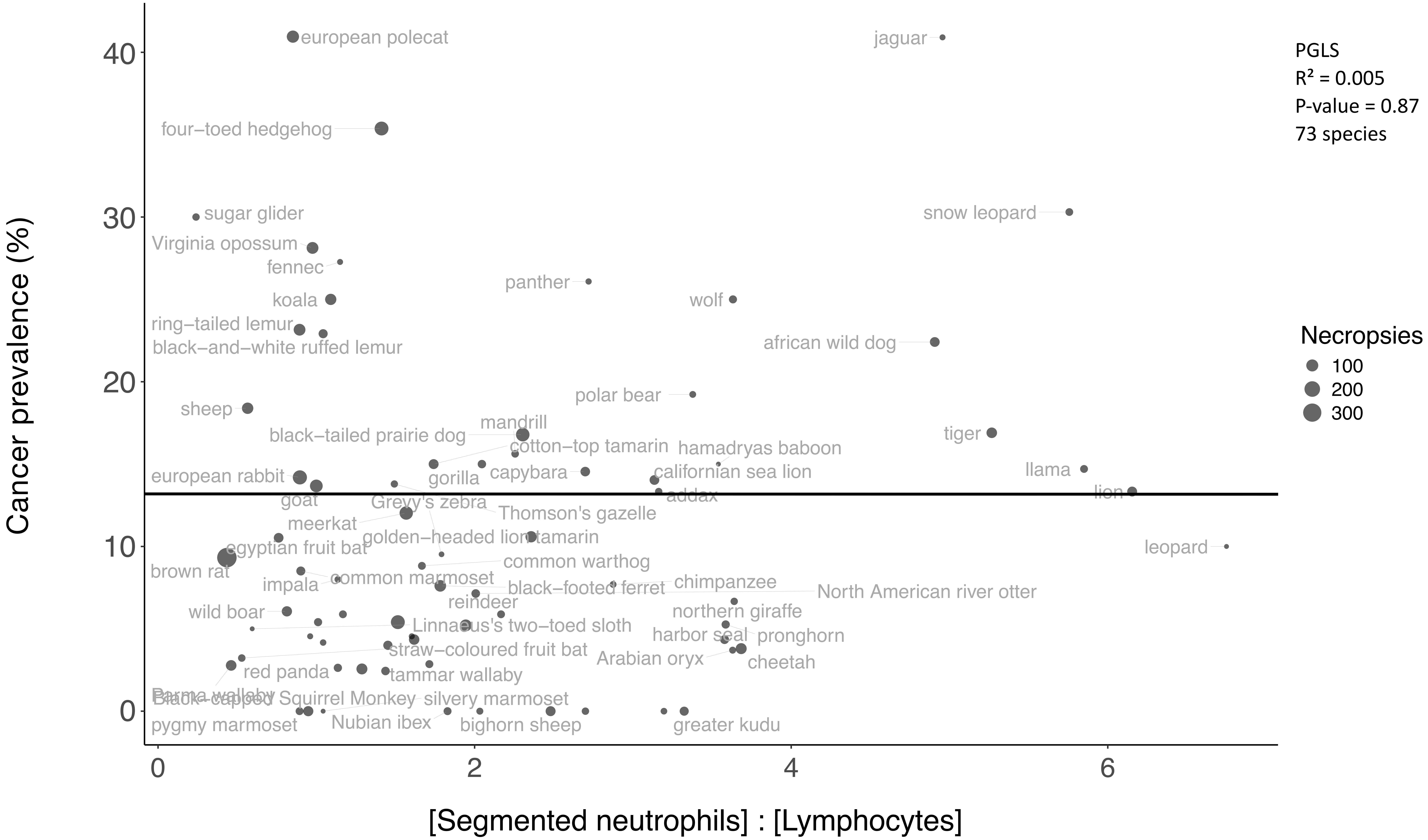
PGLS
R² = 0.005
P-value = 0.87
73 species
Necropsies
100
200
300
Cancer prevalence (%)
[Segmented neutrophils] : [Lymphocytes]
0
10
20
30
40
0
2
4
6
european polecat
jaguar
four−toed hedgehog
sugar glider
snow leopard
Virginia opossum
fennec
panther
koala
wolf
ring−tailed lemur
black−and−white ruffed lemur
african wild dog
polar bear
sheep
mandrill
black−tailed prairie dog
tiger
cotton−top tamarin
hamadryas baboon
capybara
european rabbit
gorilla
californian sea lion
llama
lion
goat
Greyy's zebra
addax
Thomson's gazelle
meerkat
golden−headed lion tamarin
leopard
egyptian fruit bat
common warthog
brown rat
common marmoset
impala
black−footed ferret
chimpanzee
North American river otter
reindeer
wild boar
northern giraffe
Linnaeus's two−toed sloth
harbor seal
pronghorn
straw−coloured fruit bat
Arabian oryx
cheetah
red panda
tammar wallaby
Parma wallaby
Black−capped Squirrel Monkey
silvery marmoset
pygmy marmoset
Nubian ibex
bighorn sheep
greater kudu

Supp.Fig.9B

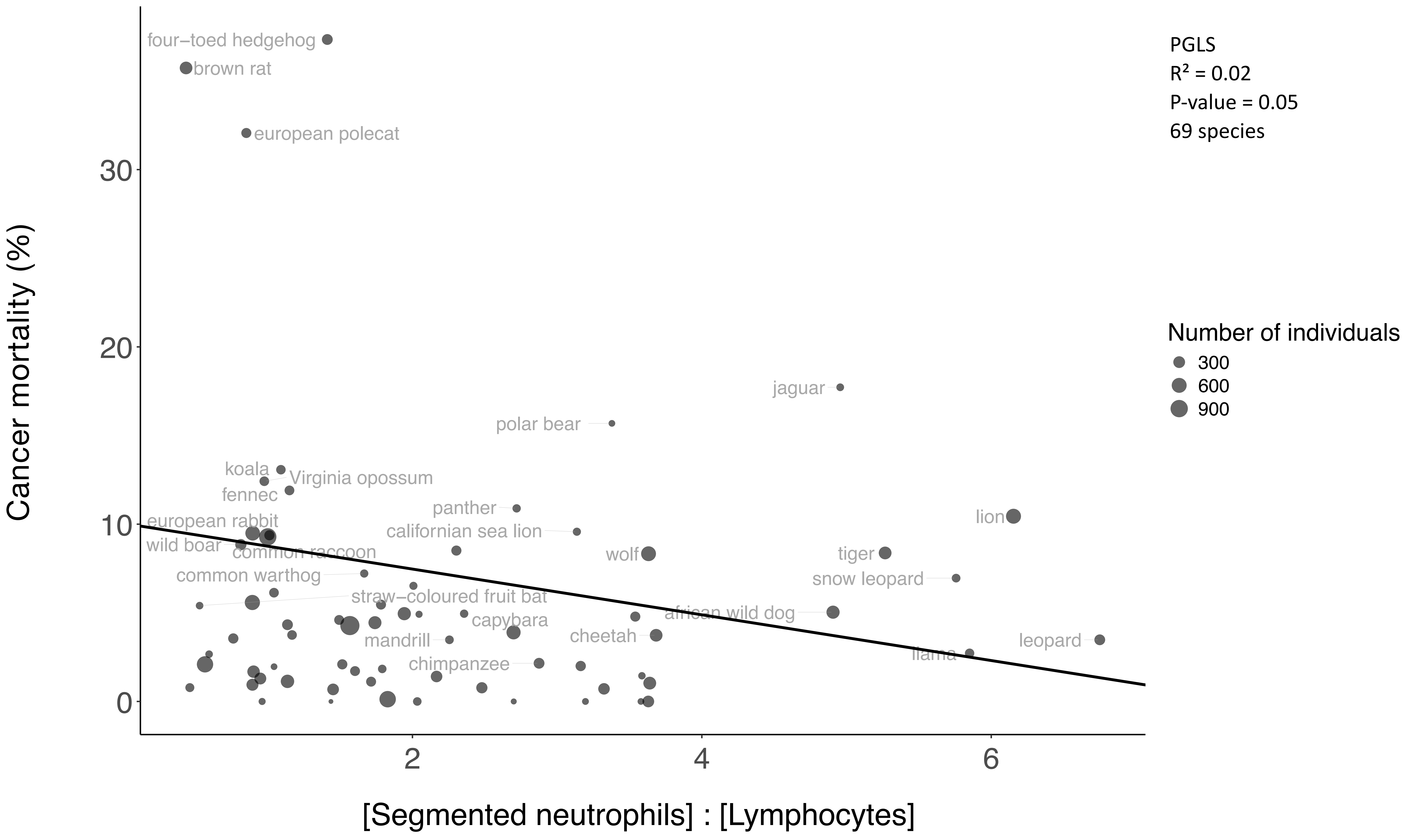

PGLS
R² = 0.02
P-value = 0.05
69 species
Number of individuals
300
600
900
Cancer mortality (%)
[Segmented neutrophils] : [Lymphocytes]
0
10
20
30
2
4
6
four−toed hedgehog
brown rat
european polecat
jaguar
polar bear
koala
Virginia opossum
fennec
panther
european rabbit
californian sea lion
lion
wild boar
common raccoon
wolf
tiger
common warthog
snow leopard
straw−coloured fruit bat
african wild dog
capybara
cheetah
mandrill
leopard
llama
chimpanzee

Supp.Fig.10A

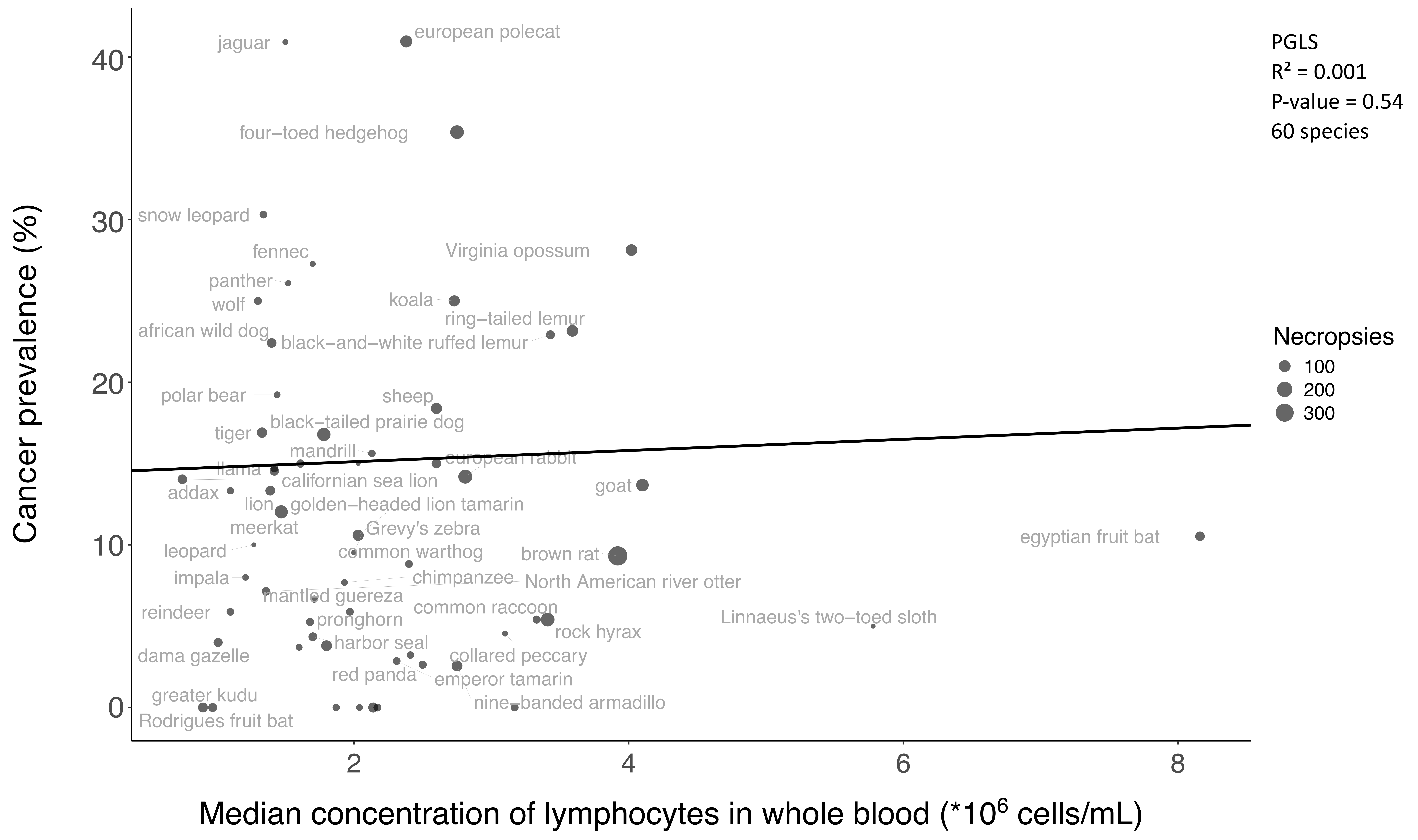
PGLS
R² = 0.001
P-value = 0.54
60 species
Necropsies
100
200
300
Cancer prevalence (%)
Median concentration of lymphocytes in whole blood (*10^6 cells/mL)
0
10
20
30
40
2
4
6
8
jaguar
european polecat
four−toed hedgehog
snow leopard
fennec
Virginia opossum
panther
wolf
koala
ring−tailed lemur
african wild dog
black−and−white ruffed lemur
polar bear
sheep
black−tailed prairie dog
tiger
mandrill
european rabbit
llama
californian sea lion
addax
goat
lion
golden−headed lion tamarin
meerkat
Grevy's zebra
egyptian fruit bat
leopard
common warthog
brown rat
impala
chimpanzee
North American river otter
mantled guereza
reindeer
common raccoon
pronghorn
Linnaeus's two−toed sloth
rock hyrax
harbor seal
dama gazelle
collared peccary
red panda
emperor tamarin
greater kudu
nine−banded armadillo
Rodrigues fruit bat

Supp.Fig.10B

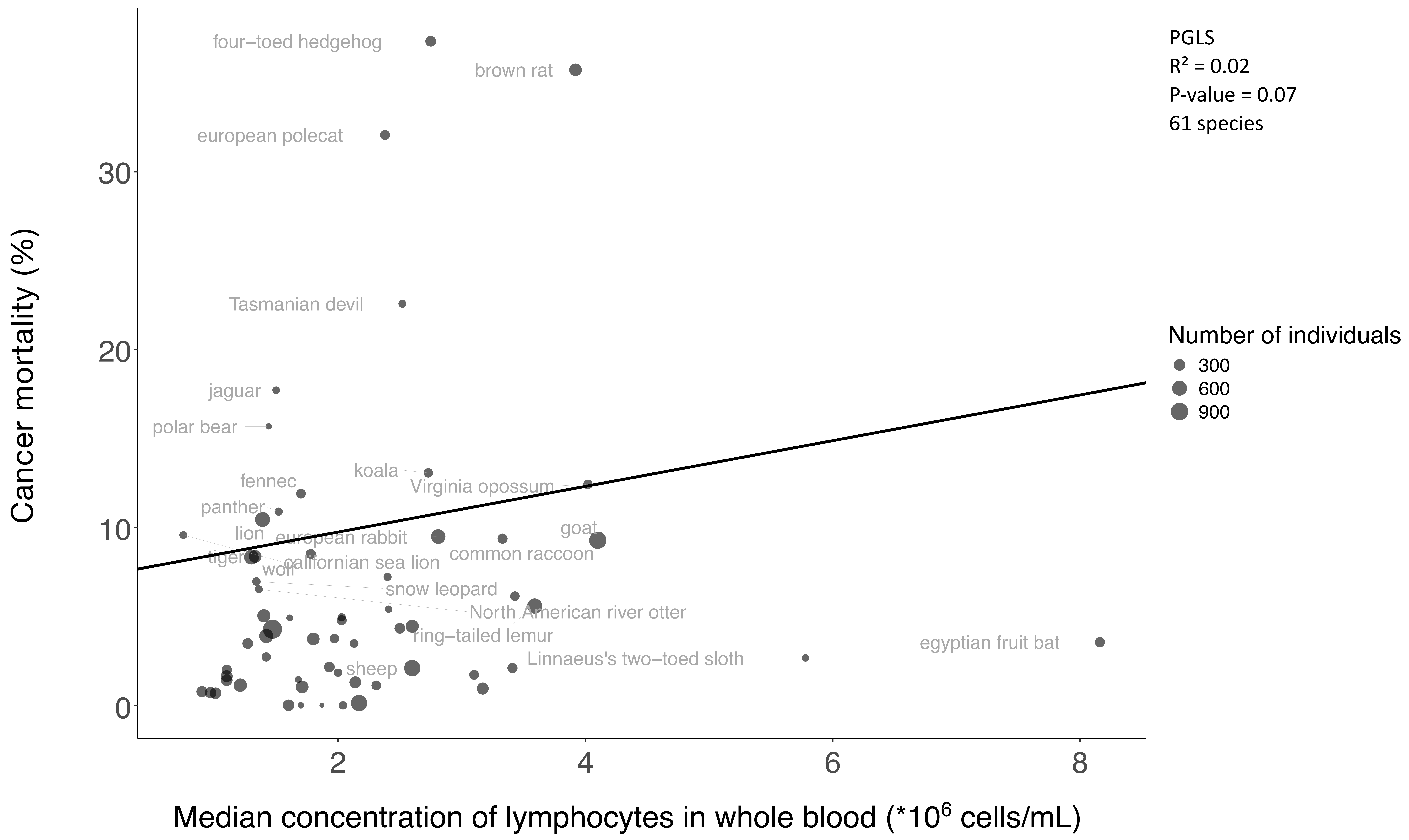

PGLS
R² = 0.02
P-value = 0.07
61 species
Number of individuals
300
600
900
Cancer mortality (%)
Median concentration of lymphocytes in whole blood (*10^6 cells/mL)
0
10
20
30
2
4
6
8
four−toed hedgehog
brown rat
european polecat
Tasmanian devil
jaguar
polar bear
koala
fennec
Virginia opossum
panther
goat
lion
european rabbit
common raccoon
tiger
californian sea lion
wolf
snow leopard
North American river otter
ring−tailed lemur
egyptian fruit bat
Linnaeus's two−toed sloth
sheep

Supp.Fig.11A

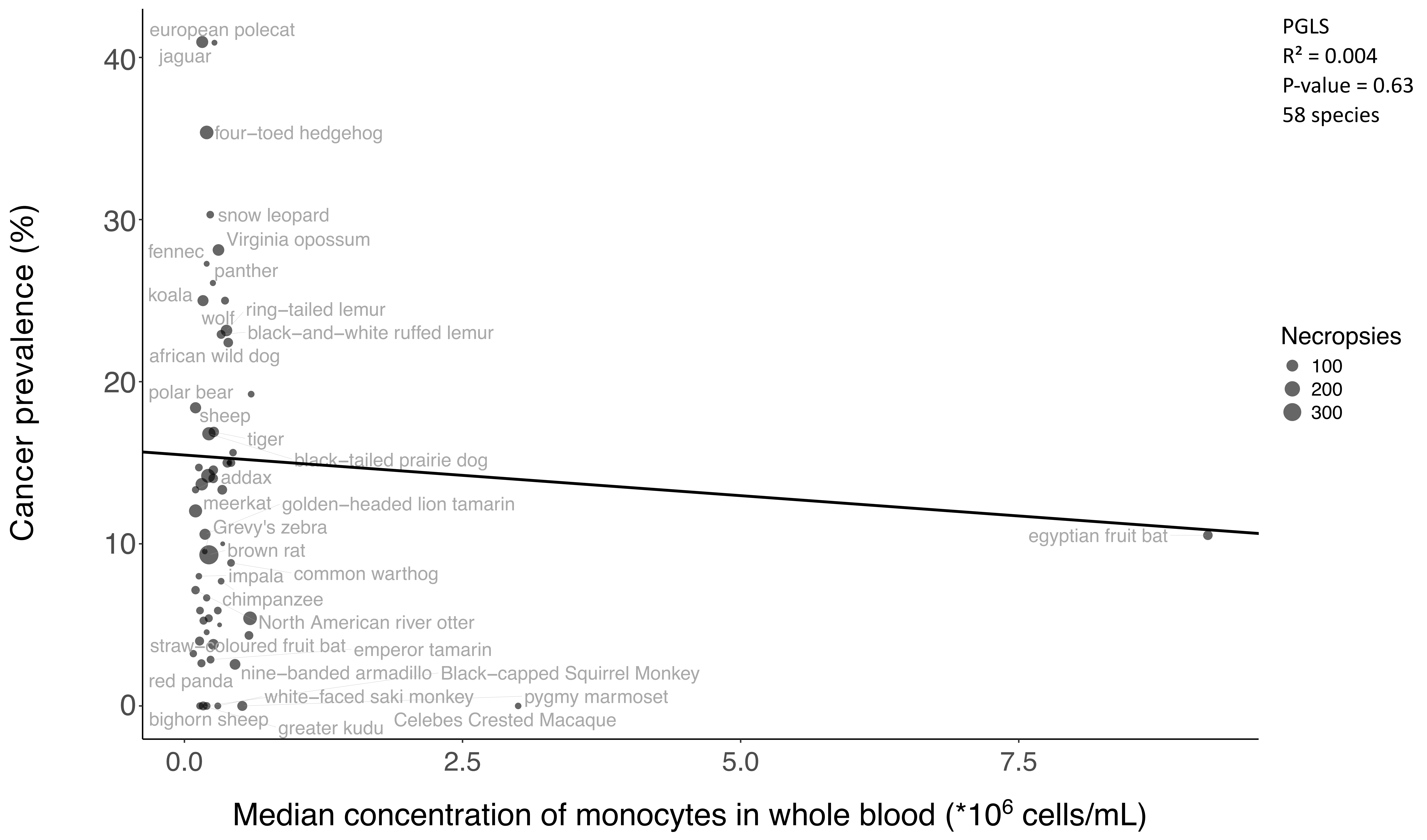
PGLS
R² = 0.004
P-value = 0.63
58 species
Necropsies
100
200
300
Cancer prevalence (%)
Median concentration of monocytes in whole blood (*10^6 cells/mL)
0.0
2.5
5.0
7.5
0
10
20
30
40
european polecat
jaguar
four−toed hedgehog
snow leopard
Virginia opossum
fennec
panther
koala
ring−tailed lemur
wolf
black−and−white ruffed lemur
african wild dog
polar bear
sheep
tiger
black−tailed prairie dog
addax
meerkat
golden−headed lion tamarin
Grevy's zebra
egyptian fruit bat
brown rat
impala
common warthog
chimpanzee
North American river otter
straw−coloured fruit bat
emperor tamarin
nine−banded armadillo
Black−capped Squirrel Monkey
red panda
white−faced saki monkey
pygmy marmoset
bighorn sheep
greater kudu
Celebes Crested Macaque

Supp.Fig.11B

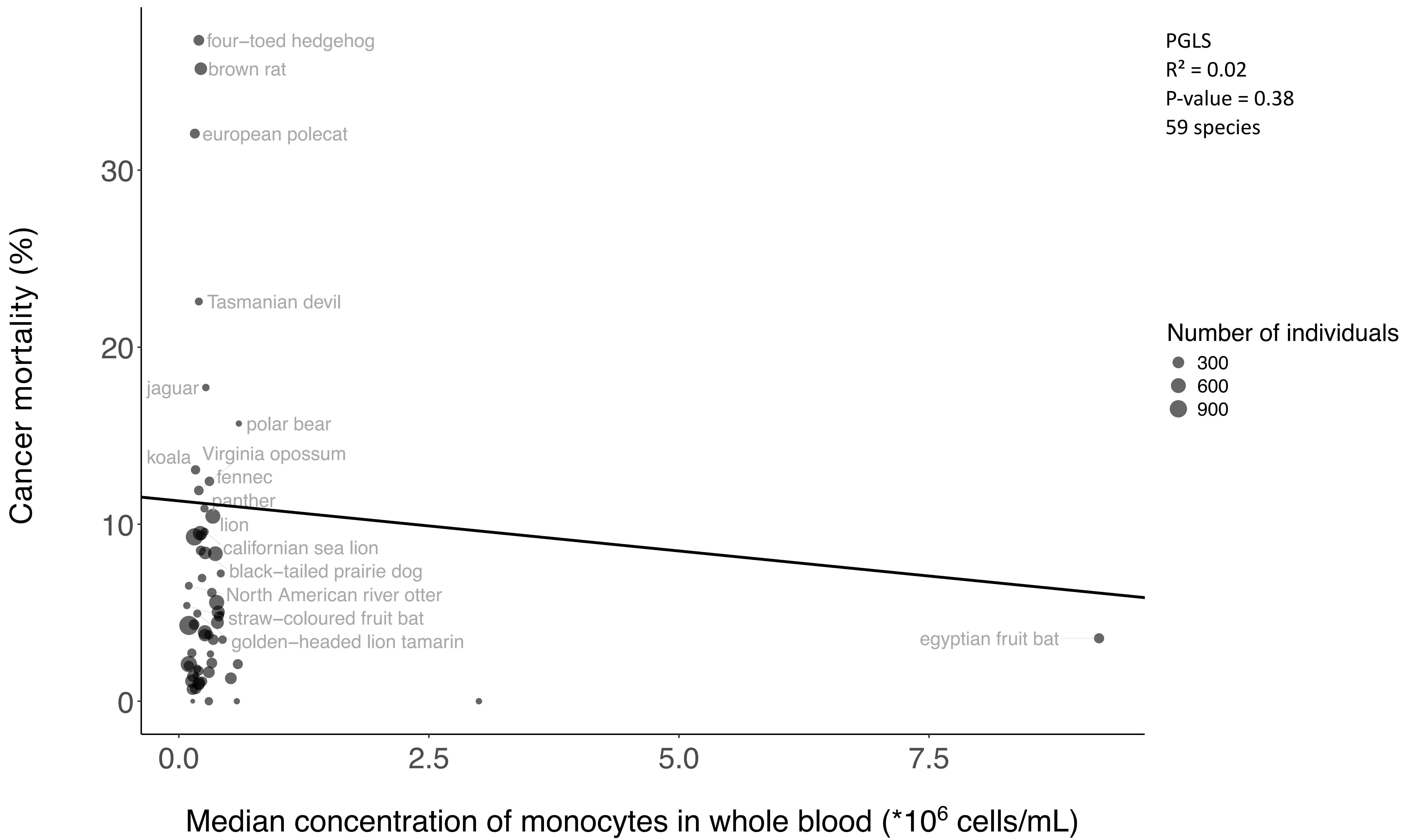
PGLS
R² = 0.02
P-value = 0.38
59 species
Number of individuals
300
600
900
Cancer mortality (%)
Median concentration of monocytes in whole blood (*10^6 cells/mL)
0
10
20
30
0.0
2.5
5.0
7.5
four–toed hedgehog
brown rat
european polecat
Tasmanian devil
jaguar
polar bear
koala
Virginia opossum
fennec
panther
lion
californian sea lion
black–tailed prairie dog
North American river otter
straw–coloured fruit bat
golden–headed lion tamarin
egyptian fruit bat

Supp.Fig.12A

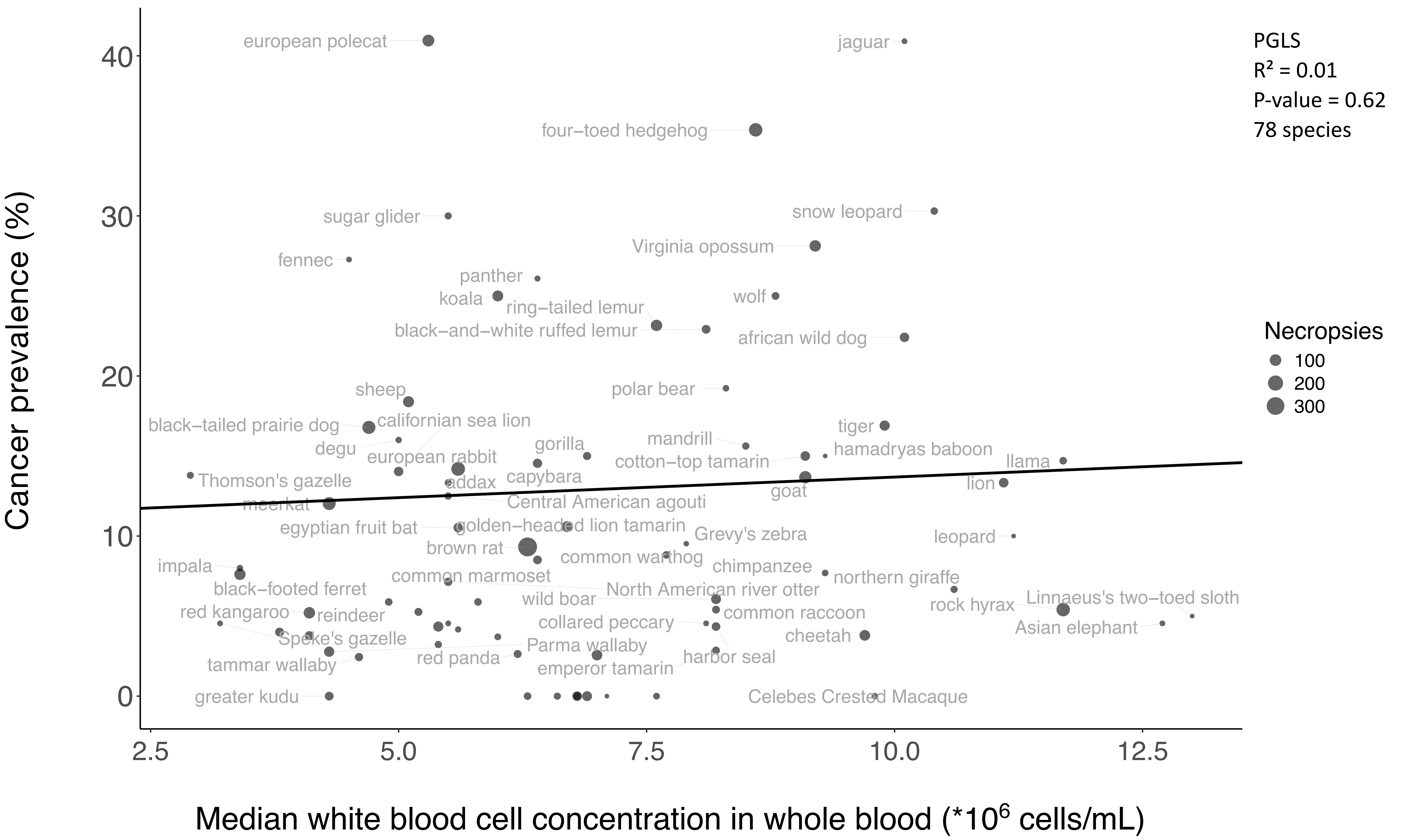
PGLS
R² = 0.01
P-value = 0.62
78 species
Necropsies
100
200
300
Cancer prevalence (%)
Median white blood cell concentration in whole blood (*10^6 cells/mL)
european polecat
jaguar
four–toed hedgehog
sugar glider
snow leopard
Virginia opossum
fennec
panther
koala
ring–tailed lemur
wolf
black–and–white ruffed lemur
african wild dog
sheep
polar bear
californian sea lion
black–tailed prairie dog
tiger
mandrill
degu
gorilla
hamadryas baboon
european rabbit
cotton–top tamarin
llama
capybara
Thomson's gazelle
addax
lion
goat
meerkat
Central American agouti
egyptian fruit bat
golden–headed lion tamarin
Grevy's zebra
leopard
brown rat
common warthog
impala
chimpanzee
northern giraffe
common marmoset
black–footed ferret
North American river otter
wild boar
rock hyrax
Linnaeus's two–toed sloth
red kangaroo
reindeer
common raccoon
collared peccary
Asian elephant
Speke's gazelle
cheetah
Parma wallaby
red panda
harbor seal
tammar wallaby
emperor tamarin
greater kudu
Celebes Crested Macaque

Supp.Fig.12B

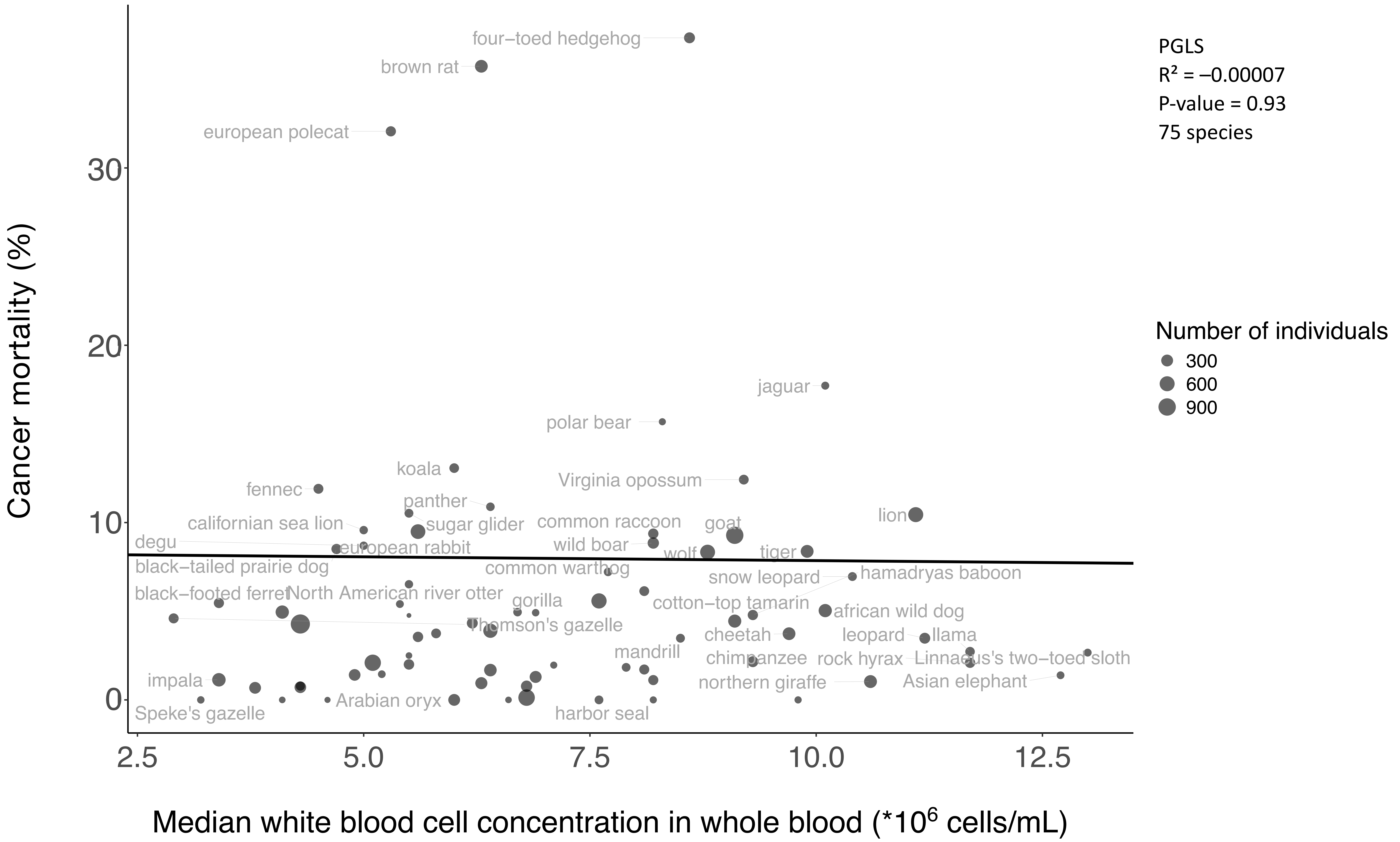

four−toed hedgehog
brown rat
european polecat
jaguar
polar bear
koala
Virginia opossum
fennec
panther
lion
californian sea lion
sugar glider
common raccoon
goat
degu
wild boar
european rabbit
wolf
tiger
black−tailed prairie dog
common warthog
snow leopard
hamadryas baboon
black−footed ferret
North American river otter
gorilla
cotton−top tamarin
african wild dog
Thomson's gazelle
cheetah
leopard
llama
mandrill
chimpanzee
rock hyrax
Linnaeus's two−toed sloth
impala
northern giraffe
Asian elephant
Arabian oryx
Speke's gazelle
harbor seal
Cancer mortality (%)
0
10
20
30
2.5
5.0
7.5
10.0
12.5
Median white blood cell concentration in whole blood (*10^6 cells/mL)
PGLS
R² = −0.00007
P-value = 0.93
75 species
Number of individuals
300
600
900

Supp.Fig.13A

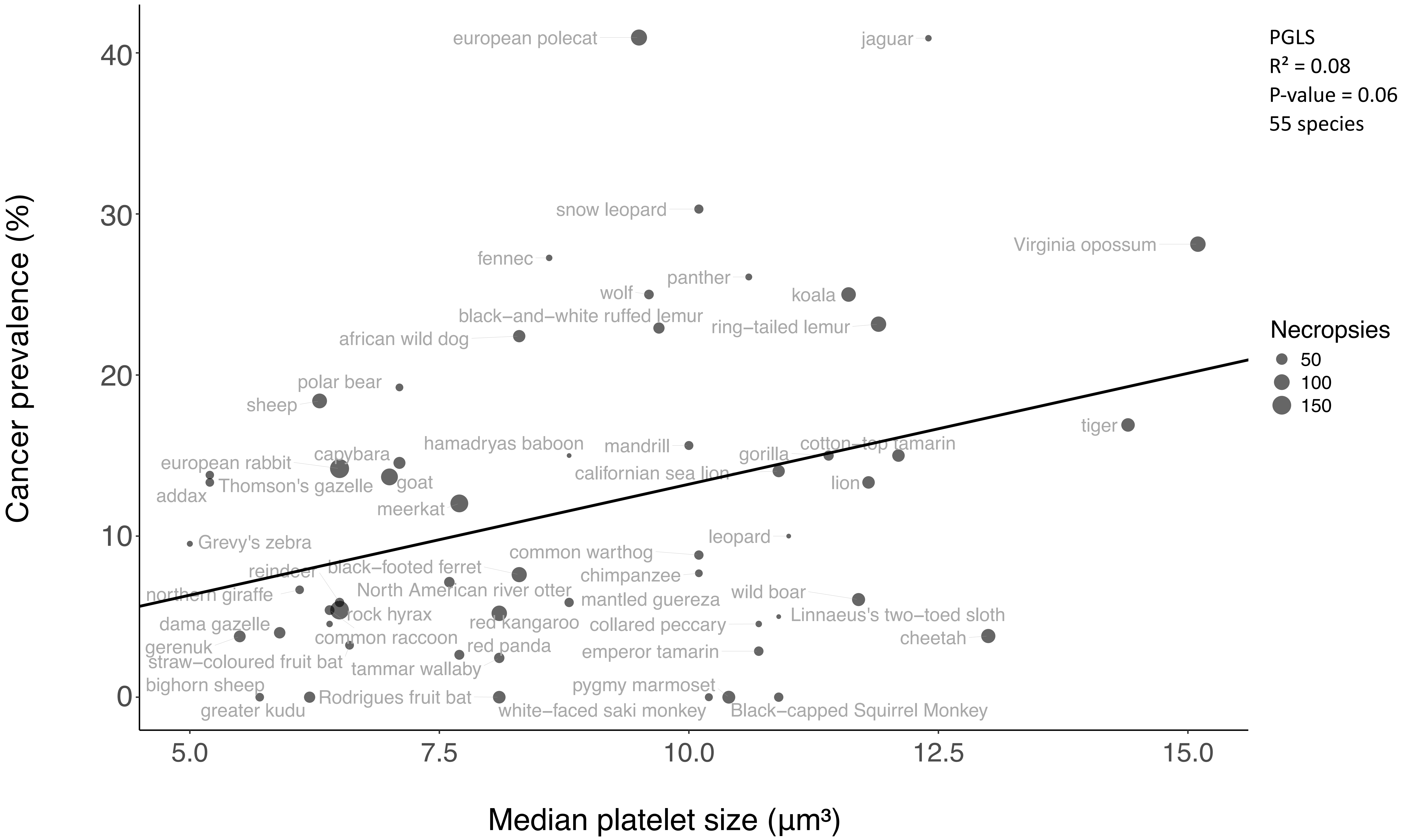
PGLS
R² = 0.08
P-value = 0.06
55 species
Necropsies
50
100
150
Cancer prevalence (%)
Median platelet size (µm³)
0
10
20
30
40
5.0
7.5
10.0
12.5
15.0
european polecat
jaguar
snow leopard
Virginia opossum
fennec
panther
wolf
koala
black−and−white ruffed lemur
ring−tailed lemur
african wild dog
polar bear
sheep
tiger
hamadryas baboon
mandrill
cotton−top tamarin
gorilla
european rabbit
capybara
Thomson's gazelle
goat
californian sea lion
lion
addax
meerkat
Grevy's zebra
leopard
common warthog
reindeer
black−footed ferret
chimpanzee
northern giraffe
North American river otter
wild boar
mantled guereza
rock hyrax
Linnaeus's two−toed sloth
dama gazelle
red kangaroo
collared peccary
common raccoon
cheetah
gerenuk
red panda
emperor tamarin
straw−coloured fruit bat
tammar wallaby
bighorn sheep
pygmy marmoset
Rodrigues fruit bat
greater kudu
white−faced saki monkey
Black−capped Squirrel Monkey

Supp.Fig.13B

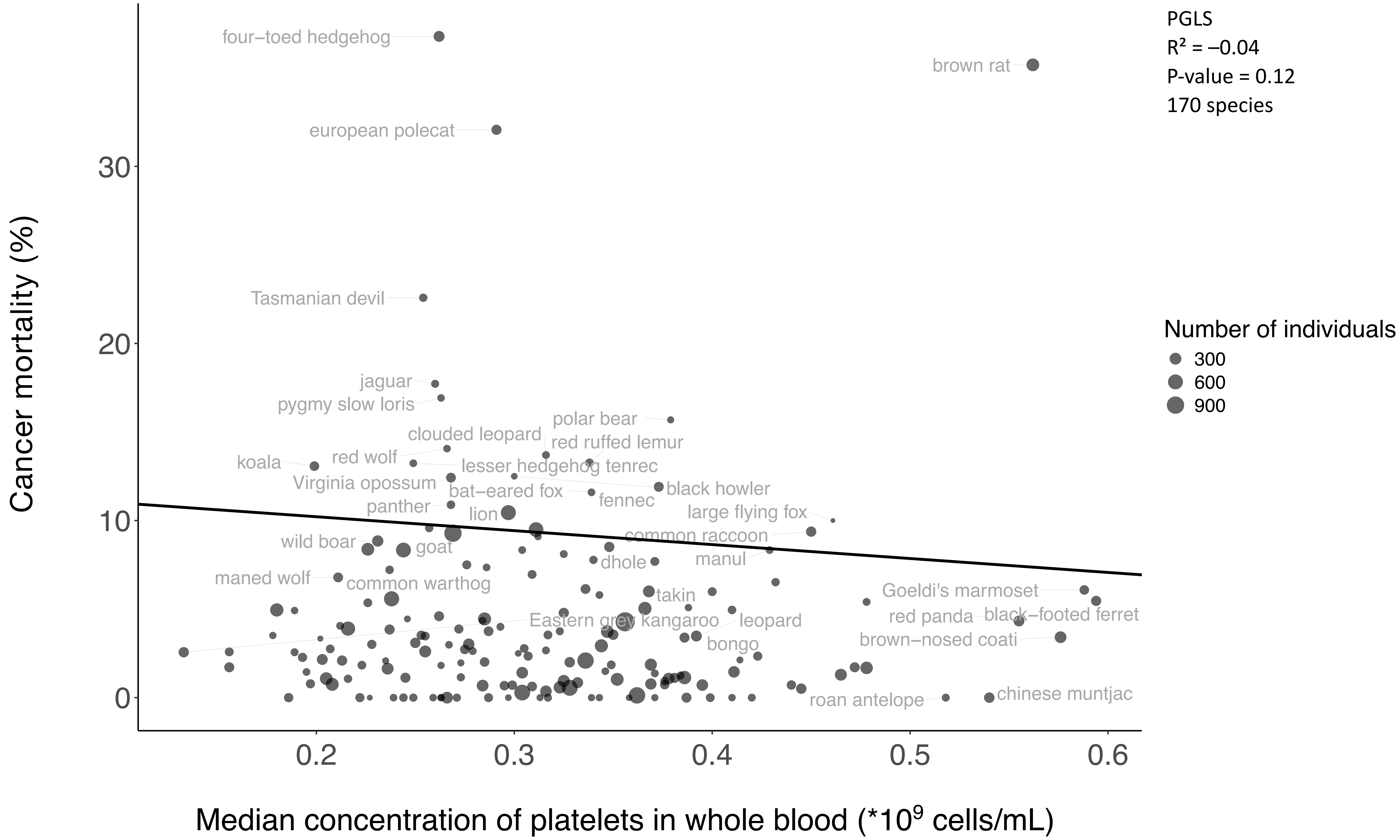

PGLS
R² = –0.04
P-value = 0.12
170 species
Number of individuals
300
600
900
Cancer mortality (%)
Median concentration of platelets in whole blood (*10^9 cells/mL)
0
10
20
30
0.2
0.3
0.4
0.5
0.6
four–toed hedgehog
brown rat
european polecat
Tasmanian devil
jaguar
pygmy slow loris
polar bear
clouded leopard
red ruffed lemur
red wolf
koala
lesser hedgehog tenrec
Virginia opossum
black howler
bat–eared fox
fennec
panther
lion
large flying fox
common raccoon
wild boar
goat
manul
dhole
maned wolf
common warthog
Goeldi's marmoset
takin
red panda
black–footed ferret
Eastern grey kangaroo
leopard
brown–nosed coati
bongo
roan antelope
chinese muntjac

Supp.Fig.13C

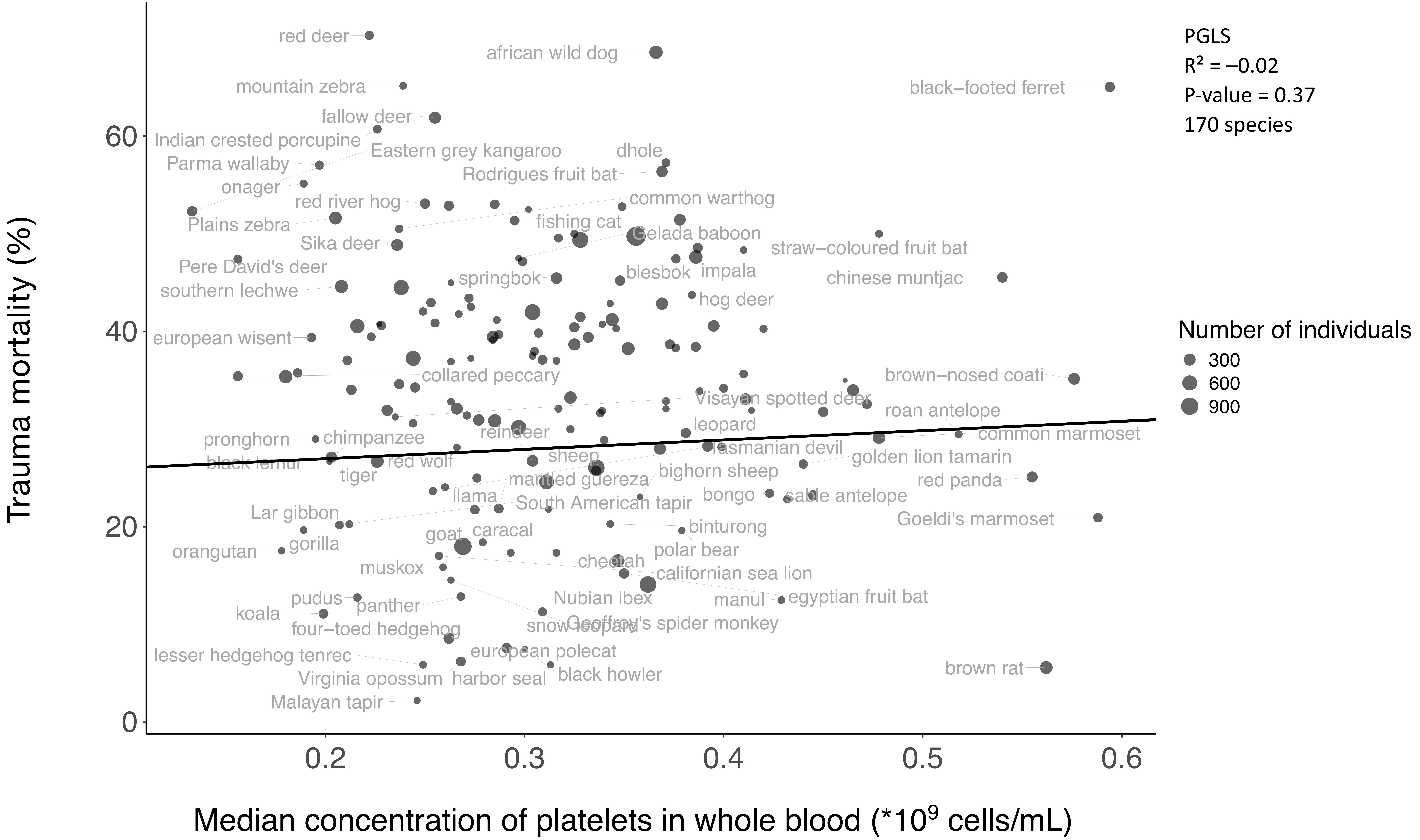

PGLS
R² = −0.02
P-value = 0.37
170 species
Number of individuals
300
600
900
Trauma mortality (%)
Median concentration of platelets in whole blood (*10^9 cells/mL)
0
20
40
60
0.2
0.3
0.4
0.5
0.6
red deer
african wild dog
mountain zebra
black−footed ferret
fallow deer
Indian crested porcupine
Eastern grey kangaroo
dhole
Parma wallaby
Rodrigues fruit bat
onager
red river hog
common warthog
Plains zebra
fishing cat
Sika deer
Gelada baboon
straw−coloured fruit bat
Pere David's deer
impala
springbok
blesbok
chinese muntjac
southern lechwe
hog deer
european wisent
collared peccary
brown−nosed coati
Visayan spotted deer
roan antelope
leopard
reindeer
common marmoset
pronghorn
chimpanzee
tasmanian devil
black lemur
red wolf
sheep
golden lion tamarin
tiger
mantled guereza
bighorn sheep
red panda
llama
bongo
sable antelope
South American tapir
Lar gibbon
binturong
Goeldi's marmoset
goat
caracal
polar bear
orangutan
gorilla
cheetah
muskox
californian sea lion
pudus
Nubian ibex
manul
egyptian fruit bat
panther
koala
snow leopard
Geoffroy's spider monkey
four−toed hedgehog
european polecat
lesser hedgehog tenrec
brown rat
Virginia opossum
harbor seal
black howler
Malayan tapir

Supp.Fig.13D

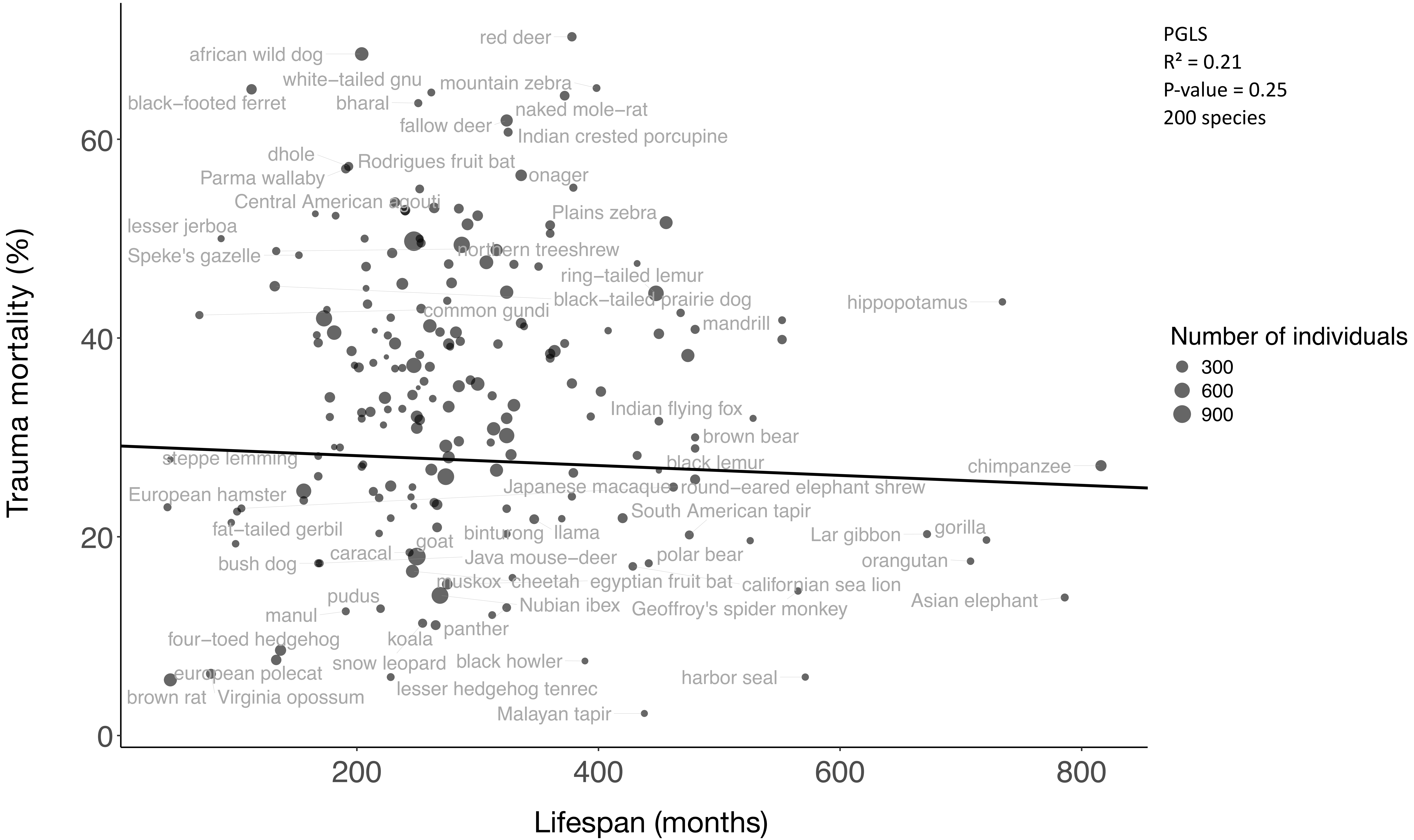

PGLS
R² = 0.21
P-value = 0.25
200 species
Number of individuals
300
600
900
Trauma mortality (%)
Lifespan (months)
0
20
40
60
200
400
600
800
red deer
african wild dog
white−tailed gnu
mountain zebra
black−footed ferret
bharal
naked mole−rat
fallow deer
Indian crested porcupine
dhole
Rodrigues fruit bat
Parma wallaby
onager
Central American agouti
Plains zebra
lesser jerboa
northern treeshrew
Speke's gazelle
ring−tailed lemur
black−tailed prairie dog
hippopotamus
common gundi
mandrill
Indian flying fox
brown bear
steppe lemming
black lemur
chimpanzee
Japanese macaque
round−eared elephant shrew
European hamster
South American tapir
fat−tailed gerbil
binturong
llama
Lar gibbon
gorilla
goat
caracal
Java mouse−deer
polar bear
orangutan
bush dog
muskox
cheetah
egyptian fruit bat
californian sea lion
pudus
Nubian ibex
Geoffroy's spider monkey
Asian elephant
manul
panther
four−toed hedgehog
koala
black howler
snow leopard
european polecat
harbor seal
brown rat
Virginia opossum
lesser hedgehog tenrec
Malayan tapir

Supp.Fig.13E

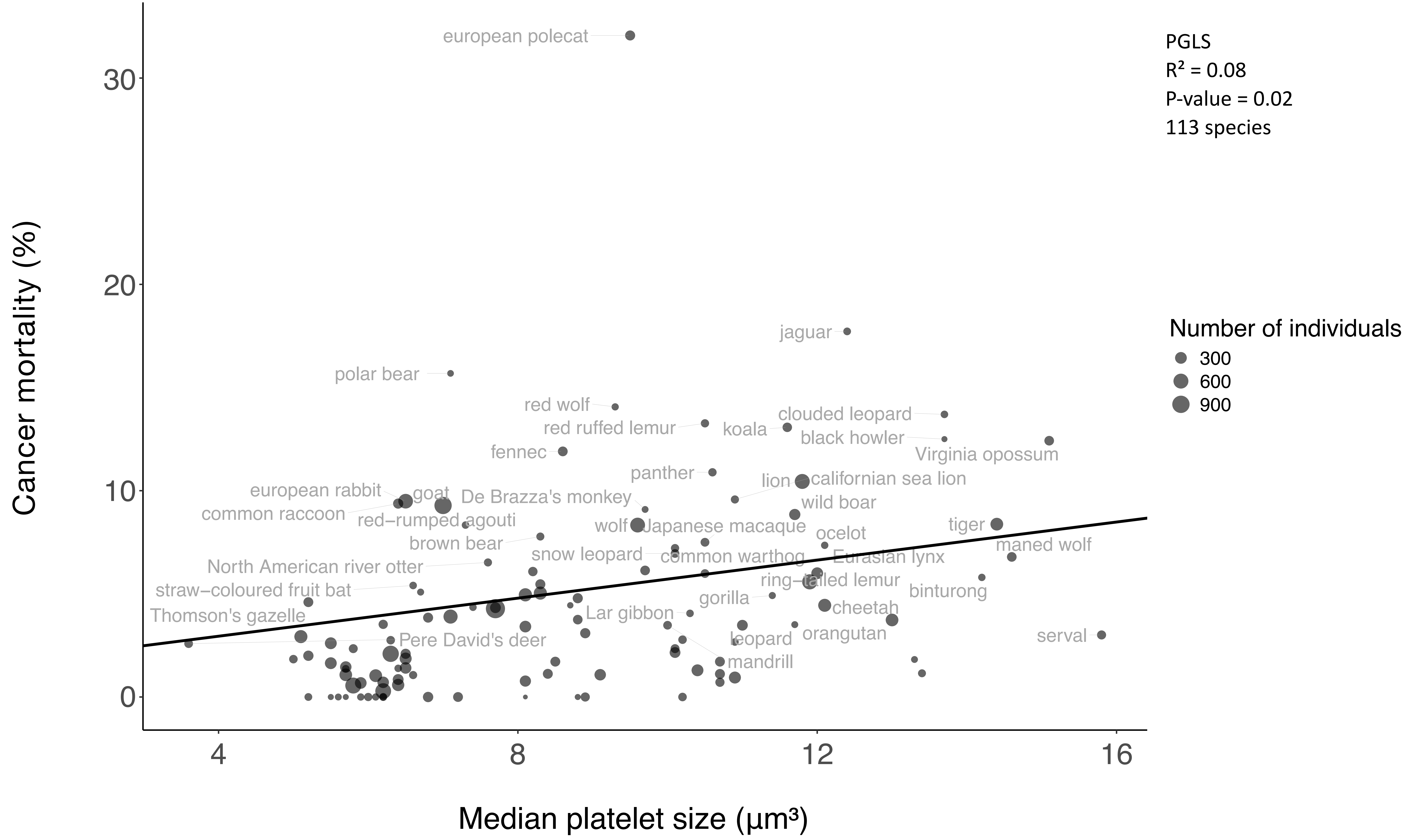

Cancer mortality (%)
Median platelet size (µm³)
0
10
20
30
4
8
12
16
PGLS
R² = 0.08
P-value = 0.02
113 species
Number of individuals
300
600
900
european polecat
jaguar
polar bear
red wolf
red ruffed lemur
koala
clouded leopard
black howler
Virginia opossum
fennec
panther
lion
californian sea lion
european rabbit
goat
De Brazza's monkey
common raccoon
red−rumped agouti
wild boar
wolf
Japanese macaque
tiger
ocelot
brown bear
maned wolf
snow leopard
common warthog
Eurasian lynx
North American river otter
ring−tailed lemur
straw−coloured fruit bat
binturong
gorilla
cheetah
Lar gibbon
Thomson's gazelle
orangutan
serval
Pere David's deer
leopard
mandrill

Supp.Fig.14A

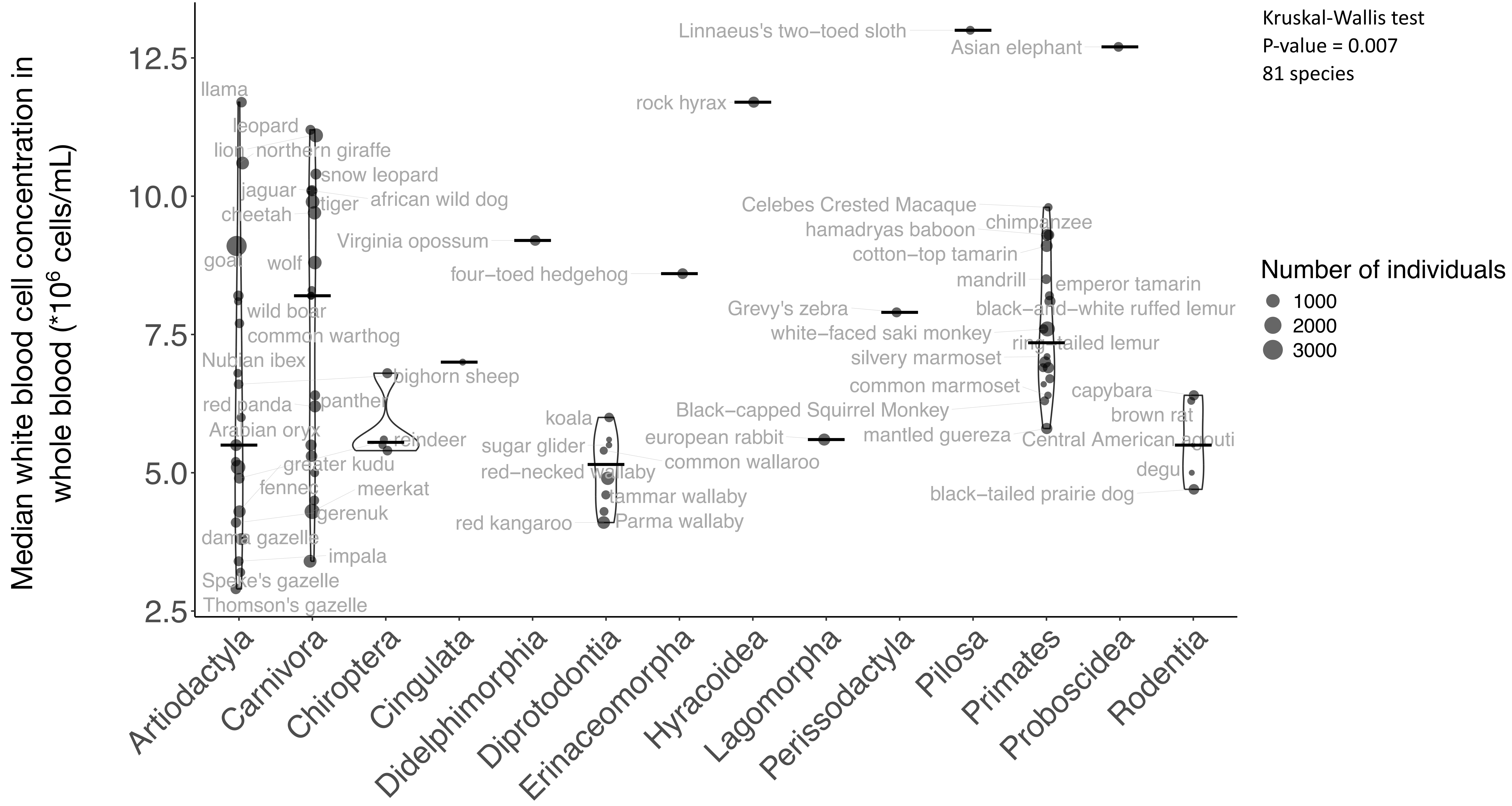
Median white blood cell concentration in whole blood (*10^6 cells/mL)
2.5
5.0
7.5
10.0
12.5
Artiodactyla
Carnivora
Chiroptera
Cingulata
Didelphimorphia
Diprotodontia
Erinaceomorpha
Hyracoidea
Lagomorpha
Perissodactyla
Pilosa
Primates
Proboscidea
Rodentia
Kruskal-Wallis test
P-value = 0.007
81 species
Number of individuals
1000
2000
3000
llama
leopard
lion
northern giraffe
snow leopard
jaguar
african wild dog
tiger
cheetah
goat
wolf
wild boar
common warthog
Nubian ibex
bighorn sheep
red panda
panther
Arabian oryx
reindeer
greater kudu
fennec
meerkat
gerenuk
dama gazelle
impala
Speke's gazelle
Thomson's gazelle
Virginia opossum
four-toed hedgehog
koala
sugar glider
red-necked wallaby
common wallaroo
tammar wallaby
red kangaroo
Parma wallaby
Linnaeus's two-toed sloth
Asian elephant
rock hyrax
european rabbit
Grevy's zebra
Celebes Crested Macaque
chimpanzee
hamadryas baboon
cotton-top tamarin
mandrill
emperor tamarin
black-and-white ruffed lemur
white-faced saki monkey
ring-tailed lemur
silvery marmoset
common marmoset
capybara
Black-capped Squirrel Monkey
brown rat
mantled guereza
Central American agouti
degu
black-tailed prairie dog

Supp.Fig.14B

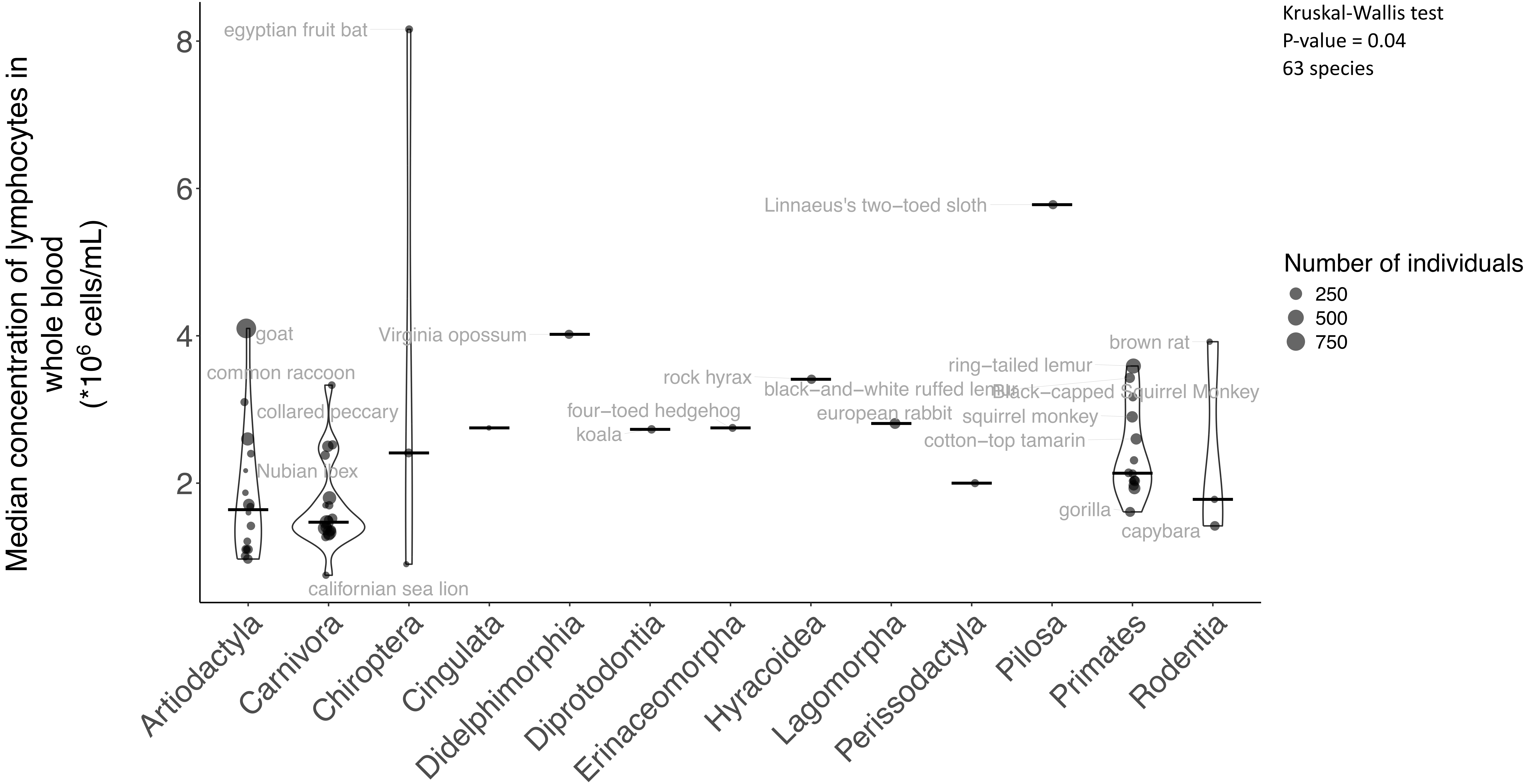

Kruskal-Wallis test
P-value = 0.04
63 species
Number of individuals
250
500
750
Median concentration of lymphocytes in whole blood (*10^6 cells/mL)
8
6
4
2
egyptian fruit bat
Linnaeus's two−toed sloth
goat
Virginia opossum
brown rat
common raccoon
ring−tailed lemur
rock hyrax
black−and−white ruffed lemur
Black−capped Squirrel Monkey
collared peccary
four−toed hedgehog
european rabbit
squirrel monkey
koala
cotton−top tamarin
Nubian ibex
gorilla
capybara
californian sea lion
Artiodactyla
Carnivora
Chiroptera
Cingulata
Didelphimorphia
Diprotodontia
Erinaceomorpha
Hyracoidea
Lagomorpha
Perissodactyla
Pilosa
Primates
Rodentia

Supp.Fig.14C

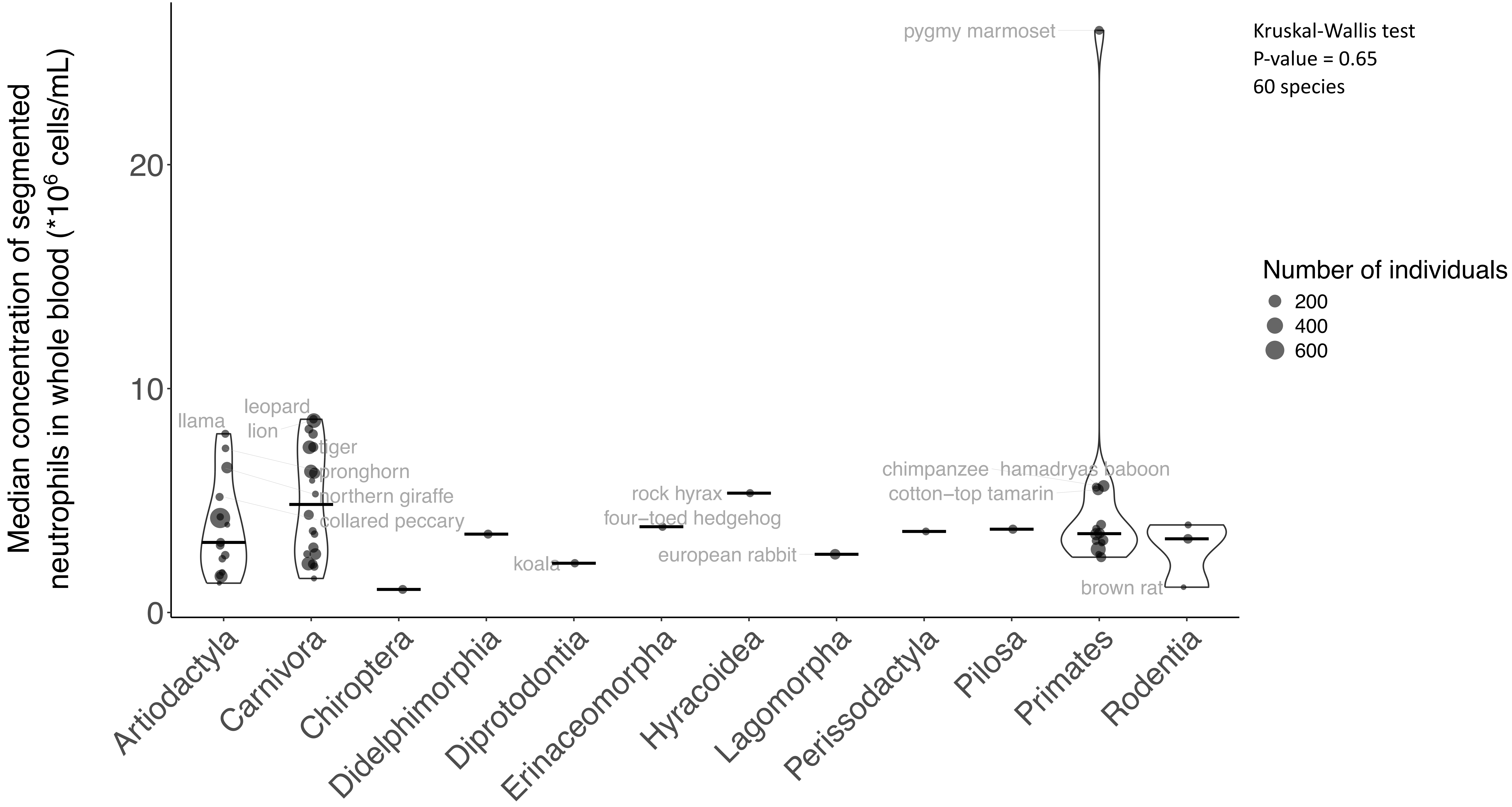

Kruskal-Wallis test
P-value = 0.65
60 species
Number of individuals
200
400
600
Median concentration of segmented neutrophils in whole blood (*10^6 cells/mL)
0
10
20
pygmy marmoset
llama
leopard
lion
tiger
pronghorn
northern giraffe
collared peccary
koala
rock hyrax
four−toed hedgehog
european rabbit
chimpanzee
hamadryas baboon
cotton−top tamarin
brown rat
Artiodactyla
Carnivora
Chiroptera
Didelphimorphia
Diprotodontia
Erinaceomorpha
Hyracoidea
Lagomorpha
Perissodactyla
Pilosa
Primates
Rodentia

Supp.Fig.14D

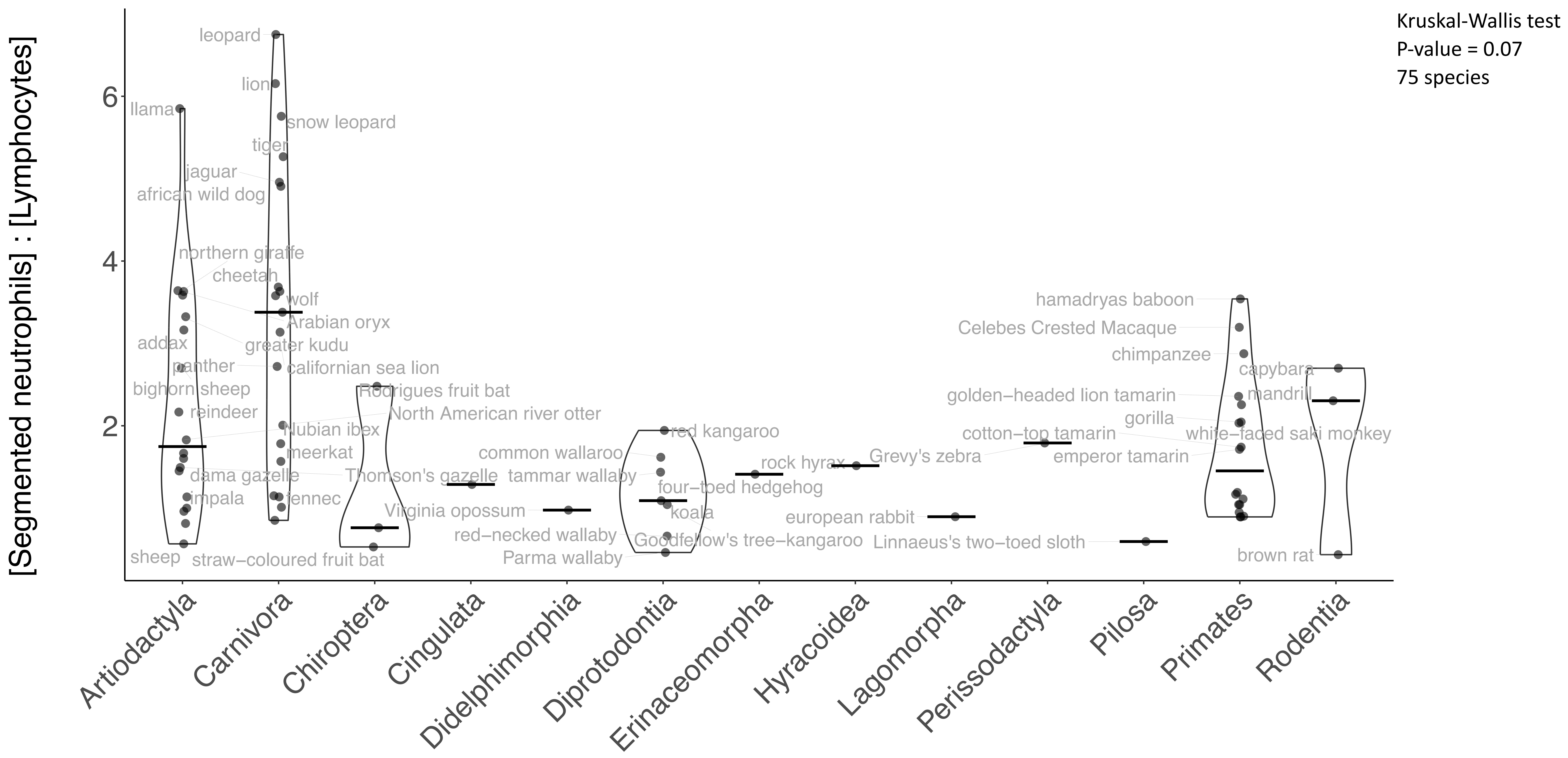
Kruskal-Wallis test
P-value = 0.07
75 species
[Segmented neutrophils] : [Lymphocytes]
0
2
4
6
Artiodactyla
Carnivora
Chiroptera
Cingulata
Didelphimorphia
Diprotodontia
Erinaceomorpha
Hyracoidea
Lagomorpha
Perissodactyla
Pilosa
Primates
Rodentia
leopard
lion
llama
snow leopard
tiger
jaguar
african wild dog
northern giraffe
cheetah
wolf
Arabian oryx
addax
greater kudu
panther
californian sea lion
bighorn sheep
Rodrigues fruit bat
reindeer
North American river otter
Nubian ibex
meerkat
dama gazelle
Thomson's gazelle
impala
fennec
sheep
straw−coloured fruit bat
common wallaroo
tammar wallaby
Virginia opossum
red−necked wallaby
Parma wallaby
red kangaroo
four−toed hedgehog
koala
Goodfellow's tree−kangaroo
rock hyrax
Grevy's zebra
european rabbit
Linnaeus's two−toed sloth
cotton−top tamarin
hamadryas baboon
Celebes Crested Macaque
chimpanzee
capybara
golden−headed lion tamarin
mandrill
gorilla
white−faced saki monkey
emperor tamarin
brown rat

Supp.Fig.14E

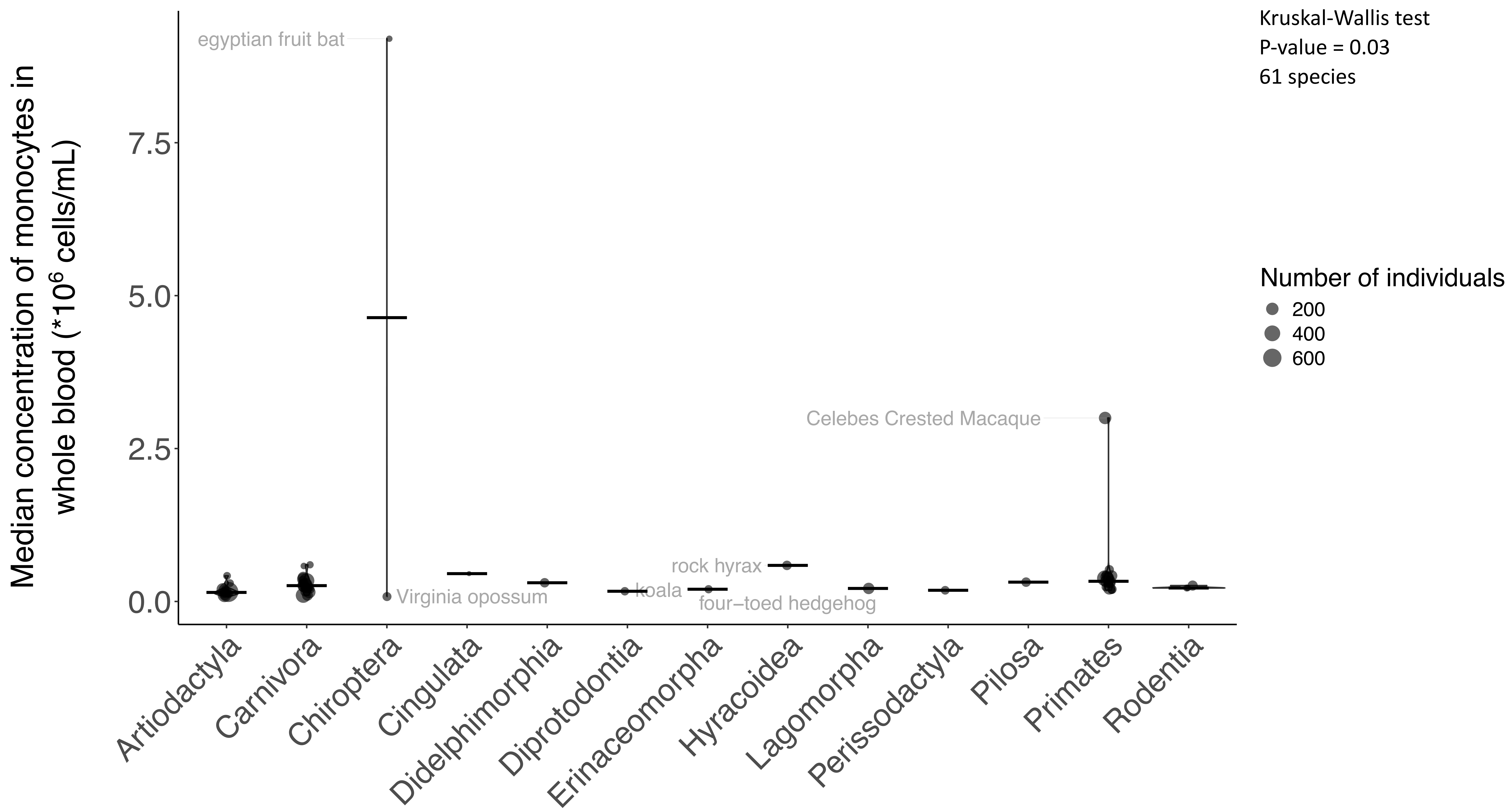

Kruskal-Wallis test
P-value = 0.03
61 species
Number of individuals
200
400
600
Median concentration of monocytes in whole blood (*10^6 cells/mL)
7.5
5.0
2.5
0.0
egyptian fruit bat
Virginia opossum
koala
rock hyrax
four−toed hedgehog
Celebes Crested Macaque
Artiodactyla
Carnivora
Chiroptera
Cingulata
Didelphimorphia
Diprotodontia
Erinaceomorpha
Hyracoidea
Lagomorpha
Perissodactyla
Pilosa
Primates
Rodentia

Supp.Fig.14F

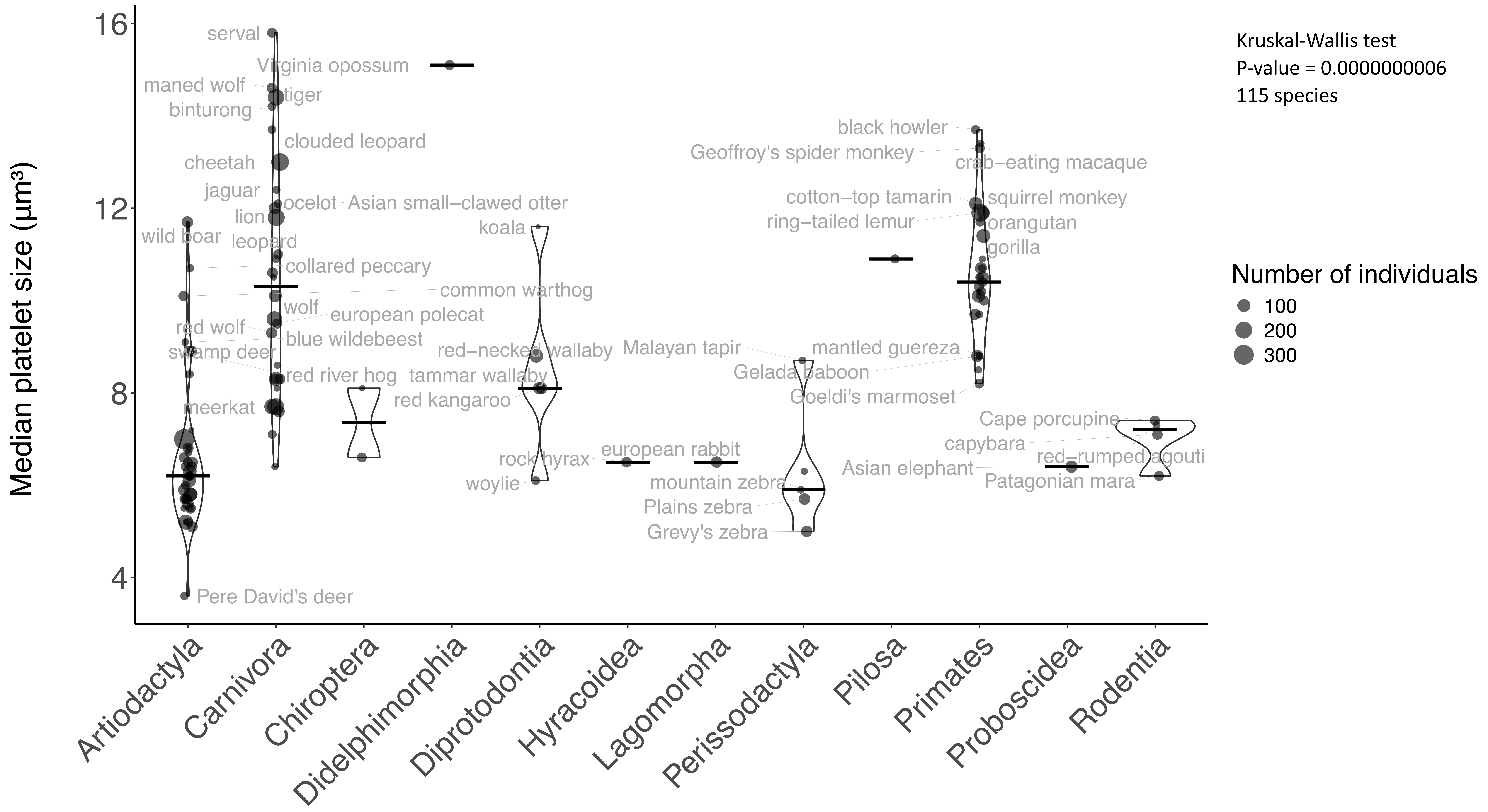
Median platelet size (µm³)
16
12
8
4
Artiodactyla
Carnivora
Chiroptera
Didelphimorphia
Diprotodontia
Hyracoidea
Lagomorpha
Perissodactyla
Pilosa
Primates
Proboscidea
Rodentia
serval
Virginia opossum
maned wolf
tiger
binturong
clouded leopard
cheetah
jaguar
ocelot
Asian small−clawed otter
lion
wild boar
leopard
koala
collared peccary
common warthog
wolf
european polecat
red wolf
blue wildebeest
swamp deer
red−necked wallaby
red river hog
tammar wallaby
red kangaroo
meerkat
rock hyrax
european rabbit
woylie
mountain zebra
Plains zebra
Grevy's zebra
Malayan tapir
Gelada baboon
Goeldi's marmoset
mantled guereza
black howler
Geoffroy's spider monkey
crab−eating macaque
cotton−top tamarin
squirrel monkey
ring−tailed lemur
orangutan
gorilla
Cape porcupine
capybara
red−rumped agouti
Asian elephant
Patagonian mara
Pere David's deer
Kruskal-Wallis test
P-value = 0.0000000006
115 species
Number of individuals
100
200
300

Supp.Fig.14G

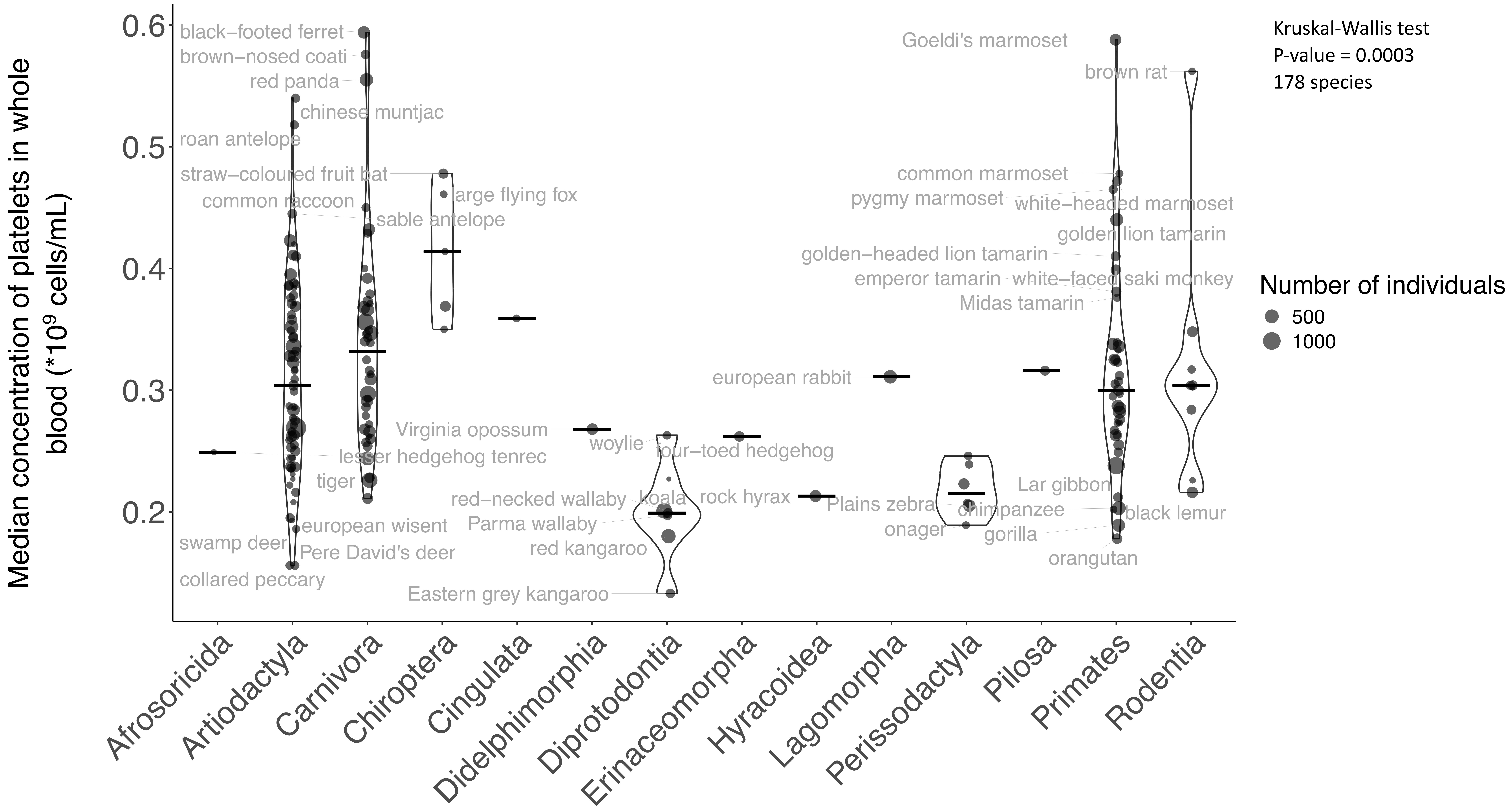

Median concentration of platelets in whole blood (*10^9 cells/mL)
0.6
0.5
0.4
0.3
0.2
Kruskal-Wallis test
P-value = 0.0003
178 species
Number of individuals
500
1000
black−footed ferret
brown−nosed coati
red panda
chinese muntjac
roan antelope
straw−coloured fruit bat
common raccoon
large flying fox
sable antelope
lesser hedgehog tenrec
tiger
european wisent
swamp deer
Pere David's deer
collared peccary
Virginia opossum
woylie
four−toed hedgehog
red−necked wallaby
koala
Parma wallaby
red kangaroo
Eastern grey kangaroo
rock hyrax
european rabbit
Plains zebra
onager
chimpanzee
gorilla
Lar gibbon
black lemur
orangutan
Goeldi's marmoset
brown rat
common marmoset
pygmy marmoset
white−headed marmoset
golden lion tamarin
golden−headed lion tamarin
emperor tamarin
white−faced saki monkey
Midas tamarin
Afrosoricida
Artiodactyla
Carnivora
Chiroptera
Cingulata
Didelphimorphia
Diprotodontia
Erinaceomorpha
Hyracoidea
Lagomorpha
Perissodactyla
Pilosa
Primates
Rodentia

Supp.Fig.14H

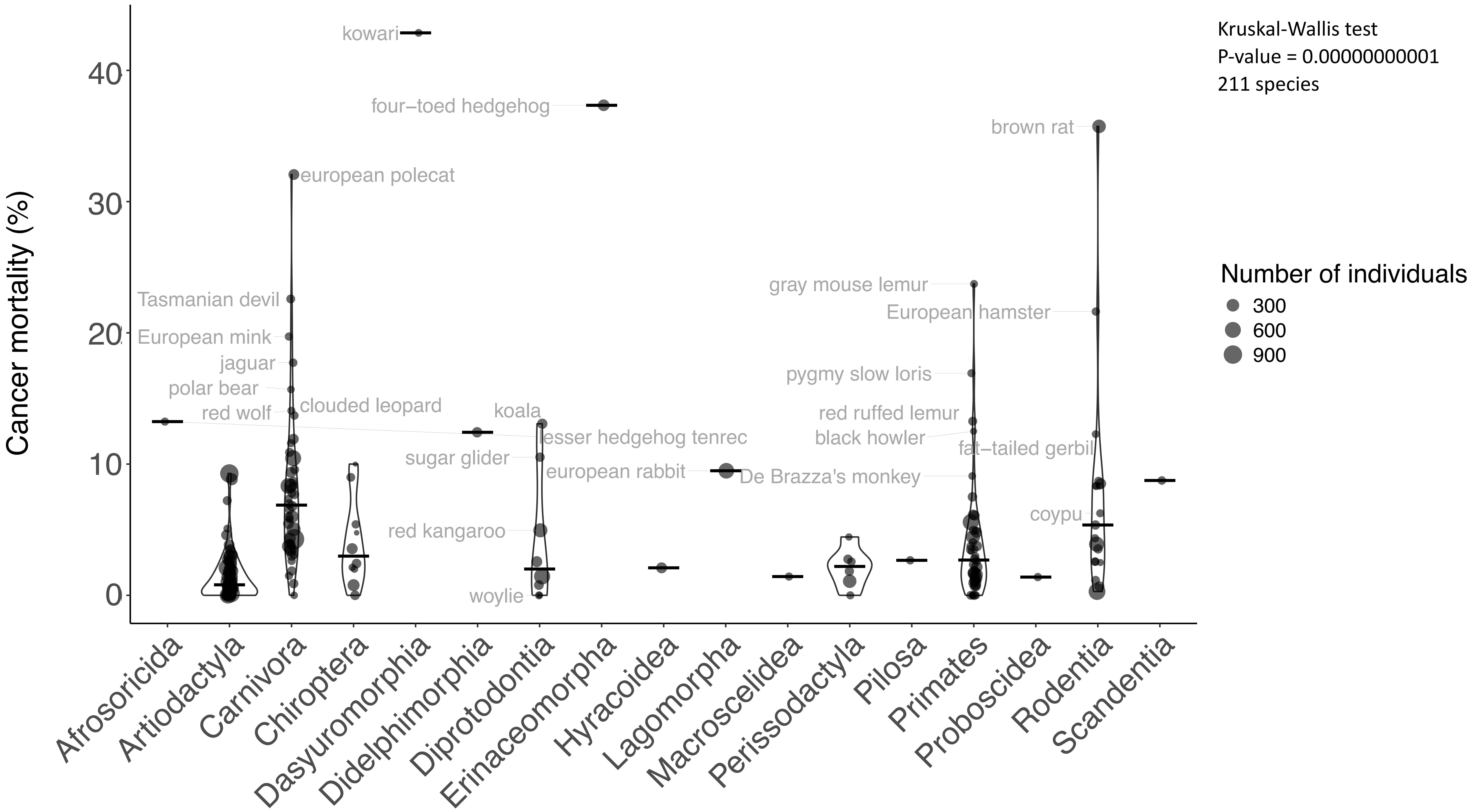

Kruskal-Wallis test
P-value = 0.00000000001
211 species
Number of individuals
300
600
900
Cancer mortality (%)
0
10
20
30
40
kowari
four−toed hedgehog
brown rat
european polecat
Tasmanian devil
European mink
jaguar
polar bear
red wolf
clouded leopard
koala
lesser hedgehog tenrec
sugar glider
european rabbit
red kangaroo
woylie
gray mouse lemur
European hamster
pygmy slow loris
red ruffed lemur
black howler
fat−tailed gerbil
De Brazza's monkey
coypu
Afrosoricida
Artiodactyla
Carnivora
Chiroptera
Dasyuromorphia
Didelphimorphia
Diprotodontia
Erinaceomorpha
Hyracoidea
Lagomorpha
Macroscelidea
Perissodactyla
Pilosa
Primates
Proboscidea
Rodentia
Scandentia

Supp.Fig.14I

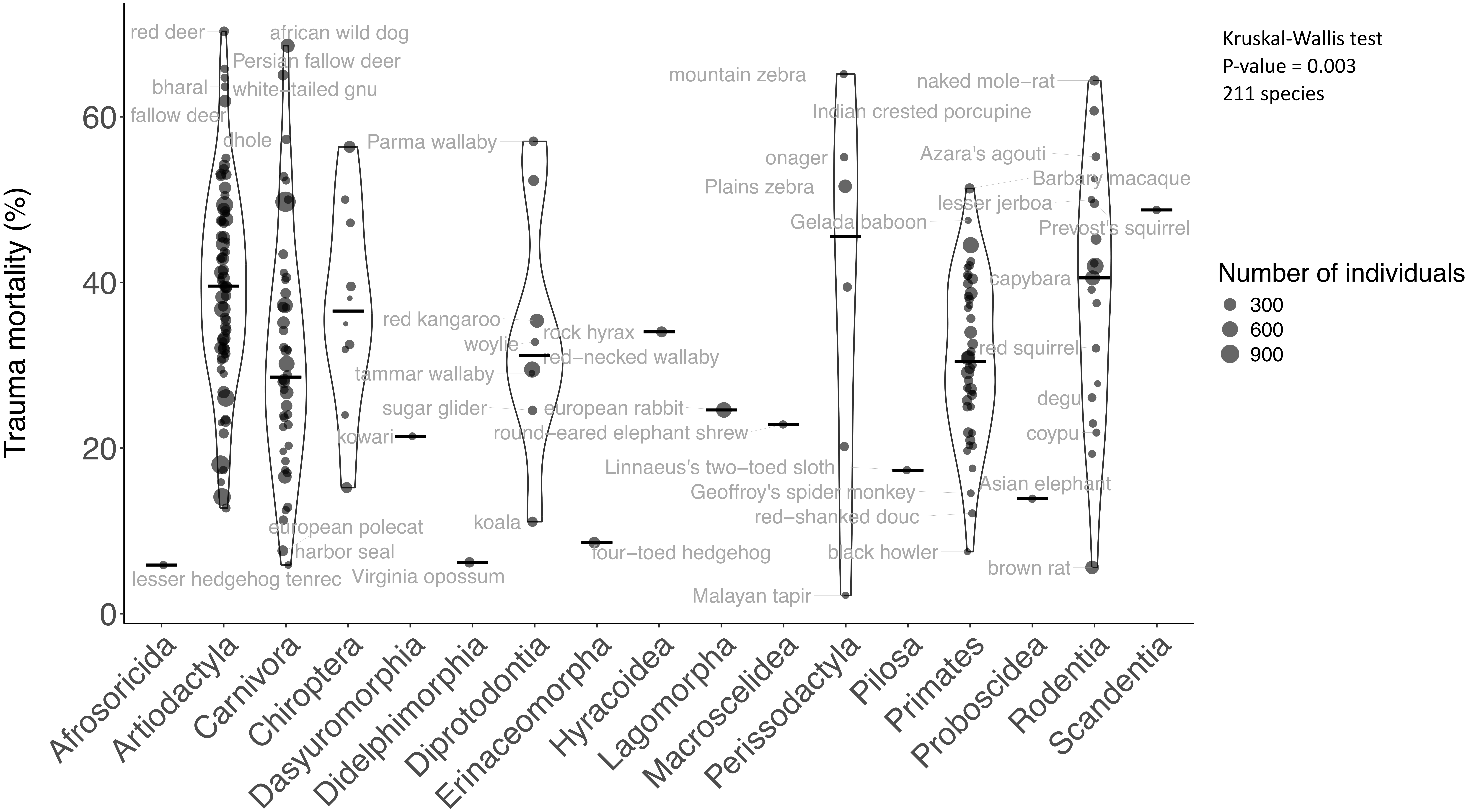

Kruskal-Wallis test
P-value = 0.003
211 species
Number of individuals
300
600
900
Trauma mortality (%)
0
20
40
60
Afrosoricida
Artiodactyla
Carnivora
Chiroptera
Dasyuromorphia
Didelphimorphia
Diprotodontia
Erinaceomorpha
Hyracoidea
Lagomorpha
Macroscelidea
Perissodactyla
Pilosa
Primates
Proboscidea
Rodentia
Scandentia
red deer
african wild dog
Persian fallow deer
bharal
white−tailed gnu
fallow deer
dhole
Parma wallaby
red kangaroo
woylie
rock hyrax
red−necked wallaby
tammar wallaby
sugar glider
european rabbit
round−eared elephant shrew
kowari
koala
european polecat
harbor seal
four−toed hedgehog
lesser hedgehog tenrec
Virginia opossum
mountain zebra
Indian crested porcupine
naked mole−rat
onager
Azara's agouti
Barbary macaque
Plains zebra
lesser jerboa
Gelada baboon
Prevost's squirrel
capybara
red squirrel
degu
coypu
Linnaeus's two−toed sloth
Geoffroy's spider monkey
Asian elephant
red−shanked douc
black howler
brown rat
Malayan tapir

Supp.Fig.15

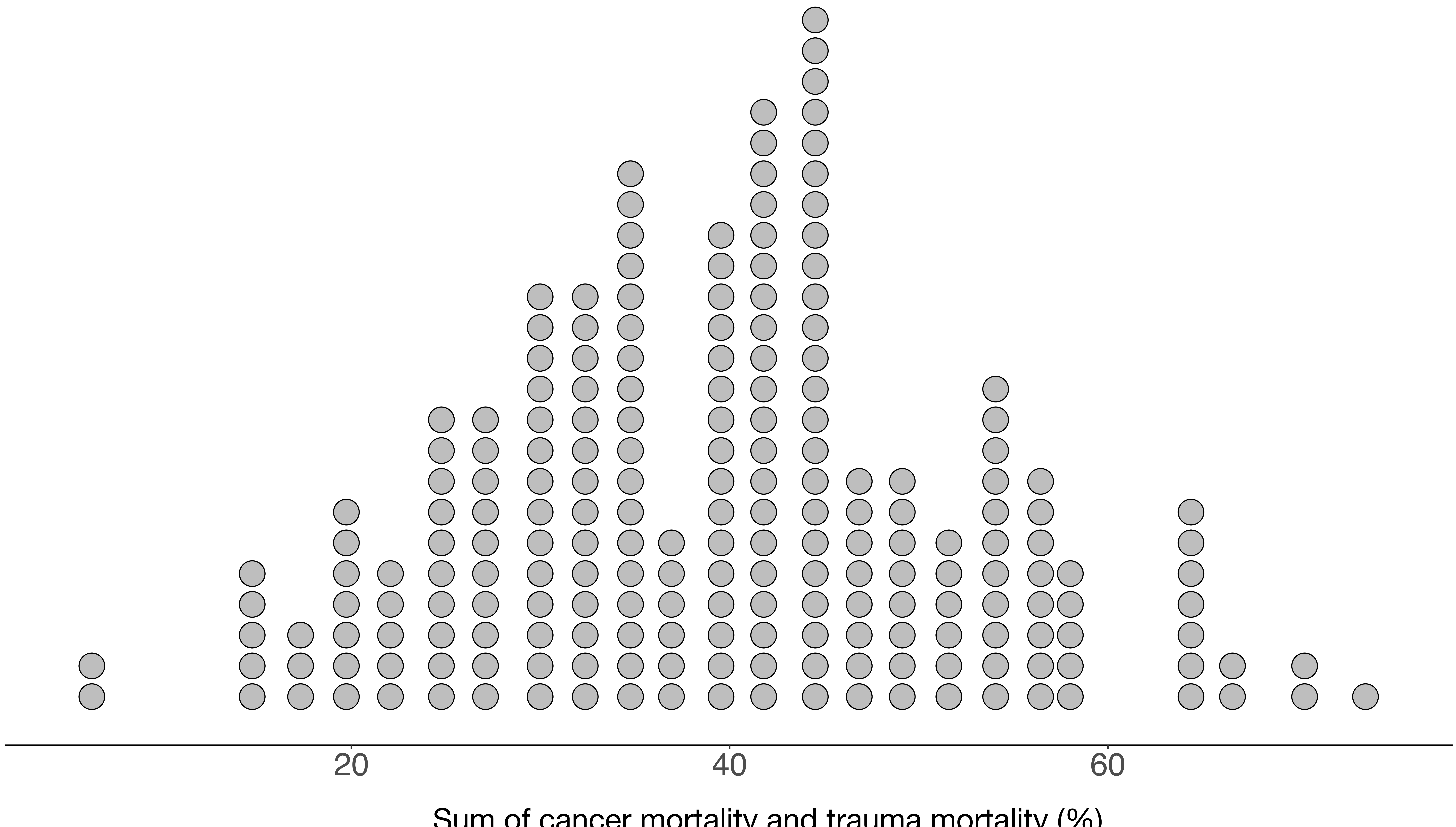

Supp.Fig.16

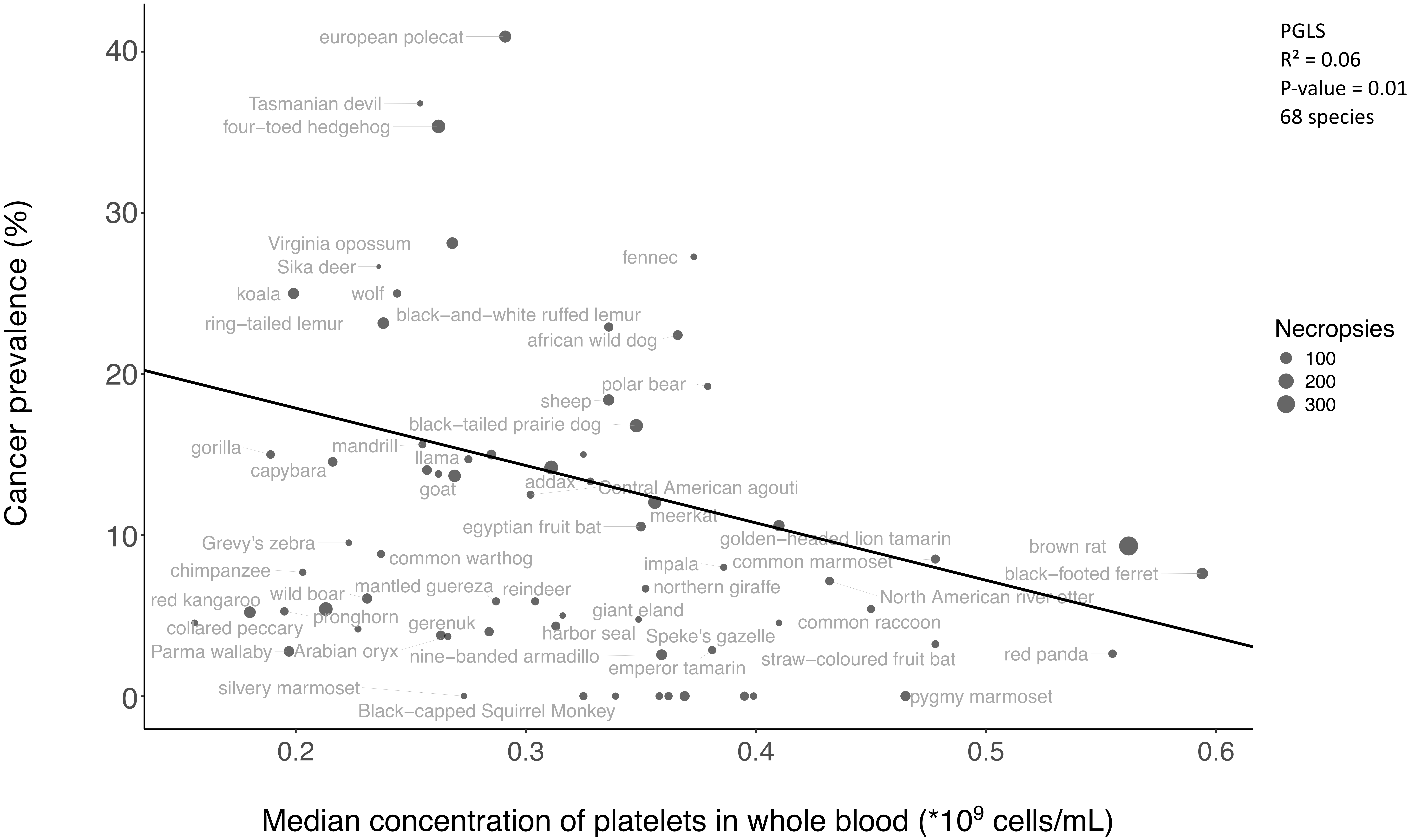

PGLS
R² = 0.06
P-value = 0.01
68 species
Necropsies
100
200
300
Cancer prevalence (%)
Median concentration of platelets in whole blood (*10⁹ cells/mL)
0
10
20
30
40
0.2
0.3
0.4
0.5
0.6
european polecat
Tasmanian devil
four−toed hedgehog
Virginia opossum
Sika deer
fennec
koala
wolf
ring−tailed lemur
black−and−white ruffed lemur
african wild dog
polar bear
sheep
black−tailed prairie dog
gorilla
mandrill
llama
capybara
goat
addax
Central American agouti
meerkat
egyptian fruit bat
golden−headed lion tamarin
brown rat
Grevy's zebra
common warthog
impala
common marmoset
chimpanzee
black−footed ferret
mantled guereza
reindeer
northern giraffe
wild boar
red kangaroo
North American river otter
pronghorn
giant eland
collared peccary
gerenuk
common raccoon
harbor seal
Speke's gazelle
Parma wallaby
Arabian oryx
nine−banded armadillo
straw−coloured fruit bat
red panda
emperor tamarin
silvery marmoset
pygmy marmoset
Black−capped Squirrel Monkey

Supp.Fig.17

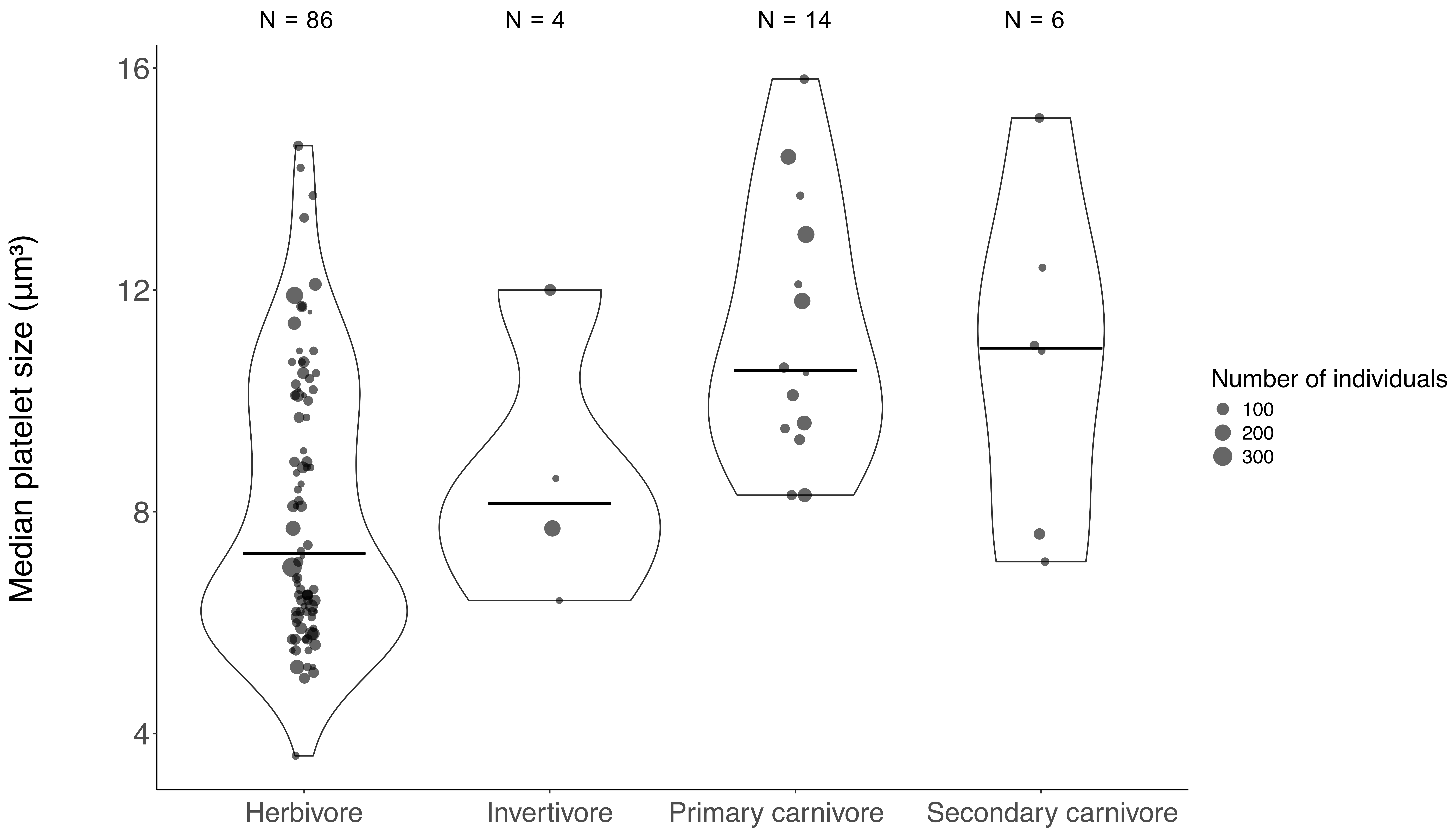

N = 86
N = 4
N = 14
N = 6
16
12
8
4
Median platelet size (µm³)
Herbivore
Invertivore
Primary carnivore
Secondary carnivore
Number of individuals
100
200
300

Supp.Fig.18

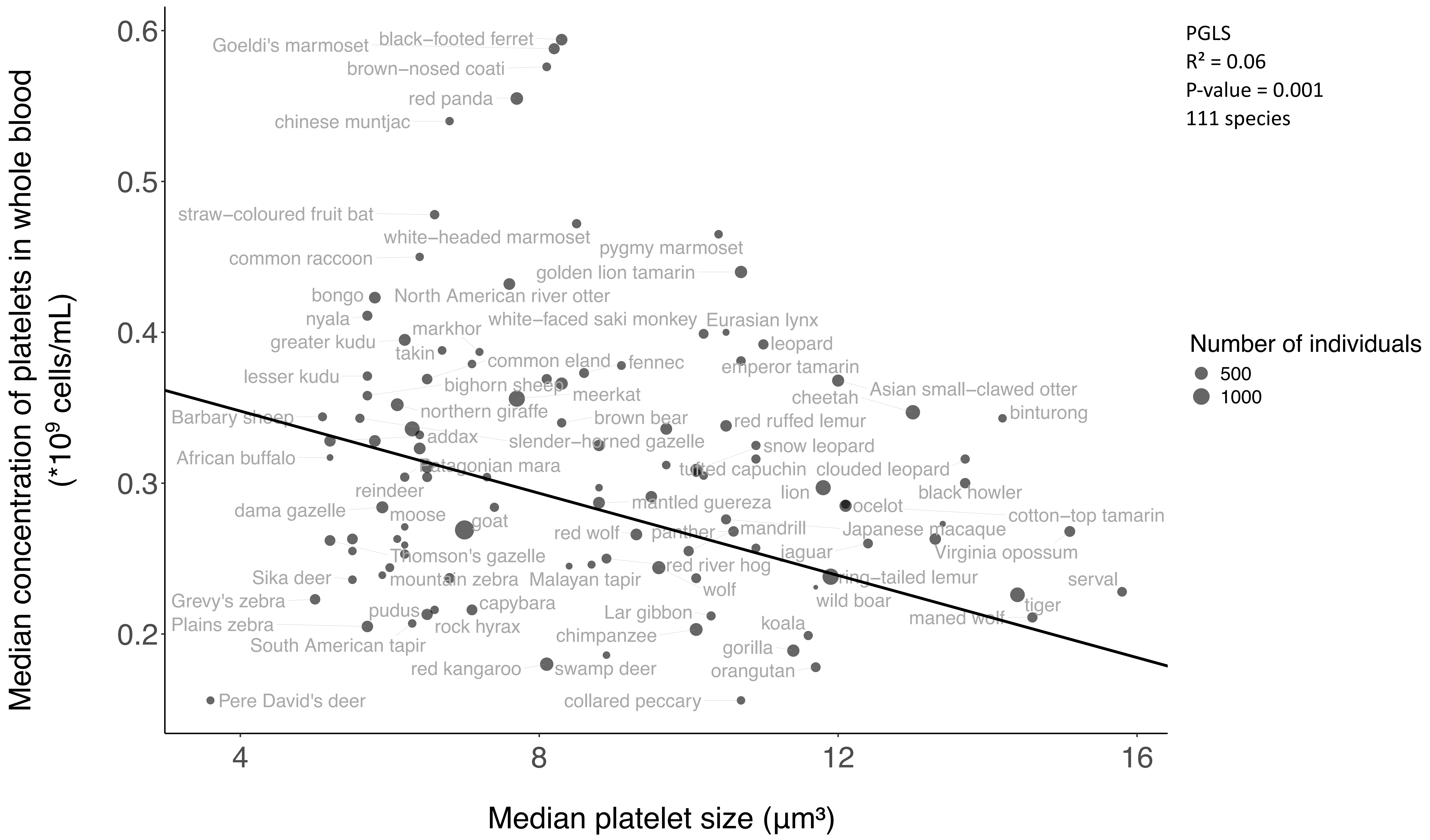
PGLS
R² = 0.06
P-value = 0.001
111 species
Number of individuals
500
1000
Median concentration of platelets in whole blood (*10⁹ cells/mL)
Median platelet size (µm³)
0.6
0.5
0.4
0.3
0.2
4
8
12
16
black–footed ferret
Goeldi's marmoset
brown–nosed coati
red panda
chinese muntjac
straw–coloured fruit bat
white–headed marmoset
pygmy marmoset
common raccoon
golden lion tamarin
bongo
North American river otter
nyala
white–faced saki monkey
Eurasian lynx
markhor
greater kudu
leopard
takin
common eland
fennec
emperor tamarin
lesser kudu
bighorn sheep
meerkat
Asian small–clawed otter
cheetah
northern giraffe
brown bear
binturong
Barbary sheep
red ruffed lemur
addax
slender–horned gazelle
snow leopard
African buffalo
Patagonian mara
tufted capuchin
clouded leopard
reindeer
lion
black howler
mantled guereza
dama gazelle
ocelot
moose
cotton–top tamarin
goat
mandrill
red wolf
panther
Japanese macaque
jaguar
Thomson's gazelle
Virginia opossum
red river hog
Sika deer
mountain zebra
Malayan tapir
ring–tailed lemur
serval
wolf
Grevy's zebra
wild boar
capybara
tiger
pudus
Lar gibbon
rock hyrax
maned wolf
Plains zebra
koala
chimpanzee
South American tapir
gorilla
red kangaroo
swamp deer
orangutan
Pere David's deer
collared peccary

Supp.Fig.19

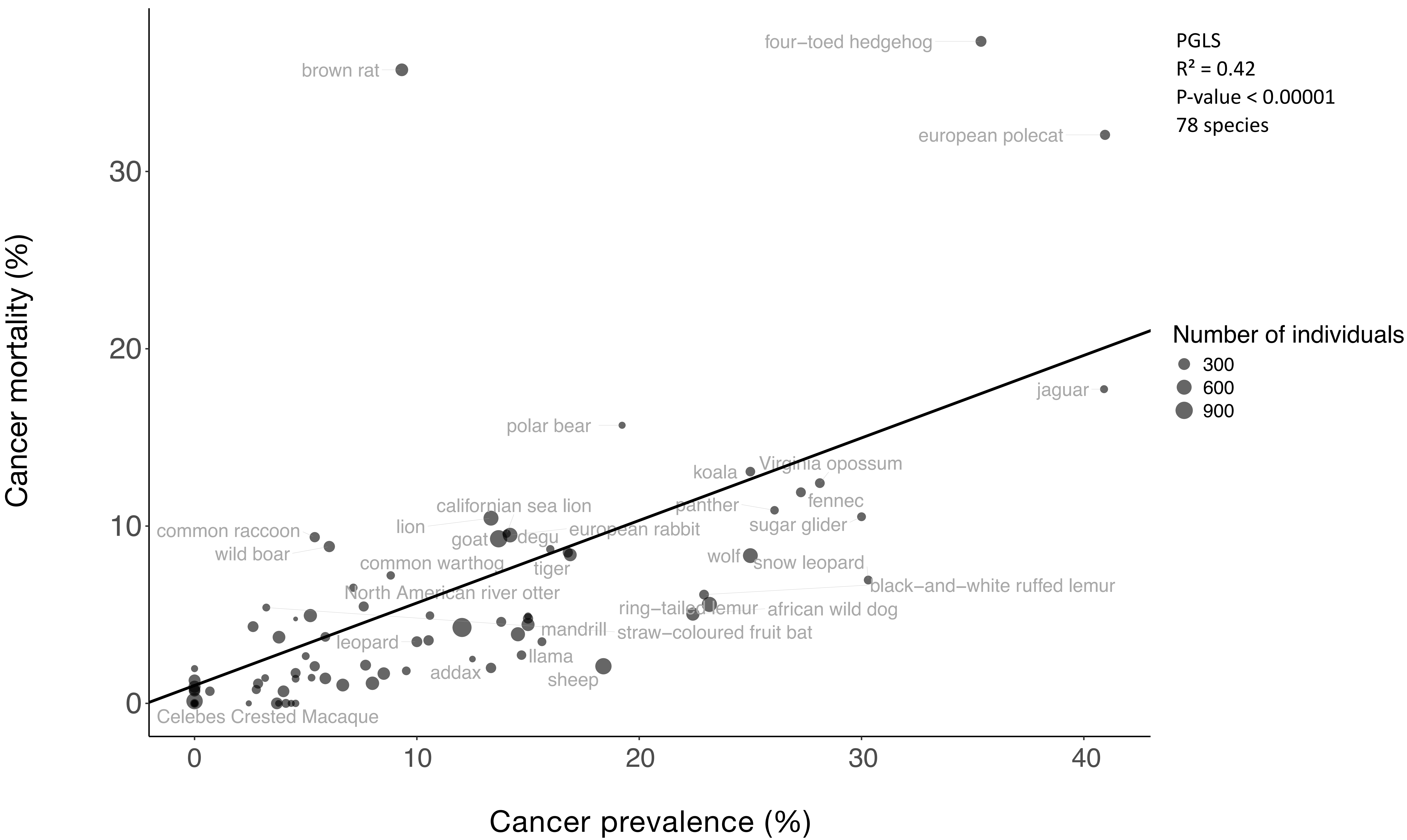

PGLS
$R^2 = 0.42$
P-value < 0.00001
78 species
Number of individuals
300
600
900
Cancer mortality (%)
Cancer prevalence (%)
four−toed hedgehog
brown rat
european polecat
jaguar
polar bear
koala
Virginia opossum
fennec
panther
sugar glider
californian sea lion
lion
goat
degu
european rabbit
common raccoon
wild boar
common warthog
tiger
wolf
snow leopard
black−and−white ruffed lemur
North American river otter
ring−tailed lemur
african wild dog
mandrill
straw−coloured fruit bat
leopard
llama
addax
sheep
Celebes Crested Macaque
